# Habitat-loss-driven predictor coupling limits inference about the independent effects of configuration in additive habitat-amount models: implications for the fragmentation debate

**DOI:** 10.64898/2026.04.12.718042

**Authors:** Juan Andrés Martínez-Lanfranco

**Affiliations:** Department of Biological Sciences, University of Alberta, Canada

**Keywords:** cross-over suppressor attenuation, fragmentation-per-se, geometric separability, habitat amount hypothesis, landscape-scale design, nonlinear coupling, observational inference, structure coefficients, variance partitioning

## Abstract

Additive habitat-amount models are widely used to infer independent configuration effects from observational landscape datasets, yet that inference depends on whether habitat amount and configuration are actually separable in the realised predictor space. Using a global multi-taxa forest dataset assembled from paired continuous and fragmented landscapes, this analysis evaluates that condition directly and shows that it is not met. Habitat amount and configuration remain embedded in a shared habitat-loss gradient with asymmetric nonlinear coupling that standard linear diagnostics do not capture, so near-zero additive fragmentation coefficients do not, by themselves, identify the intended ecological contrast. Under this geometry, the additive specification yields the classic cross-over suppressor signature: fragmentation aligns strongly with the fitted biodiversity gradient yet contributes almost no unique variance once habitat amount is included. When residual coupling is reduced to near zero, fragmentation coefficients shift uniformly negative for both local and landscape-scale diversity, and the same raw additive specification yields negative coefficients in high-cover landscapes, showing that the full-dataset null is geometry-conditional rather than stably ecological. The suppressor structure is absent in beta diversity, indicating that the attenuation is response-specific rather than a universal artefact of the dataset or modelling framework. Because these models are widely used to adjudicate fragmentation-per-se claims from observational data, this issue is a direct challenge to how null configuration coefficients have been interpreted across the fragmentation debate. These results show that a stable ecological-null interpretation is not supported in this dataset — whenever the geometric constraint is reduced, the recoverable direction is uniformly and non-trivially negative. Habitat loss generates configuration change rather than the reverse, embedding asymmetric nonlinear coupling in the attainable predictor space before any landscape is sampled. In empirical landscape datasets, additive control by habitat amount becomes informative about configuration only when the realised predictor geometry has first been shown to support the ecological interpretation being drawn.

## Introduction

Observational landscape ecology routinely uses additive regression to estimate the independent direct effects of habitat amount and configuration on biodiversity (Smith et al. 2009; Fahrig 2017; Fahrig et al. 2019). The procedure is statistically familiar, but its ecological interpretation depends on a stronger condition than model inclusion alone. The predictors must occupy sufficiently distinct regions of the observed predictor space that a partial coefficient can be read as variation along one dimension rather than as a competing allocation from a shared landscape gradient (Koper et al. 2007; Ruffell et al. 2016; Fletcher et al. 2023a). Observational studies are inherently more vulnerable to this problem than randomised experiments because they lack the control over treatment assignment that experiments provide (Shaffer and Johnson 2008). The validity of experiment-style analyses in observational settings depends on assumptions about effective predictor independence that must be empirically verified rather than assumed (McGarigal and Cushman 2002; Shaffer and Johnson 2008; Sagarin and Pauchard 2010).

When that condition fails, additive models still return valid coefficients for the quantities they define, yet those quantities need not correspond to the ecological contrast the analysis was intended to isolate (Freckleton 2002; Ruffell et al. 2016; Lundberg et al. 2021). Additive regression conflates three distinct claims — that predictors are geometrically separable in the realised predictor space, that the separable component has a detectable effect on the response, and that this effect is interpretable as an independent ecological mechanism. The most commonly cited support for partial regression coefficients in fragmentation research was generated under a restricted habitat cover range of less than 30%, where the amount–fragmentation relationship is approximately linear (Smith et al. 2009). Those results do not straightforwardly generalise to datasets spanning the full habitat-cover gradient. Partial regression coefficients are not direct measures of the relationship between predictors and the response when predictors are correlated (Graham 2003; Dormann et al. 2013). Partial coefficients can therefore generate erroneous interpretations under multicollinearity and cannot be taken as sufficient evidence that a predictor is ecologically unimportant (Ray-Mukherjee et al. 2014). Model inclusion alone secures none of these three conditions (Freckleton 2002; Lundberg et al. 2021).

Habitat fragmentation provides an unusually sharp test case for this broader identification problem because land-use change simultaneously reduces native cover, alters its spatial subdivision, and reshapes the surrounding matrix (Saunders et al. 1991; Fahrig 2003; Ewers and Didham 2006). Those changes co-generate the two predictors that additive models subsequently treat as independent (Trzcinski et al. 1999; Didham et al. 2012; Fletcher et al. 2018). Fragmentation research initially approached this through island biogeography (remnants as habitat islands surrounded by an inhospitable matrix), but syntheses showed that remnants are shaped by edge effects, connectivity, and matrix context, so fragmented systems must be understood in landscape rather than patch context (Saunders et al. 1991; Laurance 2008). That reframing made the design problem explicit. Habitat loss alters both the quantity of remaining habitat and its spatial subdivision, so amount and configuration are generated together before any landscape is sampled (Fahrig 2003; Trzcinski et al. 1999; Ewers and Didham 2006). Observed fragmentation effects can therefore be artefactual unless the two are analytically or structurally separated (Fischer and Lindenmayer 2007; Fletcher et al. 2007; Cattarino et al. 2014; Ruffell et al. 2016; Yaacobi et al. 2007; Laurance 2008).

The field has made only limited progress in operationally achieving that separation, even while increasingly treating the amount–configuration distinction as conceptually necessary (Hadley and Betts 2016; Riva et al. 2024b). The concept of habitat fragmentation per se (the ecological effect of spatial configuration independent of habitat amount) formalised the inferential target, and its estimation in observational landscapes requires statistical control of habitat amount because configuration metrics are typically correlated with amount in real landscapes (Fahrig 1997, 2002, 2003). The habitat amount hypothesis (HAH) extended this by proposing that species richness in equal-sized sample sites is determined primarily by the amount of suitable habitat in the local landscape, with no residual effect of patch size or isolation once habitat amount is accounted for (Fahrig 2013). That shifted the debate from whether patch attributes matter in principle to whether habitat amount and configuration can actually be separated in real observational landscapes. The HAH remains a hypothesis until rigorously tested under the specific site-scale conditions it requires (Fahrig 2015), and two further difficulties compound the problem. The local-landscape framing may be too narrow to address large-landscape viability questions, and disagreement persists over what the HAH predicts beyond the sample-site scale because those predictions hinge on spatial scales and response quantities the HAH does not directly address (Hanski 2015; Saura 2021a, 2021b; Fahrig 2021).

Whether habitat fragmentation affects biodiversity independently of habitat amount remains one of the most contested questions in landscape ecology and conservation science (Miller-Rushing et al. 2019; Fahrig et al. 2019; Fletcher et al. 2018). The disagreement is no longer only between different datasets or study systems. Two analyses of the same global multi-taxa dataset have now reached opposed conclusions from identical observations. Gonçalves-Souza et al. (2025a; hereafter GS25) reported lower α and γ diversity in fragmented landscapes using a categorical design contrast, accounting for among-pair variation in habitat amount with a random slope. GS25 additionally reported higher β diversity in fragmented than continuous landscapes, though this pattern disappeared when controlling for distance-decay effects (which GS25 interpreted as evidence against the species-turnover rescue hypothesis). Fahrig et al. (2026; hereafter F26) reanalysed the same landscapes with continuous, scale-matched predictors and concluded that no independent evidence for fragmentation remained once habitat amount was controlled for additively as a fixed effect. When identical data yield opposed conclusions under alternative predictor representations, the discrepancy itself becomes analytically informative. It exposes the inferential conditions under which fragmentation effects can and cannot be identified independently of habitat amount (Valente et al. 2023). Identifying those conditions requires asking whether the realised predictor geometry supports the contrast the analysis demands, whether the estimand corresponds to an ecological effect the data can identify, and whether the ecological conditions under which fragmentation effects are expected to be detectable have been explicitly accounted for.

Debate has therefore persisted not simply because different researchers favour different mechanisms, but because related questions have been evaluated under different design logics and geometric conditions, often without demonstrating that the predictor space provides the contrast required to identify the ecological quantity at issue independently (Lindenmayer and Fischer 2007; Didham et al. 2012; Hadley and Betts 2016; Riva et al. 2024b). Patch-scale and experimental studies typically measure the consequences of habitat-loss processes without additively partitioning their co-generated components, whereas landscape-scale additive analyses attempt to estimate an independent configuration effect conditional on habitat amount (Fahrig 2003; McGarigal and Cushman 2002; Fletcher et al. 2023a).

Compounding the geometric problem, fragmentation effects vary systematically with species trait-level sensitivity, matrix permeability, and habitat-amount thresholds (sources of ecological heterogeneity that make pooled inference across study contexts non-diagnostic even where predictor geometry is controlled) (Ewers and Didham 2006; Henle et al. 2004; Villard and Metzger 2014). A global multi-taxa dataset assembled from studies spanning diverse biomes, matrix types, and land-use histories inherits this heterogeneity as a structural property. Any analysis that treats it as random noise will produce results that are difficult to interpret ecologically regardless of how the predictor-geometry problem is addressed.

Statistical methods designed to separate habitat amount from configuration become unreliable when the two predictors are causally interdependent. Shared variance cannot be cleanly attributed to either predictor when both arise along a directed habitat-loss pathway (Didham et al. 2012). Amount–configuration dependence is a structural property of real landscapes rather than a nuisance sampling artefact, and the practical design corollary is that landscapes should be selected to minimise correlation between habitat amount and fragmentation metrics across the sample (Ewers and Didham 2006; Fletcher et al. 2018; Fahrig et al. 2019). Because these dependencies are often nonlinear (Koper et al. 2007), standard linear diagnostics do not adequately characterise whether the realised predictor space supports independent ecological interpretation. The unresolved question is therefore narrower than the broader amount-versus-fragmentation dispute: once continuous predictors are entered additively, has the sampled predictor geometry actually satisfied the separability condition that both sides of the debate already imply (Didham et al. 2012; Fletcher et al. 2018; Fahrig et al. 2019)?

The GS25–F26 exchange provides an empirical demonstration of these identification conditions within a debate characterised as locked in a half-century demonstration-and-counter-demonstration cycle (Valente et al. 2023). The discrepancy is therefore useful not as another demonstration in the cycle, but as evidence about whether the underlying design and model architecture can identify the ecological contrast under dispute (Valente et al. 2023). The discrepancy itself may be an instance of a known phenomenon. Suppressor attenuation under structural predictor coupling can generate opposed conclusions from identical datasets without any difference in the underlying ecology (Smith et al. 2009; Prunier et al. 2015). In a cross-over suppressor structure, one predictor aligns strongly with the fitted response gradient (measured by its structure coefficient) while its partial coefficient, measured by its beta weight, is attenuated by the other (Smith et al. 2009; Ray-Mukherjee et al. 2014). The predictor that aligns strongly but is attenuated is the subordinate predictor. The predictor that absorbs shared gradient variance and is partially inflated is the dominant predictor. Whether the GS25–F26 divergence reflects this architecture (with fragmentation as the subordinate predictor suppressed by its coupling with habitat amount along a shared habitat-loss gradient; Smith et al. 2009; Fletcher et al. 2018) is the diagnostic question the present analysis addresses.

By replacing a categorical landscape contrast with two continuous predictors measured within the same spatial extent, F26 addressed a genuine limitation GS25 had acknowledged (Gonçalves-Souza et al. 2025a; Fahrig et al. 2026), though that redesign does not by itself validate the stronger inferential claim built on top of it (Freckleton 2002; Ruffell et al. 2016; Fletcher et al. 2018). The issue is whether the redesign produced a separable predictor geometry. Once the predictors are recast in additive continuous form, do they span sufficiently distinct dimensions of predictor space to support an independent interpretation of the configuration coefficient, or do they remain embedded in the same habitat-loss gradient (Freckleton 2002; Ruffell et al. 2016)? The landscape-scale analytical approach itself implies the same design corollary: when correlation between habitat amount and fragmentation metrics is high, landscapes should be selected to minimise that correlation across the sample (Fahrig et al. 2019). Whether that condition was demonstrated in the realised predictor space of the GS25–F26 redesign is what the present analysis evaluates.

Because both analyses draw on the same underlying dataset (the same 37 study pairs and the same biodiversity observations), the remaining source of discrepancy cannot be attributed to different study systems, taxa, or spatial locations. It lies in how the alternative predictor representations allocate shared variance when habitat amount and configuration trace a common nonlinear landscape gradient. The analytical framework both analyses share assumes orthogonal predictor dimensions (Freckleton 2002; Ruffell et al. 2016; Fletcher et al. 2018), but whether the realised landscape gradient provides them cannot be assumed and must be demonstrated (Fahrig 2003; Koper et al. 2007; Villard and Metzger 2014).

Habitat fragmentation as a process, the breaking apart of habitat into smaller and more isolated patches, is conceptually distinct from fragmentation as pattern, the spatial configuration of the remaining habitat mosaic at a given point in time (Fahrig 2003). Fragmentation per se is the independent effect of spatial configuration on biodiversity at a given habitat amount, that is, the biodiversity difference between landscapes that differ in spatial arrangement but not in the amount of habitat (Fahrig 2003, 2013, 2017; Fahrig et al. 2019). That quantity is a pattern-level estimand, but the process that generates both pattern measurements co-produces them along the same habitat-loss gradient, so they are not separable in the way the additive framework assumes (Didham et al. 2012; Fletcher et al. 2018). This non-equivalence helps explain why the GS25–F26 discrepancy mirrors a broader divergence between patch-scale studies, which more often report negative fragmentation effects, and landscape-scale studies, which more often report null or positive fragmentation effects when detectable (Fahrig 2017; Fahrig et al. 2019). The debate is often posed as if habitat amount and configuration were competing parallel-additive explanations for the same biodiversity variance, so that support for one diminishes the other (Didham et al. 2012; Hadley and Betts 2016; Valente et al. 2023). That framing already assumes the two predictors can be cleanly separated in the realised landscape geometry.

Configuration metrics based on patch enumeration often do more than vary nonlinearly with habitat amount. Across class-level metrics, many respond jointly to area and aggregation, whereas only a minority are predominantly aggregation-responsive and relatively independent of area (Neel et al. 2004; Wang et al. 2014; Villard and Metzger 2014; Zhang et al. 2024; Martin et al. 2025a). This nonlinear coupling is not detectable by standard linear diagnostics, making linear-correlation tests unreliable as evidence of predictor separability (Dormann et al. 2013; Liu et al. 2024). The relationship between habitat amount and patch-number metrics is nonlinear and asymmetric (as habitat cover declines from continuous, patch count rises sharply around the percolation threshold before declining gradually as cover decreases further), a structural geometric property of habitat-loss landscapes, not a contingent feature of any particular dataset or sampling design (Andrén 1994; Fahrig 2003). Within fixed-area sampling units this coupling is partly geometrically enforced. Total habitat area (Aₗ), mean patch size (Āₚ), and patch number (Nₚ) satisfy the identity Āₚ = Aₗ / Nₚ (Fletcher et al. 2023a), making patch-number metrics functionally dependent on habitat amount through the subdivision process itself, with the high-amount/high-fragmentation corner of the predictor space constrained by this identity and the low-amount/low-fragmentation corner constrained only by ecological rarity.

In datasets generated by progressive habitat loss, where amount and configuration are co-produced along the same landscape gradient, this structural dependence is unlikely to be resolved by landscape selection alone, because the process generates the coupling in the landscape itself rather than in the analyst’s parameterisation (Fahrig 2003; Didham et al. 2012; Koper et al. 2007; Ruffell et al. 2016). That co-production does not require fragmentation to increase monotonically with habitat loss across all metrics and scales; at global extent, forest loss has been associated with both increases and decreases in fragmentation depending on fragmentation measure, landscape size, biome, and initial forest amount (Martin et al. 2026). Under this geometry, near-null additive fragmentation coefficients are the expected product of the modelling framework itself unless predictor separability has been explicitly verified (Fletcher et al. 2023a; Smith et al. 2009; Ray-Mukherjee et al. 2014).

Design failure has persisted despite decades of methodological clarification. Only 18% of fragmentation studies controlled for habitat amount in the decade following the earliest calls for separation, with no trend toward increased separation over time (Hadley and Betts 2016), and that figure remained at approximately 20% in a subsequent audit a decade later (Riva et al. 2024b). The more stringent requirement (explicitly separating habitat area from configuration effects rather than merely controlling for one) was met by only 6% of all fragmentation studies and 3% of experimental studies published between 1995 and 2000, with only one landscape-scale study achieving both separation and rigorous controls (McGarigal and Cushman 2002). In a systematic review of primate fragmentation studies, 74% attributed observed patterns to fragmentation despite not controlling for habitat loss, and 53% concluded fragmentation caused the observed pattern even though all studies were conducted at the fragment rather than the landscape scale (Arroyo-Rodríguez et al. 2013). This persistent design and interpretive entanglement have contributed to a debate whose discordance may reflect competing model architectures and study logics as much as genuine ecological disagreement (Didham et al. 2012; Hadley and Betts 2016; Arasa-Gisbert et al. 2021; Valente et al. 2023). The GS25–F26 case operationalises that diagnosis by testing whether predictor geometry prevents additive habitat-amount control from isolating the ecological contrast that fragmentation-per-se inference requires.

The process-level source of the problem, its geometric consequence, and its inferential implication are linked (Fig. 1). Habitat loss co-generates amount and configuration along a directed, asymmetric gradient before any landscape is sampled. The dashed arrow from Amount to Fragmentation encodes derived coupling rather than an independent causal pathway (Fig. 1a). The resulting predictor space is therefore asymmetric and constrained. The realised sample concentrates at high habitat amount and few patches, with a thin tail toward low amount and intermediate configuration (Villard and Metzger 2014; Andrén 1994; Fig. 1b). That geometric ceiling constrains both the statistical detectability of fragmentation effects and the behaviour of configuration metrics across the sampled gradient, independently of sample size or study design (Neel et al. 2004; Wang et al. 2014; Fig. 1c–d).

**Fig. 1.**
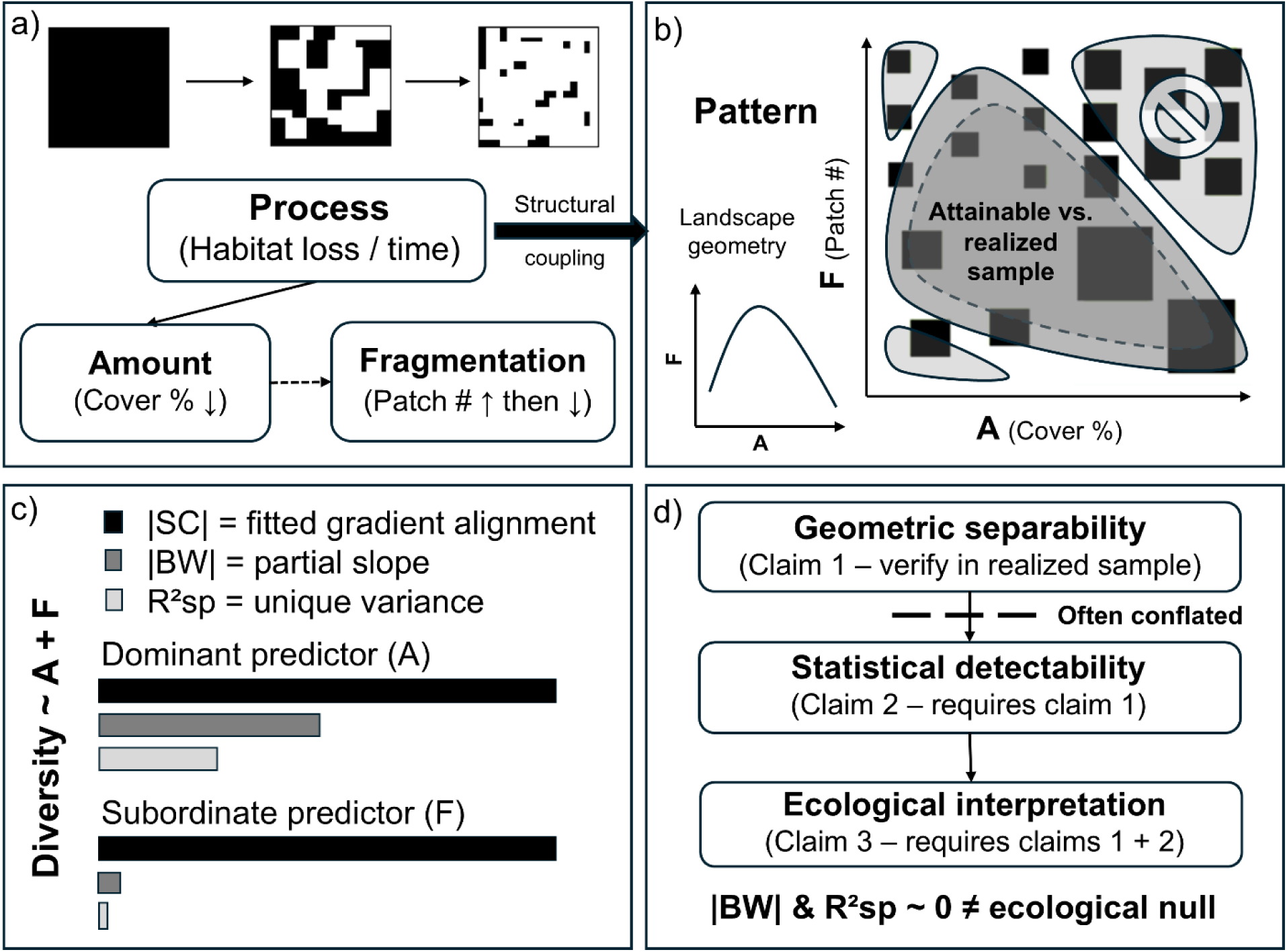
From process to inference: how habitat-loss-driven predictor coupling produces a non-diagnostic additive null. (a) Process view: habitat loss co-generates habitat amount and configuration along a directed, asymmetric gradient (structural coupling). The dashed arrow from Amount to Fragmentation encodes the derived geometric coupling direction. The inset curve shows the schematic nonlinear (asymmetric, right-skewed hump-shaped) relationship between habitat amount and patch number — steep rise as continuous habitat begins fragmenting, gradual descent as patches coalesce at low cover — the geometric basis for the attainable sample. (b) Pattern view: the outer boundary is the attainable predictor space — the geometrically possible region defined by the subdivision identity; the shaded cloud is the realised sample (dashed boundary), which is at most equal to the attainable space and typically smaller. The upper-right zone (high amount, many patches) lies outside the attainable boundary; the lower-left zone is constrained only by ecological rarity. Landscape icons outside the shaded region represent configurations structurally sparse or absent in observational datasets generated by habitat loss. The attainable ceiling constrains both statistical detectability of fragmentation effects and the validity of configuration metrics across the sampled gradient. (c) Suppressor signature in Diversity ∼ A + F (schematic): |SC| = alignment with the model-fitted diversity gradient; |BW| = unique partial slope; R²sp = unique variance contribution. A = dominant predictor (habitat amount); F = subordinate predictor (fragmentation). Bar lengths are schematic; empirical values in Table 1. (d) Inferential cascade: the three conditions required for independent configuration inference and the short-circuit produced when additive regression addresses claim 2 without verifying claim 1. The bottom note |BW| & R²sp ∼ 0 ≠ ecological null summarises the central inferential consequence.

**Table 1.** Suppressor diagnostics across model parameterisations for α and γ diversity (medians across 6 analytical variants: 3 diversity-order metrics × 2 pairing designs). SC = structure coefficient (correlation between predictor and model-fitted response ŷ); measures gradient alignment. BW = standardised beta weight (partial regression coefficient per 2 SD); measures unique predictor contribution. The cross-over suppressor pattern is: large |SC| with BW ≈ 0 for the subordinate predictor, despite that predictor aligning strongly with the fitted response gradient (Smith et al. 2009). Parameterisations: M1 = additive F26 specification (diversity ∼ amount + fragmentation); M2 = fragmentation residualised on amount; M3 = amount residualised on fragmentation. Column headers use A (habitat amount, forest cover) and F (fragmentation, patch number) as shorthand. Coupling criterion: residual GAM R² < 0.02 in both axis orientations (bidirectional orthogonality threshold). M3 is the only parameterisation satisfying this criterion in all sections. M4 (negative control, not shown) reintroduces strong coupling. Sections: A = full dataset; B1 = geometric robustness check (≥50% cover, study-level minimum filter — both landscapes must exceed 50%); B2 = biome conditionality check (tropical South America, where the M3 amount SC flips negative, indicating configuration dominates residual diversity variation once the habitat-loss gradient is removed).

|  | M1 — Additive (F26) | M2 — A + resF | M3 — resA + F |
| --- | --- | --- | --- |
| | $\alpha / \gamma$ | $\alpha / \gamma$ | $\alpha / \gamma$ |
| **A. Full dataset (n = 37 studies = 74 landscape pairs)** |  |  |  |
| *Coupling criterion* | fails | fails | met |
| SC - Habitat amount | +1.00 / +0.99 | +0.98 / +0.97 | +0.44 / +0.42 |
| BW - Habitat amount | +0.11 / +0.11 | +0.13 / +0.12 | +0.04 / +0.05 |
| SC - Fragmentation | -0.72 / -0.75 | -0.20 / -0.22 | -0.90 / -0.91 |
| BW - Fragmentation | -0.01 / -0.02 | -0.03 / -0.03 | -0.11 / -0.10 |
| **B1. Global $\geq 50\%$ cover (n = 17 studies; both landscapes $\geq 50\%$ · Tier 1)** |  |  |  |
| *Coupling criterion* | fails | fails | met |
| SC - Habitat amount | +0.94 / +0.94 | +0.96 / +0.94 | +0.36 / +0.34 |
| BW - Habitat amount | +0.06 / +0.05 | +0.12 / +0.10 | +0.03 / +0.03 |
| SC - Fragmentation | -0.95 / -0.95 | -0.27 / -0.35 | -0.93 / -0.94 |
| BW - Fragmentation | -0.08 / -0.06 | -0.03 / -0.04 | -0.12 / -0.10 |
| **B2. Tropical South America (n = 23 studies; 0–100% cover · Tier 1)** |  |  |  |
| *Coupling criterion* | fails | fails | met |
| SC - Habitat amount | +0.95 / +0.82 | +0.95 / +0.82 | -0.09 / -0.26 |

|  | **M1 — Additive (F26)** | **M2 — A + resF** | **M3 — resA + F** |
| --- | --- | --- | --- |
| BW - Habitat amount | +0.08 / +0.04 | +0.11 / +0.10 | −0.01 / −0.04 |
| SC - Fragmentation | −0.87 / −0.90 | −0.31 / −0.54 | −1.00 / −0.97 |
| BW - Fragmentation | −0.05 / −0.11 | −0.04 / −0.08 | −0.11 / −0.14 |

In a generic additive model, Diversity ∼ A + F, A denotes habitat amount and F denotes fragmentation. The fitted diversity gradient is the model-predicted surface defined jointly by both predictors. Structure coefficients (|SC|) measure predictor alignment with that gradient. Beta weights (|BW|) measure the unique partial slope, and semi-partial R²sp measures unique variance contribution (defined formally in Methods; Fig. 1c). When predictors share a dominant gradient, the subordinate predictor can show equally large SC together with near-zero BW and R²sp, the joint diagnostic signature of cross-over suppressor attenuation (Ray-Mukherjee et al. 2014). If fragmentation is the subordinate predictor and habitat amount the dominant one, fragmentation would be expected to track the fitted diversity gradient as strongly as habitat amount while contributing negligible unique variance once habitat amount is included. Independent interpretation of an additive configuration coefficient requires three conditions to be verified in sequence. The predictors must be geometrically separable in the realised predictor space. The separable component must correspond to an ecological effect the estimand can identify. The study context must be stratified in a way that makes pooled inference ecologically interpretable. Additive regression addresses the second condition without verifying the first, and neither addresses the third. Near-zero |BW| and R²sp do not constitute an ecological null when separability remains unverified and ecological context is unmodelled (Fig. 1d).

Fragmentation per se is the contrast between landscapes with equal habitat amount but different configuration (Fahrig 2003; Fahrig et al. 2026). Whether the additive reanalysis actually isolates that contrast depends on the realised geometry of the predictors in the sampled landscapes, not on the formal inclusion of both variables in the same model. Additive regression can correctly estimate partial coefficients even when predictors share strong nonlinear gradients, but those coefficients need not identify independent ecological effects when predictor geometry constrains variance attribution (Freckleton 2002; Ruffell et al. 2016). Residualising predictors does not by itself resolve that problem, because shared gradient variance is reallocated rather than removed (Koper et al. 2007; Graham 2003). Fragmentation-per-se inference depends on landscapes sampling habitat amount and configuration with enough independence that partial coefficients do not merely divide a common gradient between model terms (Smith et al. 2009; Fletcher et al. 2018; Fahrig et al. 2019). Whether the continuous redesign achieves that condition in the realised predictor space is an empirical question, not one answered by specifying both variables in the same model.

The central question is not whether controlling for habitat amount is preferable to leaving it uncontrolled, but whether the sampled predictor geometry actually furnishes the contrast that an independent configuration coefficient is meant to represent. In the GS25–F26 dataset, that link between coefficient and ecological estimand remains unresolved under the shared habitat-loss gradient (Freckleton 2002; Ruffell et al. 2016; Fletcher et al. 2018; Lundberg et al. 2021). Diagnosing that gap requires tools the methodological literature has developed but not yet assembled and applied to a large-scale empirical fragmentation dataset (Koper et al. 2007; Smith et al. 2009; Ruffell et al. 2016; Ray-Mukherjee et al. 2014; Fletcher et al. 2018). Predictor geometry is not the only source of ambiguity. Fragmentation effects vary systematically with trait-level sensitivity, matrix permeability, and habitat-amount thresholds, so pooled additive analyses across ecologically heterogeneous study contexts may be non-diagnostic even where predictor separability is demonstrated (Ewers and Didham 2006; Villard and Metzger 2014).

Using the GS25–F26 case, we ask why comparable data continue to yield opposed conclusions in the fragmentation debate even after habitat amount is controlled. The analysis evaluates whether that divergence reflects a prior inferential problem: when habitat amount and configuration are co-generated as habitat is lost (Fig. 1a–b), a near-zero additive fragmentation coefficient cannot by itself decide whether configuration is ecologically irrelevant (Fig. 1c–d). Rather than resolving the ecological question directly, the analysis establishes a prior inferential condition for when landscape evidence can address it. Specifically, it tests whether the additive fragmentation null in this dataset carries the diagnostic signature of a suppressor constraint rather than an absent ecological effect, and whether that constraint traces to the landscape data-generating process rather than to any single analytical choice. Distinguishing these cases has direct consequences for how comparative fragmentation evidence is evaluated, which metrics are selected, and what study designs can prospectively resolve the debate. The analysis documents what process-driven predictor coupling and unmodelled ecological heterogeneity produce in practice for those coefficients, and what that pattern reveals about the inferential conditions the debate requires.

## Methods

### Data

The dataset compiled and analysed by F26 was used here; it comprises paired continuous and fragmented landscapes from 37 multi-taxa studies originally assembled as the LandFrag database (Gonçalves-Souza et al. 2025a,b). Each study contributed landscape-level measurements of forest cover (habitat amount, expressed as proportion of 2-km radius buffer) and number of forest patches within the same buffer, paired across continuous and fragmented landscape types. The cover distribution in the dataset is right-skewed and concentrated at higher values (median > 50%), with a smaller number of landscapes in the low-cover range where the nonlinear constraint between patches and amount is steepest; full distributional statistics are reported in Supplementary Material S1 (Section S1.1). Three diversity components were analysed: α diversity (mean species richness per plot), β diversity (assemblage dissimilarity between paired plots), and γ diversity (total species richness per landscape). Observed richness and two rarefied Hill number orders (q = 0 weighting rare species, q = 2 weighting abundant species) were evaluated, producing 18 analytical variants (3 diversity components × 2 pairing designs [all pairs; nearest-neighbour pairs] × 3 measures [observed; q = 0; q = 2]). Response variables were log-transformed where this improved residual normality, consistent with F26.

### Predictor geometry and orthogonality diagnostics

The bivariate relationship between number of patches and habitat amount was characterised using linear regression and generalised additive models (GAMs; mgcv package, Wood 2017; Pedersen et al. 2019; Wang et al. 2014). Nonlinear coupling was quantified in both directions by fitting habitat amount as a smooth function of number of patches and number of patches as a smooth function of habitat amount. GAM R² was then compared with the corresponding linear R² in each direction. The asymmetry ratio was defined as the reverse nonlinear component, amount ∼ s(patches), divided by the forward component, patches ∼ s(amount). Values greater than 1 indicate that habitat amount is more nonlinearly constrained by patch number than the reverse — the dominant direction of the shared gradient. Assessing both directions is necessary because GAM R² is not symmetric. The proportion of variance explained depends on which variable is treated as the response, so restricting the diagnostic to one direction can understate or mischaracterise the association. Both GAM R² values and the nonlinear component (GAM R² minus linear R²) are reported for each direction; directional asymmetry in the nonlinear component is interpreted as evidence consistent with directed coupling. Full coupling outputs are in Supplementary Material S1 (Section S1.2).

Standard linear collinearity diagnostics (Pearson r, Spearman ρ, and variance inflation factors, VIF) were computed as reference values. These metrics were not used as the primary coupling assessment because their sensitivity is restricted to linear dependence; nonlinear coupling that persists after the linear association is removed is invisible to them by construction (Dormann et al. 2013; Liu et al. 2024). Residualisation was assessed by testing whether residual nonlinear coupling was reduced to near zero. This is a stricter condition than zeroing the linear correlation. Koper et al. (2007) showed that zero linear correlation is insufficient when amount–fragmentation coupling is nonlinear. The simulation benchmark most commonly cited to support partial regression coefficients as unbiased estimators in fragmentation research was derived from landscapes with less than 30% habitat cover, where the amount–fragmentation relationship is approximately linear (Smith et al. 2009). The present dataset spans the full cover gradient and therefore includes the nonlinear regime that simulation did not examine. Additive specifications spanning the residualisation space between the two predictors were then evaluated.

### Model parameterisations

Three primary parameterisations and one negative control were evaluated. The F26 specification, diversity as an additive function of raw fragmentation and raw amount (M1), was taken as the baseline. When fragmentation is residualised on habitat amount using a GAM (M2), the resulting predictor represents the component of configuration statistically independent of habitat amount under a linear model and is a commonly recommended response to landscape collinearity (Koper et al. 2007; Graham 2003). When habitat amount is residualised on fragmentation using a GAM (M3), the predictor represents the component of forest cover statistically independent of configuration. Both M2 and M3 are diagnostic decompositions of the shared habitat-loss gradient rather than ecological models of the original predictors. Neither is conceptually superior to the other, and both were evaluated against the same residual-uncoupling criterion prior to any analysis of coefficient patterns. A negative-control parameterisation involving bidirectional residualisation of both predictors (M4) was evaluated to document the full gradient variance allocation space: M4 allocates the shared gradient primarily to the fragmentation axis and is expected to produce the mirror image of M1, with large negative fragmentation BW and near-zero amount BW. The residual-coupling threshold (GAM R² < 0.02 in both diagnostic orientations) was used as a conservative, dataset-specific operational definition of near-zero residual nonlinear dependence. Full residual-coupling values and parameterisation-sequence results are in Supplementary Material S1 (Section S1.2). Residualisation used GAMs with thin-plate splines (basis dimension k selected by generalised cross-validation; results unchanged for k = 5 to k = 10).

### Model fitting

All primary models were fitted using lme4::lmer (Bates et al. 2015) with Gaussian errors and random intercepts by study (1|refshort), estimated by restricted maximum likelihood (REML), matching the model specification used by F26. A parallel set of 72 models with identical formula and random structure was fitted using glmmTMB (Brooks et al. 2017) with maximum likelihood (ML) for AICc comparisons, as ML estimation is required for valid fixed-effect model comparison (Burnham and Anderson 2002). To facilitate numerical convergence and improve comparability of effect sizes across predictors, all continuous fixed-effect predictors were centred and scaled by dividing by two standard deviations prior to model fitting (Gelman 2008; Schielzeth 2010). Beta weights therefore represent partial effects on the linear predictor per two-standard-deviation change in each predictor, making magnitudes comparable across predictors within each model. Coefficient values reported throughout derive exclusively from the REML models.

### Suppressor geometry and variance partitioning

Suppressor geometry was diagnosed using structure coefficients (SC), beta weights (BW), and semi-partial R² (R²sp) extracted via partR2 (Stoffel et al. 2022; nboot = 1000, CI = 0.95). partR2 models were fitted with ML, as required for valid fixed-effect model comparison across nested structures. Fixed-effect estimates (BW, SC) are numerically identical under REML and ML in these models. Parametric bootstrapping was used to attach repeated-sampling uncertainty to BW and variance-decomposition quantities, improving on reliance on single point estimates alone (Fieberg et al. 2020). This was particularly relevant for evaluating whether the joint suppressor signature — large |SC|, near-zero BW, and near-zero semi-partial R² — was stable under repeated sampling from the fitted model rather than a single-sample artefact. Formal definitions of structure coefficients, beta weights, the SC² = R²semi-partial + R²shared decomposition, and the algebraic basis of suppressor attenuation are in Supplementary Material S3 (Sections S3.1.2–S3.1.3). Medians and ranges for SC, BW, and R²sp are reported across six variants within each diversity component (3 metrics × 2 pairing designs) for α and γ diversity. Marginal R² (mR²) and conditional R² (cR²) were computed following Nakagawa and Schielzeth (2013) using the MuMIn package (Bartoń 2024). Ray-Mukherjee et al. (2014) provide the methodological precedent for the joint SC + BW + R²sp suppressor diagnostic. The algebraic derivation is in Supplementary Material S3 (Section S3.1.2).

### Design-reconstruction diagnostic

The suppressor geometry was additionally evaluated using a mixed-effects logistic regression of the GS25 categorical classification (continuous vs. fragmented landscape type) onto the two F26 continuous predictors (standardised number of patches and habitat amount) with study as a random intercept, testing whether the same SC–BW–R²sp mismatch structures not only the diversity response but the design correspondence itself (full outputs in Supplementary Material S1 (Section S1.1.1)). The Tjur R² was computed as the difference in mean fitted probabilities between fragmented and continuous landscapes (Tjur 2009).

### Model adequacy and subset diagnostics

Subsets were assigned to one of three evidential tiers based on sample size prior to analysis: Tier 1 (n ≥ 20 studies) received the full SC+BW+semi-partial R² suppressor tripod; Tier 2 (10 ≤ n < 20) received SC and BW only; Tier 3 (n < 10) received directional indicators only, with no independent inferential claims and no path models. Additional subset analyses were used for two distinct purposes: to test whether the full-dataset suppressor geometry was driven by particular portions of the predictor space, and to assess whether that geometry persisted within the continental grouping used by GS25. Subset analyses were conducted at two levels: robustness checks on full-dataset geometry across predictor-space regions (≥50% cover; tropical-only), and continental sensitivity checks (South American vs non-South American) paralleling GS25’s grouping.

In all subsets, coupling diagnostics, residualisation, and mixed models were rebuilt within the subset; M3 coefficients were interpreted only when the residual-uncoupling criterion was satisfied within that subset. Models were specified to account for the dependence structure of the data following GS25-F26 (study ID as random intercept), and assumptions were evaluated separately from the variance-allocation diagnostics reported below. Additional crossed sensitivity checks evaluated whether the suppressor diagnosis depended jointly on climatic (temperate vs. tropical) and continental composition (e.g., South America subset); these used the same predeclared evidential tier framework and are reported fully in Supplementary Material S2 (Section S2.2.4). A complete summary of all subset analyses, their sample sizes, ecological rationale, analytical scope, and inferential tier is provided in Supplementary Material S2 (Section S2.2), Table S2.8. Consistent with current guidance in statistical ecology, model adequacy was treated as a prerequisite for interpretation (Popovic et al. 2024).

Model adequacy was evaluated using DHARMa simulation-based residual diagnostics (Hartig 2022), Levene tests for residual variance homogeneity across landscape types, and random-effect stability checks across parameterisations. Full implementation details and outputs are in Supplementary Material S2 (Section S2.4).

A path model was fitted across all 18 analytical variants as a complementary causal-hierarchy check, following the directed Amount → Fragmentation → Diversity prior proposed by Didham et al. (2012). The model comprised a structural path (frag_z ∼ amount_z) and a response path (diversity ∼ frag_z + amount_z), both with random intercepts by study. Standardised path coefficients use conventional 1-SD standardisation and are not on the same scale as the 2-SD beta weights reported in Table 1 (Gelman 2008). Full implementation details, all 18-variant results, R², AICc comparisons, and continental extensions are in Supplementary Material S2 (Section S2.3).

## Results

### Predictor coupling and orthogonality

The landscape-level dataset spans a median habitat cover of 70% (range 34.5–98.4%) across all 74 landscape pairs, placing the sample predominantly in the nonlinear coupling regime above the ∼30% cover threshold where amount–fragmentation relationships become approximately linear. Number of patches and habitat amount exhibited nonlinear coupling in this dataset, with GAM R² exceeding linear R² in both axis orientations of the diagnostic (Fig. S1.1; full values in Supplementary Material S1 (Section S1.2)). The nonlinear component was asymmetric, being stronger when habitat amount was treated as the response and number of patches as the predictor (GAM R² = 0.70, nonlinear component = 0.243) than in the forward orientation, where number of patches was treated as the response and habitat amount as the predictor (GAM R² = 0.49, nonlinear component = 0.032), while the linear R² was identical in both directions (R² = 0.45). The reverse nonlinear component was thus 7.75 times larger than the forward (ratio computed from unrounded pipeline values; Supplementary Material S1 (Section S1.2) reports rounded values). Pearson r = −0.67 and VIF = 1.83 satisfied standard linear collinearity thresholds (|r| < 0.7; VIF < 5), yet the nonlinear component in the reverse direction represents a 55% amplification of coupling strength beyond what these linear metrics detect. In this dataset, a predictor pair satisfying both conventional thresholds nevertheless produced a ∼51-fold SC-to-BW discrepancy (α median SC = −0.725; α median BW = −0.014) — the defining signature of cross-over suppression.

### Design-reconstruction diagnostic

The pairwise directional structure of the predictor space, i.e. fragmented landscapes simultaneously having more patches and lower habitat amount in 31 of 37 studies (84%), is reported in Supplementary Material S1 (Section S1.1.2). A mixed-effects logistic regression of the GS25 categorical classification (continuous vs. fragmented) on the two F26 continuous predictors, with study as a random intercept, produced the same SC–BW mismatch as the diversity regression. SC for habitat amount = −0.985 and SC for patches = +0.708 (both large), while BW for patches was near zero (BW = +0.048) and its semi-partial R² was negative (R²sp = −0.10%) against R²sp = +2.60% for amount. The SC–BW–R²sp mismatch was present in both the diversity regression and the logistic classification model (full diagnostics in Supplementary Material S1 (Section S1.1.1)).

### Suppressor geometry in the additive specification

In the F26 additive specification (M1), fragmentation structure coefficients were large and consistently negative for α and γ diversity (α: median SC = −0.72, range −0.91 to −0.61; γ: median SC = −0.75, range −0.92 to −0.66), while fragmentation beta weights were near zero across all 18 analytical variants (α: median BW = −0.01; γ: median BW = −0.02), with sign reversal in two of 18 variants. Amount structure coefficients were large and consistently positive (α median = +0.997; γ median = +0.992), while amount beta weights were substantial and positive. Semi-partial R² confirmed the same pattern: fragmentation contributed a median of 0.006% of unique response variance for α diversity and 0.02% for γ, against a model marginal R² of approximately 0.017, yielding an ∼8,800-fold discrepancy between fragmentation’s inclusive and unique variance contributions for α diversity (∼3,000-fold for γ), and a separately ∼51-fold discrepancy between |SC| and |BW|. Together, large negative SC, near-zero BW, and near-zero R²sp constitute the joint suppressor tripod across all 18 analytical variants and both diversity components (Figs. 2–4; Table 1; Supplementary Material S2 (Fig. S2.2)).

**Fig. 2.**
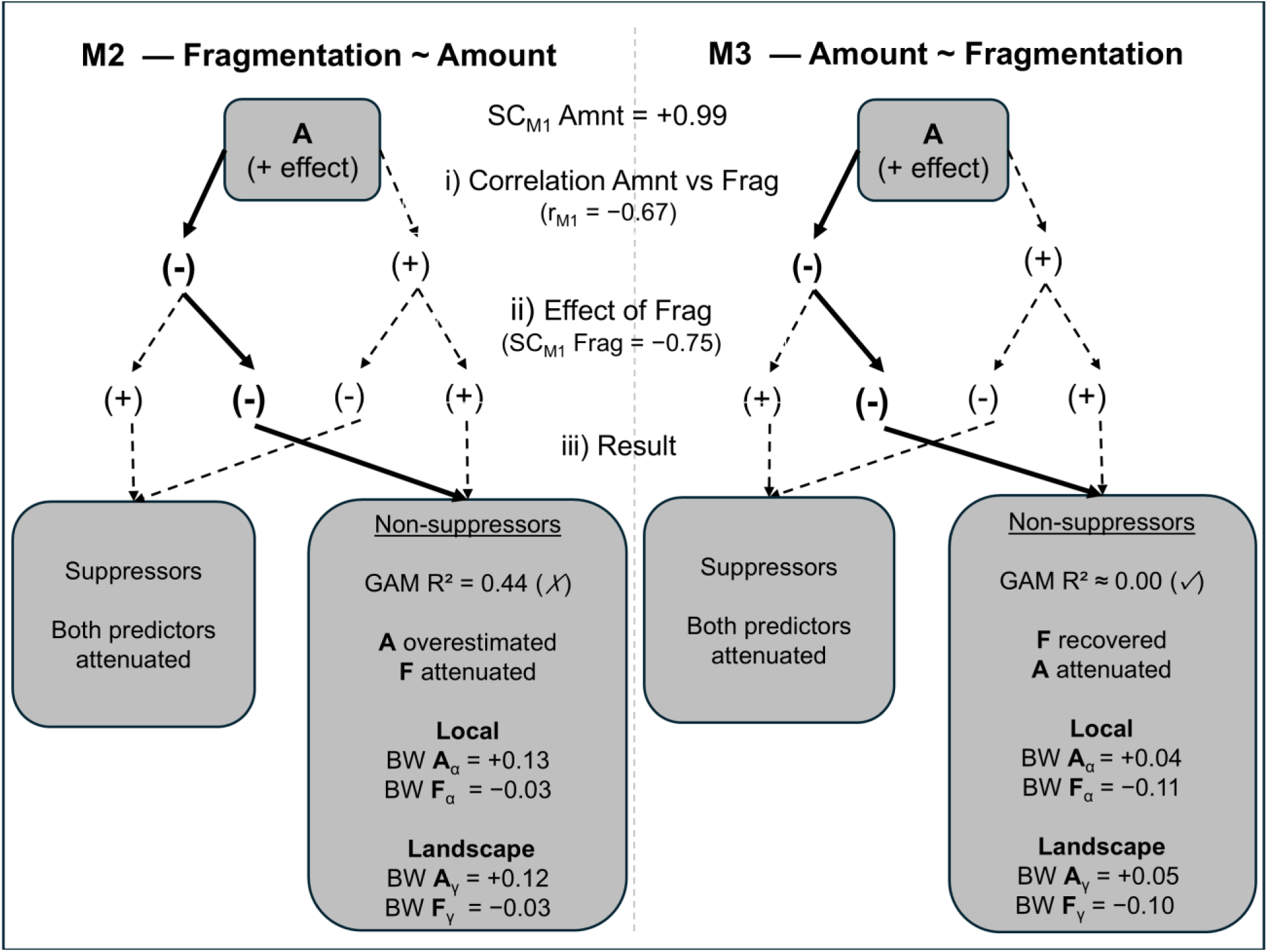
Suppressor geometry under directed predictor residualisation: M2 versus M3. The diagram traces how negative predictor correlation (r = −0.67) and the fragmentation alignment with the fitted diversity gradient (SC_M1 Frag = −0.75, γ) propagate through each residualisation direction to determine whether coefficient attenuation is eliminated or preserved. Three factors operate jointly (rows i–iii): the predictor correlation, the effect direction of fragmentation, and their interaction through the residual predictor. Under M2 (fragmentation residualised on amount), the non-suppressor path retains substantial reverse-direction nonlinear coupling (GAM R² = 0.44) leaving habitat amount inflated and fragmentation attenuated. Under M3 (amount residualised on fragmentation), the non-suppressor path achieves near-zero residual coupling in both orientations (GAM R² ≈ 0.00; ✓), enabling diagnostic recovery of the fragmentation signal and attenuation of amount. The directional asymmetry of the underlying nonlinear coupling — 7.75× stronger in the reverse orientation, amount ∼ s(patches), than in the forward orientation, patches ∼ s(amount) (16.2× in the ≥50% cover subset) — provides the process-level justification for the M3 residualisation direction. Values shown are γ-diversity medians across six analytical variants; α-diversity results are shown in the boxes for reference and closely parallel γ. Full variance decompositions in Figs. S2.2–S2.3 (Supplementary Material S2) and Table 1.

**Fig. 3.**
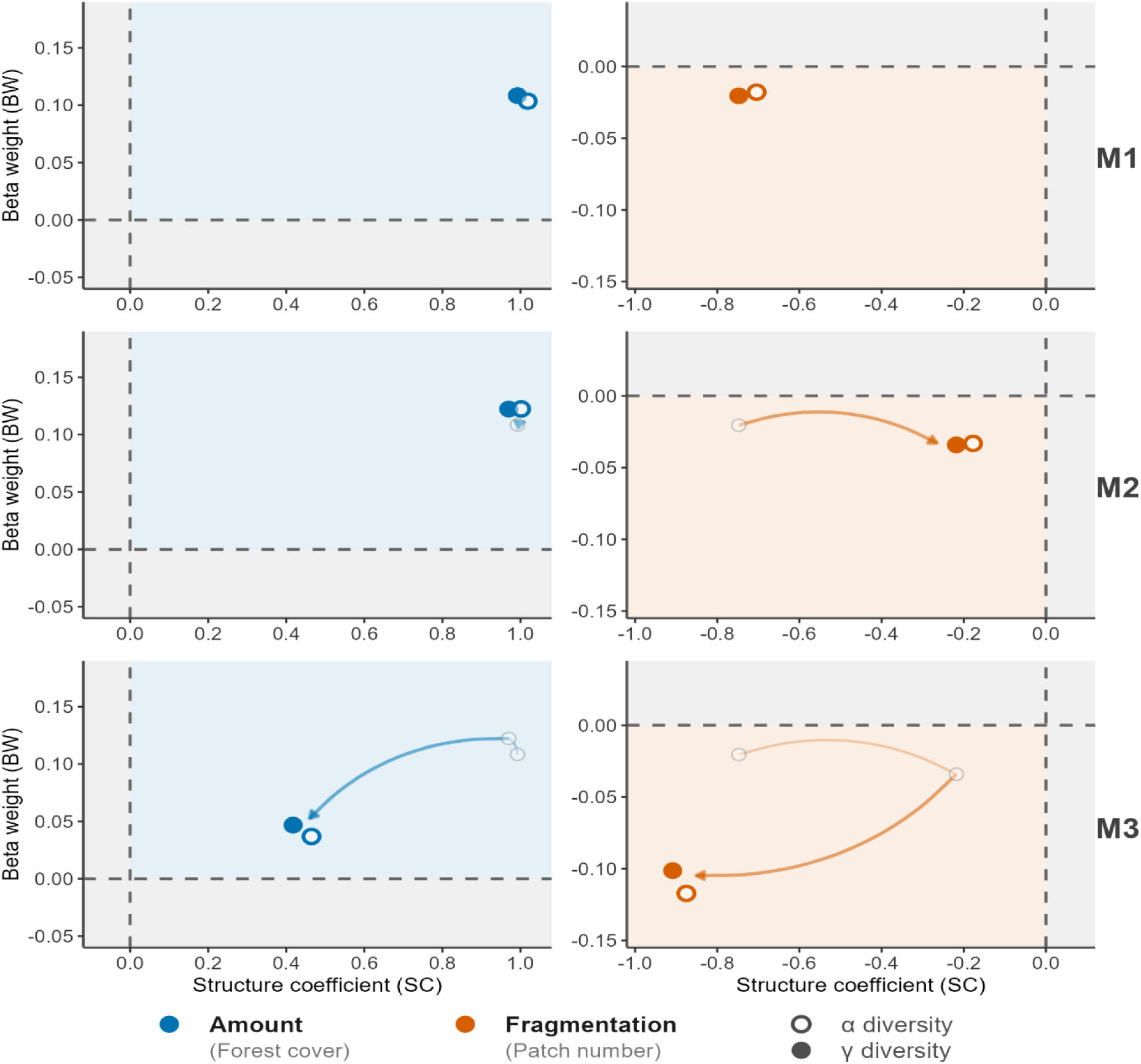
SC–BW dissociation and gradient reallocation across parameterisations M1–M3. Each panel shows the structure coefficient (SC; x-axis) against the standardised beta weight (BW; y-axis) for habitat amount (left column, blue) and fragmentation (right column, orange) across α and γ diversity (open and filled circles, respectively). Coloured backgrounds delimit the coherent-effect quadrant for each predictor: upper-right for amount (positive SC, positive BW) and lower-left for fragmentation (negative SC, negative BW). Arrows in M2 and M3 panels trace the shift in point position relative to M1, quantifying how residualisation reallocates gradient variance. Under M1, fragmentation sits in the upper region of the negative-SC half-plane with BW near zero — the defining cross-over suppressor signature: the predictor aligns strongly with the fitted diversity gradient (large |SC|) but contributes negligible independent variance once amount is controlled (BW ≈ 0). Under M2, fragmentation remains in the suppressor zone with a reduced |SC|, consistent with partially retained coupling. Under M3, fragmentation moves into the lower-left quadrant (both SC and BW negative), while the amount point shifts as the shared gradient variance is redistributed to the orthogonalised fragmentation term. Points are medians across 6 analytical variants: full variant-level decompositions in Fig. 4 and Supplementary Material S2 (Section S2.1).

**Fig. 4.**
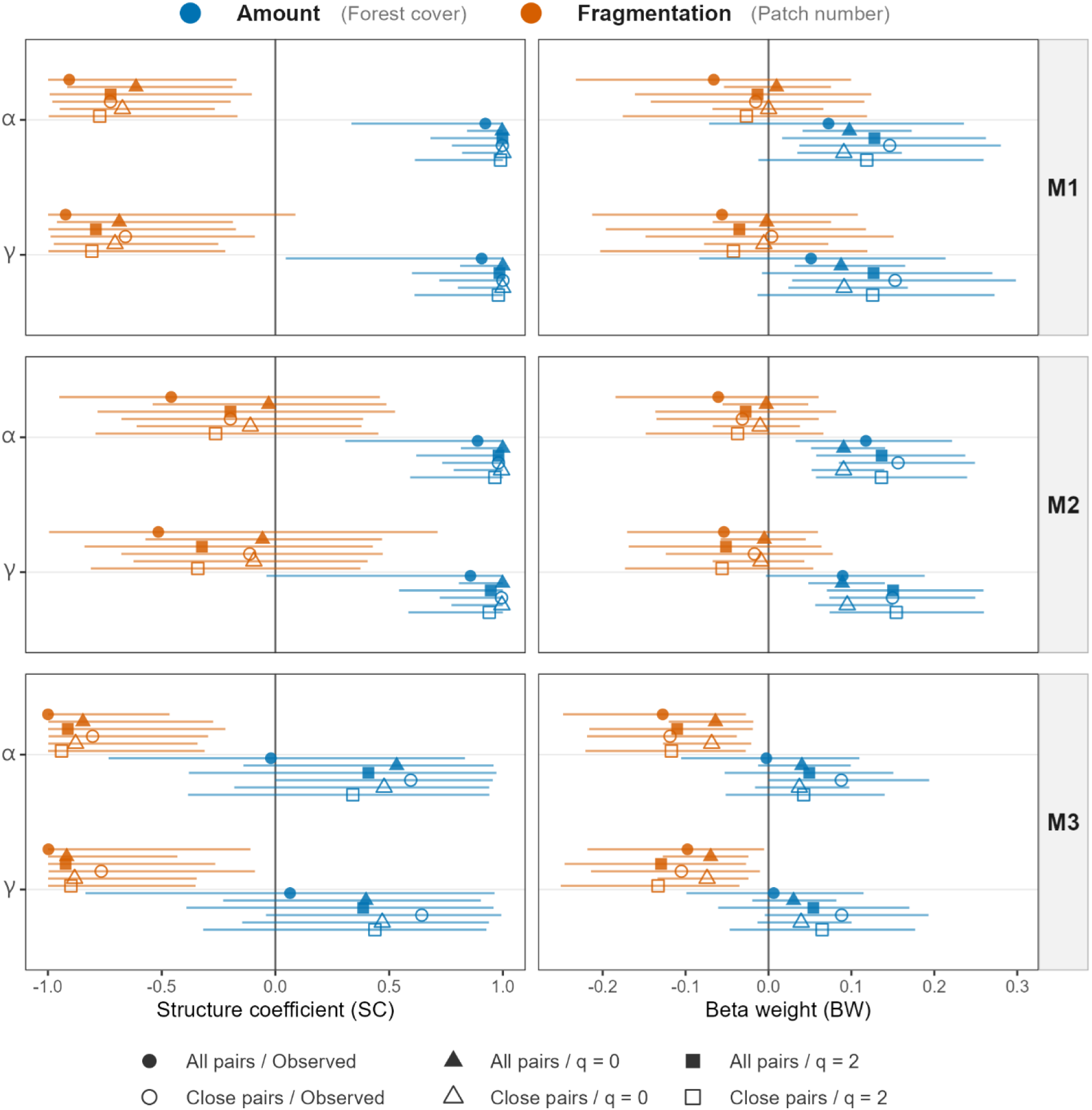
Variance partitioning invariance across 18 analytical variants — M1–M3, α and γ diversity. Each panel shows standardised beta weights (BW; left column) and structure coefficients (SC; right column) for fragmentation (pink) and habitat amount (green) across all six analytical variants within each diversity component (α, γ) and parameterisation (M1–M3). Point shapes encode pairing design and diversity metric (see legend); horizontal lines show 95% bootstrap confidence intervals. The key diagnostic contrasts are: under M1, fragmentation SC is large and negative across all 18 variants while BW is near zero — the suppressor signature is not variant-specific but geometrically invariant; under M3, BW and SC align in sign for fragmentation across all variants, confirming that gradient reallocation is the mechanism rather than a sampling artefact. Habitat amount BW is positive across all variants and parameterisations and decreases under M3 as shared variance is redistributed to the fragmentation predictor. Semi-partial R² and full 18-variant outputs including β diversity in Figs. S2.2–S2.3 (Supplementary Material S2).

### Parameterisation sequence as geometric confirmation

Coefficient patterns shifted systematically across parameterisations in the direction predicted by suppressor geometry (Figs. 2–4; Table 1). Residualising fragmentation on habitat amount (M2) allocated the shared habitat-loss gradient primarily to the amount predictor. The amount coefficient increased, while the fragmentation coefficient remained near zero. The coefficient allocation matched the prediction from the parameterisation sequence: amount BW increased while fragmentation BW remained near zero, and the attenuation was asymmetric (fragmentation near zero, amount inflated) rather than bilateral.

M2 retained residual nonlinear coupling of 0.444 in both diagnostic orientations, above the dataset-specific threshold (GAM R² < 0.02), and did not satisfy the residual-uncoupling criterion. Residualising habitat amount on fragmentation (M3) produced the opposite allocation. Fragmentation coefficients shifted consistently negative for α diversity (median BW = −0.11; all six variants negative) and γ diversity (median BW = −0.10; all six variants negative), while the amount coefficient decreased toward zero (α median BW = +0.04; γ median BW = +0.05). The frag/amount BW ratio shifts from approximately 0.13 in M1 to approximately 2.7 in M3 for α diversity, and from 0.19 to 2.2 for γ diversity — a reversal of apparent predictor importance.

The negative-control parameterisation (M4) confirmed the gradient variance allocation by producing its predicted mirror image: large negative fragmentation BW and near-zero amount BW, opposite to M1. M1 allocated almost all shared-gradient variance to amount; M4 allocated it primarily to fragmentation. M3 was the only parameterisation meeting the dataset-specific residual-uncoupling criterion (Supplementary Material S2 (Fig. S2.4)).

To facilitate direct comparison with the GS25 categorical specification, a GS25-style model (diversity ∼ patch_type + (1|refshort)) was fitted alongside the F26 additive specification M1 (diversity ∼ frag_z + amount_z + (1|refshort)) across all 18 analytical variants; |ΔAICc| < 2 was the threshold for indistinguishability. Within-variant AICc comparisons showed that M1 and M2 were indistinguishable in predictive fit (|ΔAICc| ≤ 2 in 18/18 variants; median |Δ| = 0.30), while M1 and M3 differed only marginally overall (M1 lower in 9 of 18 variants; median |ΔAICc| = 1.91). The dominant source of within-variant variability was the diversity metric rather than the pairing design. Under M3, non-rarefied observed richness yielded median BW = −0.123 (α) and −0.101 (γ), while the rarefaction-standardised Hill number q = 0 yielded −0.066 (α) and −0.072 (γ) — approximately 46% and 29% weaker in absolute magnitude. The suppressor geometry was nonetheless invariant across all 18 variants. Pairing design (all pairs vs. close pairs) contributed negligible variation (≤0.03 BW units; ≤5.4% of the M3 α median). Beta diversity showed no equivalent suppressor structure under M1, which is here interpreted as consistent with the possibility that β-diversity responses are context-dependent in ways that prevent a consistent directional signal under the shared-gradient geometry of this dataset: SC ranged widely including positive values, and BW remained near zero under both M1 and M3. The suppressor diagnosis concerns the amount–configuration partitioning problem for α and γ diversity, where F26’s null coefficient is used to infer ecological independence at the landscape scale.

As the shared amount gradient is removed through residualisation, fragmentation semi-partial R² increases 144-fold from M1 to M3 for α diversity (<0.01% to 0.86%). M3 was designated the primary diagnostic parameterisation on the basis of its near-zero residual coupling in both axis orientations alone, not on the basis of coefficient direction or magnitude — both residualisation directions were evaluated against the same criterion, and only M3 satisfied it. BW estimates under M2 and M3 were virtually identical across the full and ≥50% cover subsets (M2: BW Frag = −0.03 for both subsets in both α and γ; M3: BW Frag = −0.11 in the full dataset and −0.12 in the ≥50% cover subset).

### Robustness of the suppressor geometry

The full suppressor signature was preserved within the subset of landscapes exceeding 50% habitat cover (Table 1, Section B1). Nonlinear coupling was reduced but not eliminated in either diagnostic orientation, and the SC–BW mismatch remained consistent across parameterisations. The linear coupling component strengthened in the subset, with R² increasing from 0.454 in the full dataset to 0.634 in the subset. GAM R² was 0.893, giving a nonlinear component of 0.259. Directional asymmetry was also preserved and amplified. The reverse nonlinear component, amount ∼ s(patches), remained much larger than the forward component, 0.259 versus 0.016. That is a 16.2-fold asymmetry, compared with 7.75-fold in the full dataset.

### Continental sensitivity

Within-landscape habitat amount overlapped strongly between South American and non-South American studies and did not differ significantly in the matched-study comparison (Welch t = 0.76, df = 20.6, p = 0.455; SA mean = 68.7% [38.8–98.4%]; non-SA mean = 63.5% [34.5– 94.3%]). The continental contrast was not driven by systematic differences in the amount of forest remaining in sampled landscapes. Semi-partial R² is not reported for the South American continental subset at the sample sizes of these groups; BW and SC from the mixed-model coefficient workflow constitute the available variance-allocation diagnostics for this subset, and the continental contrast is interpreted as a secondary sensitivity check accordingly.

Subset-specific suppressor diagnostics showed that the M1 SC–BW mismatch persisted within both South American (SC = −0.850, BW = −0.043, 6/6 α variants negative) and non-South American (SC = −0.858, BW = −0.192, 5/6 α variants negative) subsets, with larger mismatch in the pooled dataset than in either group separately. The SC–BW ratio in the non-SA subset (∼4.5×) was substantially lower than in the full dataset (∼51×), consistent with the less pronounced coupling geometry in that continental group.

Within SA, M2 remained negative (6/6 α variants) and the M4 negative control — expected to return near-zero coefficients under double residualisation — also remained consistently negative (median α = −0.053, 6/6 negative). The shared gradient is sufficiently entrenched in the SA predictor space that bidirectional residualisation does not resolve it. Within non-SA, M2 flipped positive (median α = +0.110, 0/6 negative) and M4 collapsed to near-zero (median α = −0.018, 4/6 negative), behaving as the negative control should. The M3 coefficient was five-fold larger in non-SA than SA (median α = −0.541 vs −0.114). The near-zero residual coupling achieved by M3 in the full dataset generalised to both continental subsets under within-group residualisation: M3 bidirectional orthogonality was satisfied for both SA and non-SA. Subset-level M3 coefficients are interpretable as approximately orthogonalised effects within their respective predictor distributions, though within-subset results are not directly comparable in magnitude to full-dataset M3 values. The orthogonality criterion was not universally satisfied across all subsets, with the tropical-only subset (n = 32; reverse residual coupling = 0.032, marginally above threshold) and the Tropical non-SA high-cover subset (n = 7; reverse = 0.193) representing boundary conditions at small n (full outputs in Supplementary Material S2 (Section S2.2)).

Path models were additionally fitted separately for the 24 South American (SA) and 13 non-South American (non-SA) studies, directly paralleling GS25’s own continental sensitivity analysis (Extended Data Fig. 10; GS25). Because the Amount → Fragmentation structural path is estimated separately from the diversity-response component, it is invariant across diversity variants within each continental subset by construction; its coefficient was −0.694 in SA and −0.597 in non-SA. Three further patterns emerged in the diversity-response paths across the full set of variants. First, the Frag → Diversity path was consistently negative in the SA group (6/6 α variants, 6/6 γ variants; medians: α = −0.043 [−0.209, −0.010], γ = −0.095 [−0.210, −0.041]), a directionality more consistent than in the full-dataset analysis. In the non-SA group, the median was more negative for α (−0.201 [−0.544, +0.039], 5/6 negative) but with substantially greater variance, particularly under q = 2 rarefaction (−0.501 all-pairs; −0.544 close-pairs). Second, the direct Amount → Diversity path showed a marked continental contrast: near-zero or negative in SA (median α = +0.087 [−0.084, +0.141]; γ = +0.045 [−0.141, +0.135]) and strongly positive in non-SA (median α = +0.392 [+0.266, +0.526]; γ = +0.520 [+0.284, +0.612]), with several non-SA variants reaching significance (q = 0: α p = 0.023–0.027; γ p = 0.016–0.025). Third, the indirect amount-via-fragmentation effect was positive and more consistent in SA (median α = +0.030, γ = +0.066) than in non-SA (median α = +0.120, γ = +0.045), where larger point estimates were accompanied by large standard errors. Full results are reported in Supplementary Material S2 (Section S2.3).

### Tropical-only subset

Excluding the five temperate studies (as classified in GS25’s Supplementary Table 7) left the design-reconstruction asymmetry essentially unchanged: 26 of 32 remaining study pairs (81.2%) satisfied both directional conditions, compared to 31 of 37 (84%) in the full dataset. M1 returned near-zero fragmentation beta weights in the tropical subset across all six analytical variants (median α = −0.002; median γ = −0.040), matching the full-dataset values (−0.014 and −0.020 respectively), showing the additive null persists within the tropical component of the dataset independently of the temperate studies. M3 recovered negative fragmentation coefficients across all 6/6 variants for both α (median = −0.056) and γ (median = −0.069); because residualisation was rebuilt within the tropical subset, M3 magnitude comparisons against the full dataset should be interpreted cautiously, but the directional pattern is unambiguous. Full diagnostic outputs are in Supplementary Material S2 (Section S2.2.2).

### Path model — causal hierarchy check

The structural path Amount → Fragmentation was strongly negative for the full dataset (standardised coefficient = −0.69; Fig. 5). A competing correlated-errors specification — in which Amount and Fragmentation are treated as exchangeable parallel predictors rather than causally ordered — was strongly disfavoured: the directed model was preferred with ΔAIC = −70.8 and an evidence ratio of 2.4 × 10¹⁵ across the full dataset (Supplementary Material S2, Section S2.3). The direct Fragmentation → Diversity path was near-zero (median α = −0.029, negative in 5/6 variants; median γ = −0.041, negative in 5/6), whereas the direct Amount → Diversity path was strongly positive across all variants (median α = +0.231; median γ = +0.284). Full 18-variant coefficients and R² are in Supplementary Material S2 (Section S2.3).

**Fig. 5.**
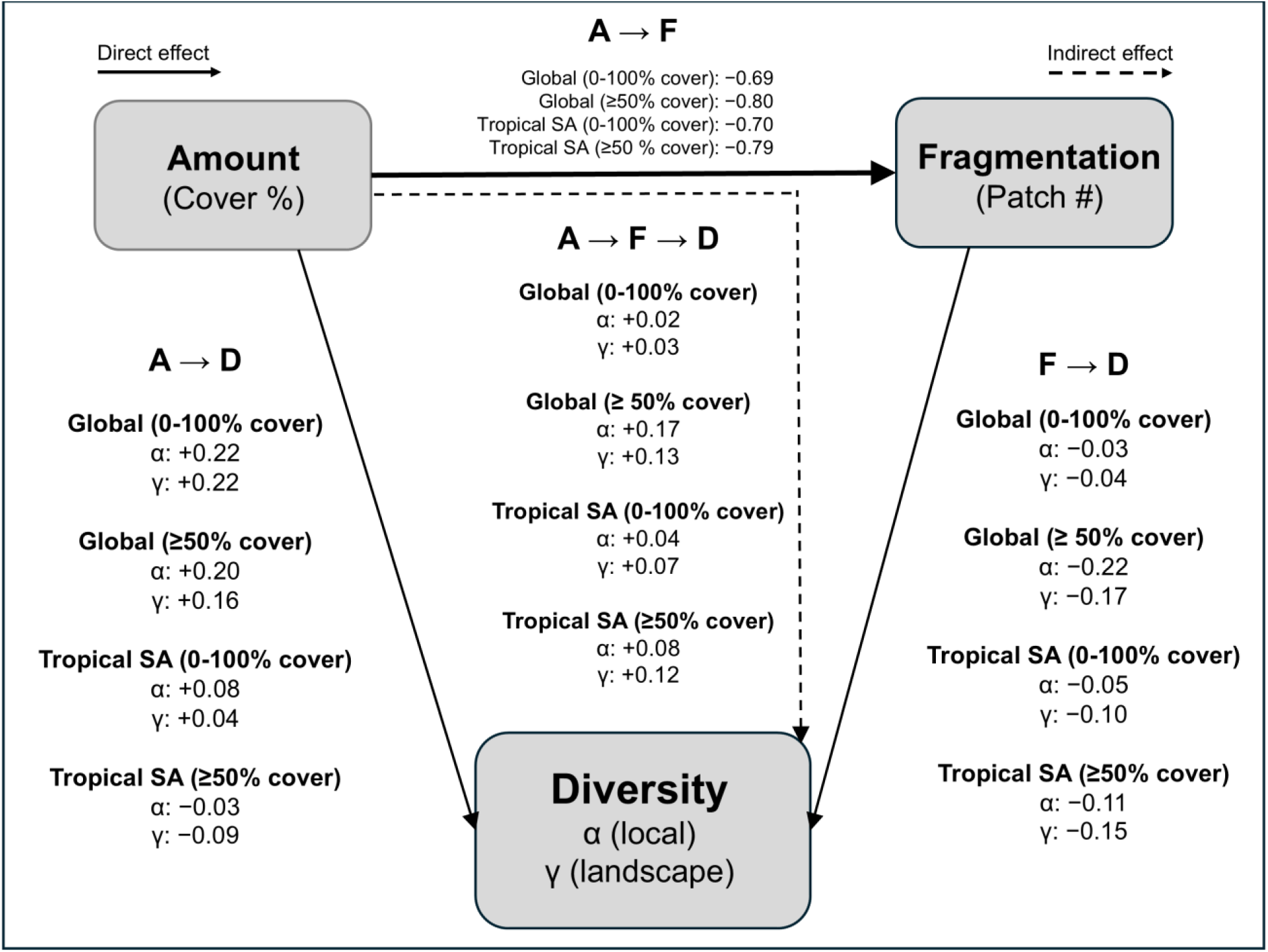
Path model causal hierarchy across four analytical groups. Nodes represent the three observed variables: habitat amount (forest cover within a 2-km buffer), fragmentation (number of patches), and biodiversity (α and γ richness). Coefficients shown on each path are medians across 6 analytical variants (observed richness, q = 0, q = 2, each in all-pairs and close-pairs designs), reported separately for four groups: Global (0--100% cover, n = 37), Global (≥50% cover, n = 17), Tropical SA (0--100% cover, n = 23), and Tropical SA (≥50% cover, n = 9; Tier 3, directional indicators only). The Amount → Fragmentation structural path (solid arrow) is strongly negative in all groups. Direct effects of fragmentation on diversity are negative and increase in magnitude under coverage-restricted subsets. Direct effects of habitat amount on diversity are positive and attenuate in the Tropical SA subsets. Indirect effects of amount on diversity via fragmentation are positive across all groups. The model is saturated (Fisher’s C = 0, df = 0, p = 1.000). β diversity paths are reported in Supplementary Material S2 (Section S2.3). Path coefficients use 1-SD standardisation.

### Crossed tropical sensitivity checks

One of the four crossed tropical cells satisfied the M3 orthogonality criterion at adequate sample size: Tropical SA (n = 23). Two further cells — Tropical non-SA (n = 9) and Tropical SA high-cover (n = 9; Tier 3) — returned 0.000/0.000 residual GAM values, but at that sample size the GAM lacks leverage to detect nonlinear structure; those values reflect estimation failure rather than confirmed orthogonality, and both cells are retained as Tier 3 directional indicators on sample-size grounds. The fourth cell — Tropical non-SA high-cover (n = 7; Tier 3) — failed outright (reverse residual coupling = 0.193). In all four cells, the forward nonlinear component was negligible (r = −0.68 to −0.89 in the forward direction). Within Tropical SA (n = 23), overall predictor coupling exceeded the full-dataset level (M4 GAM R² = 0.805 vs 0.632), consistent with a stronger shared habitat-loss gradient in the Atlantic Forest system.

In tropical South American studies (n = 23), the M1 SC–BW mismatch persisted (SC = −0.869, BW = −0.049, 6/6 α variants negative) and fragmentation coefficients became more negative under M3 (median BW = −0.112, 6/6 negative); the Tropical SA high-cover cell (n = 9; Tier 3, directional indicators only) showed directional consistency (M1 BW = −0.088; M3 BW = −0.060, 6/6 negative) but no intensification relative to the full Tropical SA result. In tropical non-South

American studies (n = 9; directional indicators only), the additive model yielded a positive fragmentation coefficient (M1 BW = +0.289, 0/6 negative) that reversed sign under M3 (BW = −0.205, 6/6 negative). Because the tropical non-South-American cell is thin (n = 9) and semi-partial R² is not reported, these patterns are directional indicators only (full outputs in Supplementary Material S2, Section S2.2.3).

### Model adequacy

Simulation-based residual diagnostics (DHARMa) showed no significant deviations in uniformity, dispersion, or outliers across all representative models. Levene tests for homogeneity of residual variance between fragmented and continuous landscape types returned no significant results in any variant across all four parameterisations (M1–M4). Per-study random intercept distributions were stable across parameterisations, with intercept shifts small in absolute magnitude relative to the random-effect standard deviation (maximum shift ±0.113 vs. SD ≈ 0.867), though moderately correlated with study-mean predictor values in some parameterisations (full values in Supplementary Material S2 (Section S2.4)). Total landscape explanatory power was small across all parameterisations (marginal R² ≈ 0.017 in M1), with study-level random effects absorbing the remainder (conditional R² ≈ 0.94).

## Discussion

The fragmentation debate has persisted in part because related ecological questions have been treated operationally as equivalent (Didham et al. 2012; Haila 2002; Miller-Rushing et al. 2019; Valente et al. 2023). Patch-scale and experimental studies ask whether biodiversity within remnant habitat declines beyond passive-sampling expectations after habitat loss (Chase et al. 2020a). Landscape observational studies such as GS25 and F26 ask whether configuration covaries with biodiversity after controlling for habitat amount. The fragmentation-per-se hypothesis, formally, that configuration exerts an independent causal effect on biodiversity separable from habitat loss, the canonical alternative to the HAH (Fahrig 2003, 2013, 2017), asks something narrower and more demanding than either observational approach, namely whether that separable component can be identified from the predictor geometry a given dataset provides (Fletcher et al. 2023a; Lundberg et al. 2021). These questions are related but not interchangeable. Treating them as equivalent has helped produce the locked demonstration/counter-demonstration cycle Valente et al. (2023) describe, where different study designs can appear to disagree even when they are not identifying the same ecological quantity. The conditions under which the second question can be read as evidence about the third are not assumed in the present study but evaluated. Under the observed predictor geometry, they are not met.

The geometry fails the separability condition for a structural reason that precedes any modelling choice: habitat loss generates configuration change through the subdivision process itself, embedding amount and configuration in a shared habitat-loss gradient with asymmetric nonlinear coupling before any landscape is sampled (Didham et al. 2012; Fletcher et al. 2023a; Koper et al. 2007; Ruffell et al. 2016; Fig. 1a). The attainable predictor space is therefore asymmetric and bounded. The upper-right zone of high amount and many patches is geometrically forbidden by the subdivision identity, leaving the realised sample concentrated at high cover with a thin tail toward intermediate configuration (Fig. 1b). Under this geometry, the additive parameterisation yields the classic cross-over suppressor signature. Fragmentation aligns strongly with the fitted biodiversity gradient yet contributes almost no unique variance once habitat amount is included (Ray-Mukherjee et al. 2014; Smith et al. 2009; Fig. 1c). This produces a near-zero beta weight that reflects predictor architecture rather than ecological independence (Fig. 1d). The near-zero full-dataset coefficient is therefore geometry-conditional. Wherever the geometric constraint is reduced, the recoverable fragmentation-associated signal is consistently and non-trivially negative (Ruffell et al. 2016).

### From predictor coupling to suppressor attenuation

Redundancy and suppression are not the same inferential problem, and the distinction matters for how the near-zero coefficient should be interpreted. Collinearity between predictors generates shared variance that reduces the precision of partial coefficients (Dormann et al. 2013; Graham 2003), a problem familiar from landscape ecology (Koper et al. 2007; Smith et al. 2009). Suppressor attenuation is a geometrically specific subtype of redundancy, whereby the subordinate predictor correlates with the dominant predictor in the direction opposite to its own relationship with the response, so the dominant predictor’s inclusion drives the subordinate’s partial coefficient toward zero while leaving its alignment with the fitted gradient intact (Smith et al. 2009; Ray-Mukherjee et al. 2014). The SC–BW dissociation is the diagnostic fingerprint that distinguishes them. Under redundancy, SC and BW shrink together as shared variance is absorbed (Smith et al. 2009; Prunier et al. 2015); under suppression, SC is disproportionately preserved relative to BW, because the dominant predictor overcorrects in a direction fixed by the sign geometry of the predictor pair.

In this dataset, that dissociation is extreme. Fragmentation’s SC remains large and negative across all 12 α and γ variants in M1 while BW collapses toward zero (Table 1; Fig. 1c). The magnitude tracks the residual coupling across parameterisations. In M1 the SC²/R²sp ratio reaches ∼8,800-fold for α and ∼3,000-fold for γ. Under M2, partial coupling removal reduces these to ∼59-fold and ∼39-fold. Under M3, verified orthogonality collapses them to ∼94-fold and ∼111-fold while SC and BW align in sign, confirming that the residual ratio under M3 reflects genuine signal rather than geometric suppression (Ray-Mukherjee et al. 2014; Prunier et al. 2015). Whether that progression reflects suppression specifically or redundancy more generally is what the following three results address. The process coupling documented here, habitat loss co-generating amount and fragmentation along a directed asymmetric gradient before any landscape is sampled (Fahrig 2003; Didham et al. 2012; Fletcher et al. 2018; Ruffell et al. 2016), is what converts collinearity into suppression. It fixes the correlation sign between predictors and the response direction of fragmentation such that amount’s inclusion systematically deflates fragmentation’s partial coefficient regardless of its true ecological magnitude.

Three results show that this pattern is suppressor attenuation rather than ordinary redundancy. First, M3 shifts fragmentation’s BW uniformly negative across all 12 α and γ analytical variants. Suppressor geometry predicts directional recovery once the shared gradient is reduced; ordinary redundancy does not, because it provides no reason to expect the coefficient to recover in any specific direction after shared variance is reallocated.

Second, the path model provides process-level confirmation. The directed specification (Amount → Fragmentation → Diversity) is preferred over the correlated alternative with an evidence ratio of 2.4 × 10¹⁵, meaning the data overwhelmingly support amount structurally predicting fragmentation rather than the two predictors operating as exchangeable parallel causes. When collinearity has this causal basis, with one predictor structurally generating the other along a directed pathway, multiple regression gives unbiased estimates of direct effects but biased estimates of total effects because the partial fragmentation coefficient absorbs not only shared variance but also the indirect effect of amount operating through configuration. This divergence arises specifically when the conditioning predictor shares nonlinear variance with the predictor of interest (Ruffell et al. 2016). Additive regression cannot isolate those components.

Third, the SC–BW dissociation is present in α and γ diversity but absent in β diversity under M1. That pattern is consistent with the suppressor signature being architecture-specific rather than a universal property of the dataset or the modelling framework. A universal artefact would produce the same SC–BW dissociation across all diversity dimensions equally; its restriction to α and γ is what the suppressor diagnosis predicts and what distinguishes it from a dataset-wide confound. What may account for this restriction is that compositional turnover responds to processes such as distance-decay of similarity, habitat heterogeneity, and matrix permeability that are not reducible to the amount–configuration partial-regression geometry producing suppressor attenuation in the richness components (Püttker et al. 2014; Fletcher 2025). Land-use change alters dispersal, environmental filtering, and biotic interactions in ways that need not affect local assembly and among-site turnover in the same direction or at the same scale (Chase et al. 2020b), so the absence of suppressor structure in β is plausible without requiring a separate explanation specific to that diversity dimension.

The directed asymmetry is what makes M3 diagnostic rather than arbitrary. Habitat loss is the generating process. As cover declines, the remaining habitat subdivides into more and smaller patches, so amount is causally prior to configuration. Patch number cannot exceed what the available area can generate. The reverse dependency has no equivalent mechanism. The nonlinear component is 7.75-fold larger in the reverse orientation (Amount ∼ s(Patches)) than in the forward (Patches ∼ s(Amount)), because amount responds to patch number along the strongly curved percolation surface that habitat loss generates, whereas patch number is only a partial predictor of amount. M3 exploits this directionality. Residualising amount on fragmentation removes the downstream echo of the causal process from the amount predictor, leaving a fragmentation predictor estimated against an amount predictor no longer dominated by the shared gradient (Koper et al. 2007; Graham 2003; Ruffell et al. 2016). M2 does the opposite. Residualising patches on amount removes the causal signal from the predictor that carries it, which is why it fails the orthogonality criterion. Zeroing the linear correlation leaves substantial nonlinear coupling intact in the reverse orientation (residual GAM R² = 0.44), leaving the shared gradient unreduced (Fig. 2). Only M3 satisfies the criterion in both orientations, and only M3 shifts the fragmentation coefficient in the negative direction that suppressor geometry predicts from the sign of its structure coefficient (Ray-Mukherjee et al. 2014). The 7.75-fold asymmetry is the quantitative expression of that directionality, a degree of coupling anisotropy that Koper et al.’s (2007) symmetric framing does not anticipate and that M3 exploits by operating in the causal direction (Figs. 3–4). This coefficient shift occurs with no improvement in predictive fit, ruling out the interpretation that M3 recovers a negative coefficient because it better represents the ecological signal.

The double residualisation of both predictors simultaneously (M4) closes the architectural argument. Under M4, neither predictor dominates the shared gradient. Fragmentation’s SC collapses and its BW is no longer suppressed relative to its SC (Ray-Mukherjee et al. 2014), while amount’s SC retreats and its BW attenuates (Supplementary Material S2, Fig. S2.4), confirming that the gradient dominance of amount in M1 and the suppression of fragmentation are two sides of the same geometric coin, both inexplicable under the ecological independence interpretation (Smith et al. 2009). The suppressor signature that M1 inherits is therefore not a property of fragmentation as a predictor or of the additive specification as a model. It is a property of the asymmetric landscape-level coupling that precedes any modelling choice. A fragmentation predictor functioning solely as a proxy for habitat amount would also produce near-zero partial slopes once amount is included. What it could not produce is the SC–BW mismatch, because a pure proxy’s alignment with the fitted gradient would collapse once the correlated predictor is included. That collapse does not occur here. Under the observed predictor geometry, the F26 null is therefore compatible with suppressor attenuation and cannot be read as an ecological null unless predictor separability is first demonstrated (Freckleton 2002). In this dataset, it is not (Fig. 1c–d).

The suppressor geometry is not an artefact of the diversity model. It is already present in the predictor space before any response variable is introduced. When landscape type itself is regressed onto the two F26 continuous predictors, the same SC–BW–R²sp mismatch reappears, large structure coefficients, near-zero beta weight, and near-zero semi-partial R² for the patch predictor, confirming that the identification problem is structural rather than specific to a particular biodiversity response or model specification (Supplementary Material S1, Section S1.1.1). The M1 suppressor signature is therefore the regression-level consequence of a shared predictor-space geometry, present across landscape classification, continuous predictor space, and diversity regression alike, rather than evidence that configuration is ecologically unimportant.

The directed coupling asymmetry is itself ecologically interpretable, not a statistical property of the predictor cloud but a reflection of the process that generates landscapes. Habitat loss at high cover initially subdivides habitat into more patches, whereas continued loss at low cover increasingly removes patches altogether, producing a hump-shaped amount–patch number relationship that is asymmetric: the rise is steep as continuous habitat begins fragmenting, the descent is gradual as patches coalesce at low cover (Fletcher et al. 2023a; Wang et al. 2014). This threshold pattern is documented in tropical fragmented landscapes where the cover–edge relationship reverses sign near 30% forest cover (Andrén 1994; Hazard et al. 2026), and the 7.75-fold asymmetry of the nonlinear coupling component documented here is its statistical expression. The ecological-conditionality argument, including trait-level fragmentation sensitivity, matrix quality as a moderating factor, and the compounding of predictor-geometry and context-dependence ambiguities in multi-study syntheses, is developed fully in Supplementary Material S2 (Section S2.2.5).

### What does the additive null identify?

In this dataset, an estimable coefficient and an identified ecological effect are not the same thing. The additive specification used by F26 yields statistically valid partial coefficients — the question is whether those coefficients identify the independent ecological effect the analysis is intended to test (Freckleton 2002; Lundberg et al. 2021). A coefficient identifies an independent ecological effect only when the predictor geometry provides the contrast the estimand requires in the fragmentation-per-se case, landscapes that vary in configuration while holding habitat amount constant. That contrast is structurally scarce in this dataset (Fig. 1b; Supplementary Material S1, Section S1.1), which means the additive coefficient estimates a quantity whose ecological interpretation differs from the hypothesis being tested, regardless of the statistical validity of the estimation procedure.

F26 acknowledged directly that their primary causal interpretation, that the GS25 signal originated from forest amount differences at the broader landscape scale, could not be evaluated with the available data (Fahrig et al. 2026), placing that interpretation outside the empirical scope of their reanalysis. The identification gap is not incidental to this dataset but structural: the design-reconstruction analysis confirms that same-amount, different-configuration landscape pairs are structurally scarce throughout the predictor space, a scarcity that persists even within the highest habitat-amount stratum (77–100%), where the nonlinear coupling is steepest (Supplementary Material S1, Section S1.1; Martínez-Lanfranco 2026; Fig. 1b). The upper-right zone of the attainable predictor space — high habitat amount, many patches — is geometrically forbidden by the subdivision identity, so the contrast the fragmentation-per-se hypothesis requires is not merely undersampled but unattainable under habitat-loss-generated landscape gradients.

The configuration coefficient F26 report estimates the partial association between patch number and biodiversity conditional on the shared habitat-loss gradient (Koper et al. 2007; Smith et al. 2009), not the effect of configuration independent of habitat loss, which is the ecological quantity the fragmentation-per-se inference requires (Fahrig 2003; Fletcher et al. 2023a; Lundberg et al. 2021). The M1 fragmentation coefficient removes the component of patch number that lies on the linear projection of forest cover, then estimates the residual association. But that residual is not patch number with habitat loss held constant, because the nonlinear coupling means the shared gradient is not eliminated by linear conditioning (Freckleton 2002; Koper et al. 2007; Fahrig 2003; Didham et al. 2012; Fletcher et al. 2018; Ruffell et al. 2016; Supplementary Material S1, Section S1.1; Martínez-Lanfranco 2026). The model therefore estimates a quantity whose ecological interpretation differs from the hypothesis being tested.

Additive habitat-amount control within a shared observational gradient cannot, by itself, be taken as a rigorous test of the habitat amount hypothesis unless the framework’s own separability conditions are first demonstrated in the realised data (Fahrig 2013, 2015; Fahrig et al. 2019; Hanski 2015). Statistical adjustments for collinearity remove statistical dependence between predictors but cannot dissolve the underlying functional correlation. Absent experimental manipulation, no adjustment technique can establish whether the variables are truly separable as ecological causes (Graham 2003). The shift from GS25’s random-slope treatment of forest amount to F26’s fixed-effect additive control does not resolve the separability problem. It changes the precision of the amount coefficient estimate, not the geometry of the predictor space.

Partial coefficients remain unbiased under high collinearity when habitat amount and fragmentation are manipulated independently, producing approximately orthogonal predictors — Smith et al. (2009) established this under simulated landscapes below 30% habitat cover, where the amount–fragmentation relationship is approximately linear and the nonlinear coupling that dominates higher cover values has not yet emerged. F26 cite that framework to justify the additive model, but this dataset, with a median cover of 70% spanning the full gradient, occupies the nonlinear regime that simulation did not examine and whose conditions it therefore does not validate. The same orthogonality condition underlies early models in which loss and fragmentation were independently assigned by design (Fahrig 1997, 1998, 2001, 2002). Those results are ecologically valid precisely because orthogonality was built in by construction rather than assumed from observational data.

A coefficient can be statistically valid as an estimator while its estimand does not correspond to the independent ecological effect the analysis is intended to measure (Freckleton 2002; Lundberg et al. 2021). Ruffell et al. (2016) formalise this distinction by showing that multiple regression gives unbiased estimates of direct effects but biased estimates of total effects when indirect pathways exist. This divergence arises specifically when the conditioning predictor shares nonlinear variance with the predictor of interest. At the most fundamental level, the allocation is deterministic. When fragmentation is residualised on habitat amount, the fragmentation index retains only the variance unique to fragmentation, while the habitat amount index retains both its own unique variance and all joint variance shared with fragmentation. The latter is entirely absorbed by the unresidualised predictor regardless of its true ecological source (Koper et al. 2007). That this geometry went undiagnosed for so long is itself informative: Smith et al. (2009) reviewed 33 landscape fragmentation studies spanning six statistical methods and found no study that had reported or tested for suppressor relationships, meaning the field had been producing near-zero configuration coefficients under conditions where suppressor attenuation was a predictable and undetected output of the analytical architecture.

The asymmetry of the attenuation follows directly from the sign configuration of the predictors in this system. Habitat amount has a positive bivariate association with biodiversity, fragmentation has a negative association, and the two predictors are themselves negatively correlated, so the partial regression formula adds a positive contamination term to an already-negative fragmentation coefficient, pushing it toward zero. Because amount dominates the fitted gradient, the cross-contamination from amount into fragmentation is large, whereas the reverse contamination is smaller. When amount does not dominate the fitted gradient to the same degree, as in the non-SA subset, where the nonlinear coupling is less pronounced and both predictors retain larger partial slopes, the contamination term is correspondingly smaller, the SC–BW ratio is lower (∼4× versus ∼51× in the full dataset), and the suppressor is less severe without being absent. The SC–BW ratio is therefore the appropriate diagnostic readout: it measures how much of the predictor’s alignment with the fitted biodiversity gradient survives the conditioning on amount, and it does so without reference to statistical significance, model fit, or sample size (Ray-Mukherjee et al. 2014; Prunier et al. 2015). A large SC with a near-zero BW signals that the ecological signal is present in the fitted gradient but geometrically unavailable to the partial coefficient once the shared gradient has been absorbed.

The suppressor diagnostic rests on the SC–BW mismatch in M1, large negative structure coefficients alongside near-zero partial slopes, not on the statistical significance of the M3 fragmentation coefficient (Ray-Mukherjee et al. 2014; Smith et al. 2009). The habitat amount hypothesis remains “just a hypothesis” until evaluated against empirical data under the design conditions it requires (Fahrig 2015); a null additive coefficient under unverified predictor geometry does not constitute such a test (Freckleton 2002). A predictor that contributes no information to the fitted gradient would produce both near-zero structure coefficients and near-zero partial slopes (Ray-Mukherjee et al. 2014). In M1, the first condition does not hold for fragmentation. SC remains large and negative while BW collapses, which is the suppressor signature rather than a null predictor signature. The tight confidence intervals around F26’s near-zero fragmentation coefficients reflect the stability of the suppressor constraint across model evaluations, not ecological precision. Under cross-over suppressor attenuation, the beta weight is driven toward zero by the geometric architecture regardless of the true ecological signal, so interval width measures geometric stability rather than effect absence.

Accordingly, the M1 fragmentation coefficient is a conditional association along a shared habitat-loss gradient, not an estimate of fragmentation per se. M3 is the least confounded quantity this dataset supports, but it is not a definitive estimate of fragmentation per se either. Three limits prevent that reading. First, the residual amount predictor in M3 retains the nonlinear component of the amount–fragmentation relationship. The conditioning is partial, not complete, because GAM residualisation removes the component of amount predictable from patches but the equal-amount contrast the estimand requires is geometrically foreclosed regardless of how amount is residualised (Freckleton 2002; Lundberg et al. 2021). Second, the equal-amount, different-configuration contrast that fragmentation per se requires is geometrically foreclosed in this dataset. The predictor space does not contain it, and no reparameterisation of M1 through M3 recovers it (Fahrig 2003; Ruffell et al. 2016; Fletcher et al. 2018; Fig. 1b; Supplementary Material S1, Section S1.1; Martínez-Lanfranco 2026). Third, even a geometrically clean coefficient averages over ecologically heterogeneous study systems (differing in trait composition, matrix permeability, and fragmentation history), so the pooled estimate conflates contexts in which fragmentation effects are expected to be strong with those where they are not (Ewers and Didham 2006; Villard and Metzger 2014). M3 fixes the geometry but not the pooling problem. Thus, M3 is diagnostic rather than definitive. It shows that when the dominant coupling direction is removed, the recoverable signal is consistently negative across all 12 α and γ variants, identifying the direction of suppression and ruling out a stable ecological null, without estimating the magnitude of an independent fragmentation effect (Ray-Mukherjee et al. 2014; Fletcher et al. 2018; Haddad et al. 2017; Koper et al. 2007).

### What the data can and cannot establish

Three interpretations F26 draw from their null result, that fragmentation has no independent effect once habitat amount is controlled, that the positive amount coefficient identifies the primary ecological driver, and that any prior categorical signal reflected uncontrolled amount variation (Fahrig et al. 2026), each presuppose that additive parameterisation achieved predictor separability in the realised landscape sample, because only under that condition do partial coefficients map onto distinct ecological contrasts (Freckleton 2002). That prerequisite is not met in this dataset: the SC–BW mismatch in M1, the 7.75× nonlinear coupling asymmetry in the Amount ∼ Fragmentation orientation, and the design-reconstruction result that 84% of study pairs co-vary directionally in both predictors jointly indicate that the additive coefficients remain constrained by shared predictor geometry, so their ecological interpretation depends on demonstrated separability in the realised sample rather than on model form alone (Table 1; Figs. 2–4; Supplementary Material S1, Section S1.1; Martínez-Lanfranco 2026). That separability failure is not the only problem for the null. Wherever the geometric constraint on the predictors is reduced, whether by cover stratification, continental subsetting, or residual uncoupling, the fragmentation coefficient moves in the same negative direction rather than remaining stably near zero (Ray-Mukherjee et al. 2014; Smith et al. 2009). A stable ecological null predicts stability near zero regardless of which landscapes are sampled or which parameterisation is applied; the F26 model does not return that stability in this dataset.

Near-zero fragmentation coefficients in the full dataset sit inside a suppressor signature and therefore cannot be read as ecologically empty. Four lines of evidence converge on a directionally negative fragmentation-associated structure: large negative M1 structure coefficients for fragmentation before any reparameterisation (Table 1; Fig. 2); stabilisation of the raw M1 coefficient to consistently negative across all twelve α and γ variants in the ≥50% cover subset (Table 1; Supplementary Material S2 (Section S2.2)); consistently negative M3 coefficients under verified residual uncoupling (Table 1); and negative Fragmentation → Diversity paths in all four path-model subgroups (Figs. 5–6; Supplementary Material S2 (Section S2.3)). No single result identifies a definitive independent fragmentation-per-se effect size. In this dataset, a stable ecological-null interpretation is less defensible than a geometry-conditional one. The full-dataset near-zero coefficient is non-diagnostic under suppressor geometry; the underlying signal is not zero. Fragmentation aligns strongly with the fitted diversity gradient (SC = −0.725 for α) while contributing almost no unique variance, which is the suppressor-attenuation pattern, not the zero-signal pattern. Wherever the geometric constraint is reduced, the coefficient is consistently and non-trivially negative. A stable ecological null must account for all of these results simultaneously with no coherent mechanism; the directional interpretation accounts for all of them under one: a negative fragmentation-associated signal geometrically attenuated under the full-dataset additive specification. This interpretation does not require the GS25 categorical signal to be dismissed. GS25 detected lower α and γ diversity in fragmented landscapes using a categorical contrast that encodes fragmentation as a process-level difference between landscape types, a signal consistent with the directional pattern documented here (Fahrig et al. 2026; Fletcher et al. 2018; Fletcher et al. 2023a; McGarigal and Cushman 2002).

**Fig. 6.**
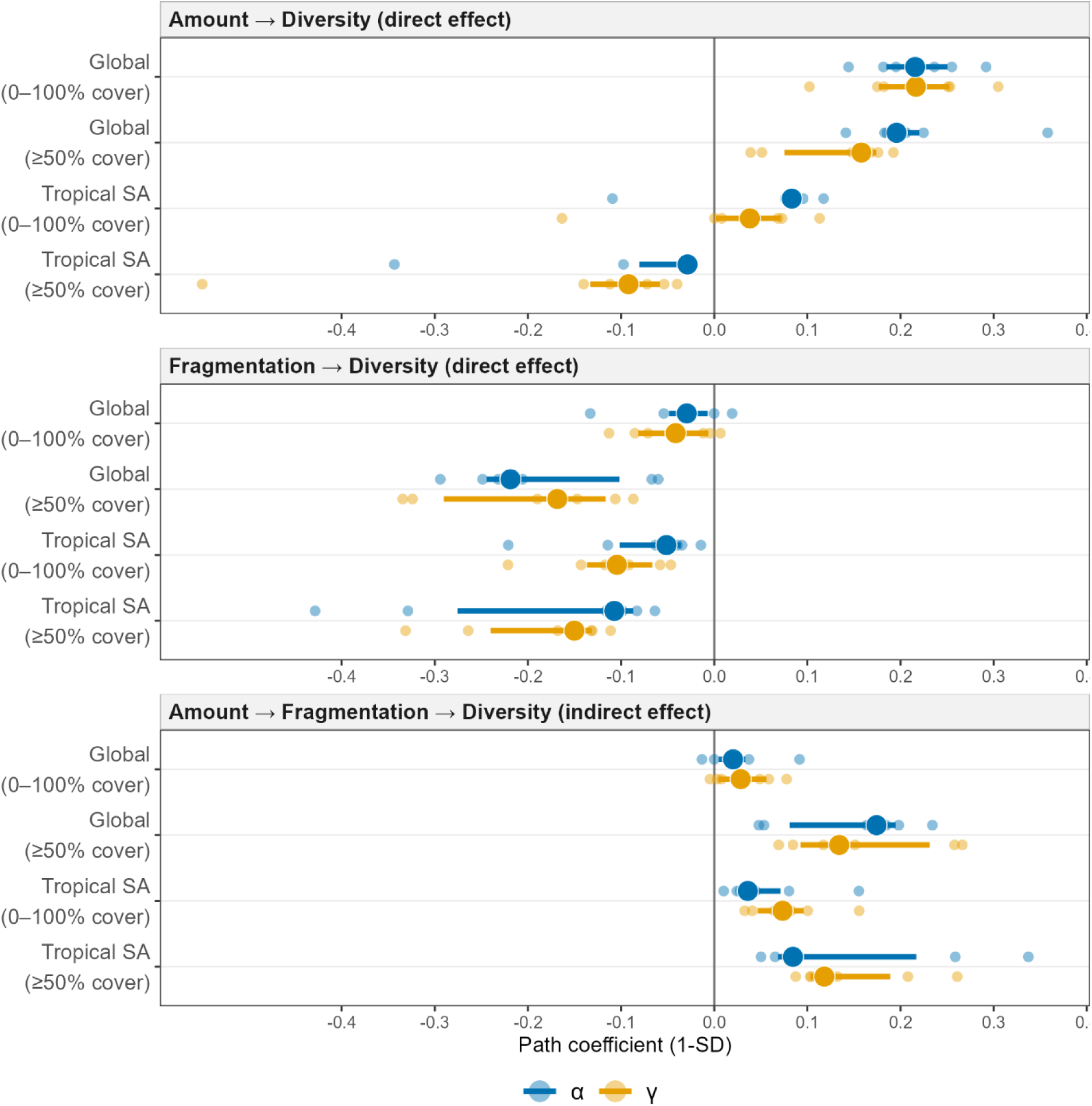
Path model estimates across analytical groups — direct and indirect effects on diversity. Each panel shows one estimable quantity from the directed SEM: the direct effect of habitat amount on diversity (top), the direct effect of fragmentation on diversity (middle), and the indirect effect of amount mediated through fragmentation (bottom). Groups are on the y-axis; large filled circles show medians across 6 analytical variants (observed richness, q = 0, q = 2 × all-pairs and close-pairs); horizontal lines show interquartile ranges; small translucent points show individual variant estimates. Blue = α diversity; orange = γ diversity. The four groups represent successive constraint on the predictor distribution: Global (0--100% cover, n = 37), Global (≥50% cover, n = 17), Tropical SA (0--100% cover, n = 23), and Tropical SA (≥50% cover, n = 9; Tier 3, directional indicators only). Fragmentation direct effects are negative across all four groups and both diversity components, with effect magnitude increasing under coverage-restricted subsets. Indirect effects are positive across all groups. Path coefficients use 1-SD standardisation and are not directly comparable in scale to the beta weights (use 2-SD) in Figs. 3–4.

In the ≥50% cover subset, the same raw F26-style additive specification, with identical model form and identical predictor definitions, produces consistently negative fragmentation coefficients across all 12 α and γ diversity variants, while the full-dataset specification returns near-zero (Table 1; Supplementary Material S2, Section S2.2). This shift is not produced by a relaxation of predictor coupling — the reverse-dominant nonlinear asymmetry is more pronounced in the high-cover subset (16.2× vs 7.8×) and overall coupling strength is higher (Fig. S2.5 vs Fig. S2.1). Two coefficient patterns accompany the sign recovery: amount’s partial coefficient is smaller in the high-cover subset and fragmentation’s structure coefficient is more negative (Table 1; Supplementary Material S2, Section S2.2). These patterns are consistent with two non-exclusive interpretations that the continental comparison below helps evaluate. What the data establish unconditionally is that the sign recovery occurs under stronger, not weaker, geometric coupling. This rules out coupling relaxation as the explanation and confirms that the full-dataset near-zero coefficient is geometry-conditional rather than ecologically stable. A stable ecological null predicts stability near zero regardless of which landscapes are sampled. The F26 model does not return that stability.

The path model provides a complementary structural check, not an independent test of effect size. The Amount → Fragmentation structural path was strongly negative and invariant across all analytical groups (Supplementary Material S2, Section S2.3), supporting a directed causal hierarchy consistent with amount generating fragmentation rather than the reverse. This is the process-level form of the coupling asymmetry documented in the predictor geometry. The near-zero direct fragmentation path is the expected geometric consequence of retaining the shared gradient. It is the same phenomenon as the near-zero M1 beta weight, viewed from a complementary causal angle. M3 then shows what happens when that shared gradient is removed. Fragmentation coefficients shift consistently negative across all 12 α and γ variants, completing the inferential arc from suppressor diagnosis through causal structure to signal recovery (Figs. 5–6; Supplementary Material S2, Section S2.3). It is the F26 null coefficient, not the negative direction, that requires a statistical explanation, and that explanation is the suppressor geometry documented here.

A power limitation applies to all subset analyses reported here and to the path model response paths. With n = 74 landscape pairs from 37 studies in the full dataset, the study has limited power to detect fragmentation effects in the ecologically relevant range. A true negative effect would return significance in fewer than one in four replications under the observed standard errors (Supplementary Material S2, Section S2.4). Subset analyses reduce n further, widening standard errors and lowering power proportionally. Coefficient directions and magnitudes in subset analyses should therefore be treated as directional indicators rather than precise effect estimates; statistical non-significance in any subset does not establish ecological absence (Popovic et al. 2024).

This is why M3 remains diagnostic rather than definitive. The three limits established above apply here as well: partial conditioning, a geometrically foreclosed equal-amount contrast, and a pooling problem that persists after the geometry is fixed. M3 additionally estimates only the direct association of patch number with biodiversity given residualised amount — not the total ecological effect of fragmentation, which under causal coupling includes indirect pathways through which configuration change accompanies and compounds habitat loss (Ruffell et al. 2016). The near-zero direct fragmentation path in the path model is subject to the same constraint. The geometric suppression that attenuates the M1 coefficient also attenuates the direct path in the structural model, so a near-zero direct path does not establish that fragmentation effects are ecologically negligible. Additive habitat-amount control within this shared observational gradient is necessary but not sufficient for that conclusion (Fahrig 2013, 2015; Fahrig et al. 2019; Hanski 2015).

The additive null result cannot distinguish ecological absence from suppressor attenuation under the observed predictor geometry, regardless of parameterisation. This is a general property of observational inference (Shaffer and Johnson 2008; Sagarin and Pauchard 2010). Causal attribution from non-experimental data requires multiple converging lines of evidence and explicit acknowledgement of what the design can and cannot rule out, rather than confidence in any one analysis (Schrodt et al. 2025). F26 provides a particularly clear instance of this limit. Null fragmentation coefficients are exactly what shared-gradient geometry predicts even when configuration remains ecologically implicated. A redesign that reproduces rather than resolves that geometry cannot resolve the ecological question it is invoked to address (Koper et al. 2007; Ruffell et al. 2016). The stronger the claim being made, here that this dataset supports fragmentation’s ecological irrelevance after habitat-amount control, the stronger the required demonstration of separability and estimand validity (Freckleton 2002; Lundberg et al. 2021).

A near-zero additive coefficient cannot be read as evidence that biodiversity loss has been ecologically compensated, because compensatory turnover can preserve aggregate community metrics while masking strong species-level change (Ernest and Brown 2001; Supp and Ernest 2014; Russildi et al. 2016; Matthews et al. 2014; Banks-Leite et al. 2014; Chetcuti et al. 2020). In a direct fragmentation-context demonstration, predictor separation was achieved by design (r = 0.22 between patch size and forest amount across 202 forest patches), and interior forest specialists declined significantly with decreasing patch size while total species richness remained flat (Valente and Betts 2018). That is precisely the trait-stratified masking that the pooled α and γ metrics in GS25/F26 cannot resolve: the ecological signal is present in the community but invisible to the aggregate diversity index used to test for it. A parallel result from woody plants in the Dry Chaco, where amount and fragmentation per se were likewise weakly correlated by design (r ≈ 0.2), found that habitat amount predicted species richness while fragmentation per se had a stronger effect on species composition and functional traits. Aggregate richness can support the habitat amount hypothesis while understating configuration effects on community structure (Herrero-Jáuregui et al. 2022).

The convergence prediction provides an explicit falsifiability condition: the suppressor diagnosis would be refuted if M1 and M3 coefficients converged in a dataset where residual coupling between amount and configuration metrics is near zero by design. For example, in experimental fragmentation studies where configuration is manipulated at constant habitat amount (Fletcher et al. 2023a,b; Loke et al. 2019). In datasets where configuration varies primarily at constant habitat amount (the isolating condition identified by Fletcher et al. 2023a) or where configuration metrics reflect connectivity rather than subdivision (Villard and Metzger 2014), the structural driver asymmetry is expected to reverse or disappear, and additive coefficients should become interpretable. A null result at one fixed scale cannot establish scale-general ecological absence.

The framework’s own formulations require contrasts in which habitat amount and local patch attributes can be distinguished rather than co-inherited from the same habitat-loss gradient. When correlation between these predictors is high, landscapes should be selected to minimise that correlation across the sample (Fahrig 2013, 2015; Fahrig et al. 2019; Hanski 2015; Riva et al. 2024b).

Within both continental subsets, M3 residualisation reduces residual coupling to near zero in both axis orientations, and fragmentation coefficients remain consistently negative. This is the paper’s closest approach to the separability condition required for independent fragmentation-per-se inference, in the sub-distributions where that signal should be most recoverable. In tropical South America, M3 additionally reverses the amount predictor’s alignment with diversity from strongly positive under M1 to near-zero, showing that once the shared habitat-loss gradient is dissolved, the residual diversity variation in this system is configuration-dominated rather than amount-dominated (Table 1; full subgroup diagnostics in Supplementary Material S2 (Section S2.2.3)).

Habitat loss generates configuration change rather than the reverse. As a result, the isolating condition that fragmentation-per-se inference requires, configuration varying at constant habitat amount, is structurally rare in landscapes shaped by progressive habitat loss. This is consistent with the reverse-dominant nonlinear coupling in the full dataset, its amplification in the ≥50% cover subset, and the stronger overall predictor coupling in Tropical South America. The rarity is a property of the landscape-generating process, not a failure of any particular analysis. That rarity does not imply that habitat loss must increase fragmentation monotonically in every landscape; at global extent, forest loss has been associated with both increases and decreases in fragmentation depending on fragmentation measure, landscape size, biome, and initial forest amount (Martin et al. 2026). In real forest landscapes, this structural coupling is reinforced by anthropogenic land-use processes. Fragmentation has been increasing in the majority of tropical forests, driven primarily by shifting agriculture and forestry rather than by neutral geometric variation (Zou et al. 2025), reinforcing that amount and configuration are co-generated rather than independently varying in most observed landscapes. This broader entanglement is also evident at large spatial extents — across terrestrial ecoregions, habitat-loss and fragmentation threats are strongly positively correlated, and although habitat loss dominates on average, fragmentation can contribute substantially in severely fragmented systems (Kuipers et al. 2021).

The subset robustness analysis confirms that this structural rarity is not contingent on the full nonlinear coupling strength. Suppressor attenuation can persist through shared linear gradient structure even when nonlinear curvature is reduced. Nonlinearity amplifies the coupling problem but is not its necessary condition (Koper et al. 2007). Additive regression has no mechanism for representing the directional asymmetry that generates this coupling (Koper et al. 2007; Ruffell et al. 2016), it allocates shared gradient variance between coefficient terms without regard for the causal direction of the landscape-generating process. The path hierarchy, M3 orthogonalised coefficients, and subset geometry all converge on the same inferential conclusion in this dataset.

### Context-dependence and ecological heterogeneity

The continental sensitivity analysis shows that the direction and relative magnitude of diversity response paths differ between the South American and non-South American study groups in a way that is consistent with differences in fragmentation history and matrix permeability. In the South American group, direct amount effects are near-zero or negative once fragmentation is included while the Fragmentation → Diversity path remains consistently negative, whereas in non-South America the amount effect dominates and the fragmentation path is secondary (Supplementary Material S2, Section S2.3). These contrasts are directional indicators rather than precise effect estimates: subgroup sample sizes (n = 13–24 studies) yield wide response-path standard errors, and the power to detect effects in the ecologically relevant range is substantially lower than in the full dataset. The directional reversal, not merely a difference in magnitude, is nonetheless consistent with a hard-matrix, long-fragmented system where dispersal mortality, site fidelity, and short movement distances make configuration the operative driver once the shared habitat-loss gradient is removed (Püttker et al. 2020). The continental contrast is not explained by within-landscape habitat cover, which was statistically indistinguishable between groups.

The biome contrast within this dataset illustrates this directly. The GS25–F26 dataset is dominated by Atlantic Forest studies — a tropical biome where recent and intense deforestation has produced assemblages with high proportions of range-restricted specialists that Henle et al. (2004) identify as carrying the sensitivity profile most responsive to spatial configuration, while temperate forest bird assemblages in continental North America show fragmentation effects modulated by land-use history rather than biome-level compositional filtering (Martin et al. 2025b). Those contrasts matter because they show how pooled additive coefficients can flatten ecologically meaningful response variation across systems with different fragmentation histories and species pools. Context-specificity and geometric indeterminacy are independent sources of ambiguity that compound in multi-study syntheses spanning heterogeneous biomes and land-use histories (Didham et al. 2012; Valente et al. 2023; Semper-Pascual et al. 2021).

Globally, fragmentation-sensitive forest-core species are more prevalent in low-disturbance regions, consistent with stronger recoverable configuration sensitivity in Atlantic Forest assemblages than in the non-South-American subset, where repeated disturbance may already have filtered the most sensitive species (Banks-Leite et al. 2014; Betts et al. 2019). More broadly, fragmentation effects can be masked rather than absent: time lags, trait-filtered species loss, and variation in matrix permeability alter which species remain available to express configuration sensitivity in any given system, even before the predictor-geometry problem is considered (Ewers and Didham 2006; Henle et al. 2004; Banks-Leite et al. 2014). Trait filtering tends to favour generalists over specialists, producing biotic homogenisation in which aggregate richness appears stable or even increases while ecologically meaningful composition degrades (McKinney and Lockwood 1999).

Ecological heterogeneity across study contexts constitutes a second source of ambiguity that operates independently of the predictor-geometry problem, pooling responses across systems that differ in fragmentation history, matrix hardness, and species pool composition compounds rather than merely obscures both problems simultaneously, especially when configuration is a meaningful ecological constraint in some contexts but not others (Andrén 1994; Henle et al. 2004). Averaging context-specific responses against near-zero or opposing ones from other contexts produces a pooled estimate that cannot resolve whether fragmentation is ecologically meaningful in any particular system, even in the absence of the predictor-geometry problem (Rybicki et al. 2020; Zhang et al. 2024). Remnant species richness declines more steeply where matrix quality is poorer and surrounding forest cover is lower, and less hostile matrices with greater nearby tree cover can buffer extirpation from even small forest remnants, with the strongest effects in forest-dependent species (Reider et al. 2018; Bueno et al. 2026). Configuration effects should be most recoverable when functional reachability and connectivity constrain persistence over particular portions of the habitat-amount gradient rather than uniformly across all landscapes (Villard and Metzger 2014). This second layer of ambiguity has been repeatedly identified in fragmentation research, where matrix contrast, response traits, trophic structure, lagged responses, and biodiversity dimension can all modulate whether configuration effects are detectable even when richness shows little net change (Ewers and Didham 2006; Wilson et al. 2016; Henle et al. 2004).

Across all the contextual variation documented above, the Amount → Fragmentation backbone remained near-invariant across all subsets and continental strata (Supplementary Material S2, Sections S2.2–S2.3). Cover domain and biome context modulate how strongly the shared gradient attenuates the fragmentation-associated signal, not whether habitat loss generates fragmentation. The ≥50% cover result and the continental contrasts therefore measure variation in biodiversity response expression under a stable structural hierarchy rather than oscillation in the generating mechanism itself. That stability is what makes the subset patterns informative rather than merely consistent, and what grounds the inferential limits documented in this section in a single coherent geometric framework rather than a collection of dataset-specific observations.

### Structural mismatch between data, metrics, and inferential target

The configuration metrics most often preferred in fragmentation research, including patch number, edge density, patch density, perimeter–area ratio, and mean patch size, are also the class-level metrics most likely to be structurally or empirically coupled to habitat amount in habitat-loss landscapes. That coupling is not incidental. It follows from the subdivision identity and its metric-specific variants. Most of these metrics respond jointly to area and aggregation rather than aggregation alone, whereas only a minority are predominantly aggregation-responsive and relatively area-independent (Neel et al. 2004). Several metrics previously treated as weakly abundance-dependent in fact co-vary with both abundance and spatial aggregation across the full cover gradient, and they function most cleanly only within specific abundance windows (Wang et al. 2014). Patch density performs most cleanly at roughly 10–30% cover, and edge density at roughly 30–70%. Outside those windows, the metrics increasingly track abundance as much as or more than configuration.

The entanglement also runs through the apparent area signal. In Atlantic Forest bird communities, detectable area effects disappear once edge effects are controlled (Banks-Leite et al. 2010), meaning that some of the variance attributed to habitat amount in additive models reflects configuration responses at patch boundaries rather than independent habitat extent. Subsequent analysis confirms that complex composite indices designed to jointly represent fragmentation, edge, and habitat quality reduce upon decomposition to being predominantly habitat-amount proxies (Martin et al. 2025a), and the recommended solution (substituting simpler, direct fragmentation measures such as patch density or patch number) does not dissolve the underlying problem. Both metrics are directly constrained by the subdivision identity (Al = Āₚ × Nₚ; Fletcher et al. 2023a): a landscape cannot simultaneously maximise patch number and total habitat area because raising patch count at fixed geometry requires smaller patches, which bounds the area any given patch density can represent. The consequence is visible in Martin et al.’s own empirical data. Patch density is unimodally coupled to forest amount across the landscape gradient, rising to a peak at intermediate cover then collapsing toward zero at both extremes (Martin et al. 2025a), reproducing in a different dataset the exact predictor geometry that Wang et al. (2014) documented for this metric class and that generates suppressor attenuation in additive models. The inferential problem diagnosed here is therefore not one of metric complexity but of geometric separability: subdivision-based metrics are the class of fragmentation indices most directly constrained by habitat amount by construction.

The data are generated by progressive habitat loss, which co-produces amount and configuration before sampling begins. The model assumes additive separability of predictors, and the inferential target requires the same-amount, different-configuration contrast that such landscapes rarely populate. Linear collinearity diagnostics address only the linear projection of the predictor cloud and cannot certify the nonlinear geometric separability that independent-effect claims require. In this dataset, Pearson r = −0.67 and VIF = 1.83 satisfy standard thresholds, yet the nonlinear GAM component exceeds the linear fit by 55% and the suppressor signature remains fully expressed. The operational implication is straightforward. Minimising linear amount–configuration correlation is necessary, but it does not demonstrate separability; metric–amount dependence must be evaluated empirically at the sampling extent used (Fahrig et al. 2019; Wang et al. 2014; Smith et al. 2009). Habitat loss co-generates amount and configuration before sampling, and the realised sample occupies only a restricted portion of the attainable predictor space (Fig. 1a,b). The additive model is therefore asked to estimate an independent configuration contrast from a landscape geometry that rarely supplies it.

The misalignment extends ecologically as well. Subdivision metrics describe how habitat is split but do not encode the spatial arrangement of patches or the resistance of the surrounding matrix. A configuration metric must therefore represent the operative ecological mechanism, not just correlate with abundance, if it is to function as an independent predictor in additive models — a requirement that becomes especially acute in ecosystems without strong patch-matrix structural contrasts, where structural subdivision metrics may fail to capture functional connectivity altogether (Benitez et al. 2024).

Metric choice is not a neutral measurement decision. Landscape-metric research has shown that metric proliferation, redundancy, and composition–configuration confounding make the appropriate metric question-dependent, with objectives, scale, and process guiding metric selection rather than the reverse (Cushman et al. 2008; Gustafson 2019). In the present dataset, a log-transformed patch count produces a stronger linear coupling with habitat amount than the raw count (Amount → log(patches) standardised coefficient = −0.784 vs −0.688 for raw patches), with corresponding changes to suppressor severity. Sensitivity analysis across metric transformations is warranted before interpreting additive fragmentation coefficients (Amiot et al. 2021; Supplementary Material S2 (Section S2.3)). Inference should be assessed across multiple defensible parameterisations, with disagreement among formulations treated as evidence about identifiability rather than ecological resolution. Convergence across parameterisations is more informative than any single coefficient (Koper et al. 2007). Studies should vary predictor and response scale wherever possible, especially in multi-taxa datasets, because fragmentation responses are biologically scale-dependent and a null at one fixed grain cannot resolve process whose relevant scale varies across taxa and diversity components (Jackson and Fahrig 2012, 2015; Martin 2018; McGarigal et al. 2016; Miguet et al. 2016). More broadly, any landscape-scale additive habitat-amount analysis whose conservation conclusions rest on a near-zero configuration coefficient should include empirical demonstration of predictor separability in the realised landscape geometry as a reporting standard (Hadley and Betts 2016; Popovic et al. 2024), rather than treating the coefficient itself as evidence of ecological independence.

### Implications for fragmentation research

The fragmentation debate has largely been conducted within a binary framing that assumes habitat amount and fragmentation can be cleanly separated as parallel additive causes, either habitat amount is the primary driver and fragmentation becomes irrelevant once amount is controlled, or fragmentation has an independent direct effect that survives amount control (Fahrig et al. 2019; Fletcher et al. 2018; Valente et al. 2023). This assumption fails in the present dataset. Habitat amount strongly structures configuration, so the predictors are not parallel causes but hierarchically linked components of the same habitat-loss process, and part of the amount–diversity association operates through fragmentation rather than independently of it (Didham et al. 2012; Ruffell et al. 2016). That question is not answerable from data where the predictors share a dominant landscape gradient. Resolving it requires understanding whether the ecological pathway linking habitat loss to biodiversity is hierarchical or parallel-additive, and whether the model architecture chosen to study it is capable of recovering that distinction (Ruffell et al. 2016). These results are consistent with a hierarchical rather than parallel-additive generating process in which habitat loss drives configuration as a downstream consequence (Püttker et al. 2020), though establishing that hierarchy causally from observational data would require a more nuanced and deliberate integration of study design, metric choice, and model architecture than the field has yet broadly adopted (Ruffell et al. 2016).

Patch-scale evidence indicates that biodiversity in smaller fragments can decline beyond passive-sampling expectations, showing that habitat loss can have real consequences within remnant habitat (Chase et al. 2020a). But those analyses are conducted within individual habitat patches and do not resolve whether spatial configuration has an independent effect at the landscape scale when habitat amount is controlled; patch-scale process evidence and landscape-scale additive identification address different inferential targets. Additive habitat-amount control became the standard landscape-scale operationalisation of the separability requirement, statistically controlling for amount so that configuration metrics could represent fragmentation per se in observational landscapes (Fahrig 2003, 2017; Fahrig et al. 2019, 2026; Hadley and Betts 2016; Smith et al. 2009). That operationalisation does not guarantee the geometric separability it was meant to provide when amount and configuration remain embedded in the same habitat-loss gradient (Didham et al. 2012; Fletcher et al. 2018, 2023a; Ruffell et al. 2016). The persistent divergence between experimental designs which achieve the isolating condition by construction (Haddad et al. 2017) and landscape-scale additive observational designs which operate in the coupled gradient (Fahrig 2017), is likely a design-geometry consequence rather than only an ecological disagreement about mechanism (Fletcher et al. 2023a; Ruffell et al. 2016).

Suppressor geometry provides a parsimonious statistical explanation for part of the divergence between patch-scale and experimental studies, on the one hand, and landscape-scale additive studies, on the other. Patch-scale and experimental studies often report negative fragmentation effects, although those effects vary widely across taxa, spatial scales, and time horizons (Haddad et al. 2015, 2017; Debinski and Holt 2000; Ewers and Didham 2006). Landscape-scale additive studies more often return null or positive configuration coefficients under amount-control frameworks (Fahrig 2017; Watling et al. 2020). This broader divide has become a locked-in debate driven partly by study design (Valente et al. 2023). The suppressor geometry diagnosed here identifies one concrete inferential pathway by which that happens. When habitat-loss landscapes generate nonlinearly coupled amount and configuration predictors, additive models can return stable near-null fragmentation coefficients even when configuration remains strongly aligned with the fitted biodiversity gradient. That possibility does not explain all disagreement in the field, but it does explain why opposed conclusions can emerge from analytically defensible models applied to the same data (Gonçalves-Souza et al. 2025a; Fahrig et al. 2026) without any necessary difference in the underlying ecology (Valente et al. 2023; Fletcher et al. 2023a).

The study-design-driven divergence was quantified directly in Neotropical vertebrates (Vetter et al. 2011). The taxonomic and geographic group dominating the GS25/F26 dataset. Patch-scale studies showed 79% negative species responses while landscape-scale studies showed no significant study-design difference in effect sign (Vetter et al. 2011), showing that the divergence between study architectures is not merely a sampling artifact of the taxa or regions examined but a structural feature of how different designs handle the amount–configuration coupling. Studies that explicitly attempted to decouple the two predictors through strategic landscape selection with PCA-based orthogonalisation (Mortelliti et al. 2010), through natural patch geometry at patch scales (Yaacobi et al. 2007), or through structured subsampling to enforce low amount–fragmentation correlation by design (Melo et al. 2017) also found habitat amount dominant and fragmentation per se negligible. Those studies align with the additive null, but they reached it through analytical architectures that do not share the same predictor-coupling structure.

Fragmentation research shifted from patch-scale to landscape-scale amount-controlled designs once the scale problem became explicit, addressing one inferential layer without resolving the second (Fahrig 2003, 2013, 2017, 2019; Fahrig et al. 2026; Fletcher et al. 2023a). The predictor-separability problem was carried forward rather than solved by that shift. The GS25/F26 dataset provides a direct empirical demonstration of what that carried-forward problem looks like in practice. The same attribution problem had previously been identified at the patch scale. In studies testing for both habitat area and edge effects simultaneously, many were confounded in design or analysis, and nonconfounded analyses more often supported edge than area, implying that some apparent area effects were edge effects in disguise rather than independent patch-size effects (Fletcher et al. 2007). The shift from patch to landscape scale replicated the structure of that problem rather than escaping it.

Landscape-scale studies that used subdivision-based configuration metrics in habitat-loss landscapes and verified predictor independence only with linear diagnostics (Pearson r, VIF; Koper et al. 2007; Smith et al. 2009) are the most vulnerable to suppressor attenuation. In that class, near-null fragmentation coefficients are an expected inferential risk of the analytical architecture, arising from the algebraic identity between patch number, mean patch size, and habitat amount (Fletcher et al. 2023a; Wang et al. 2014) and from the shared gradient-generating process itself, before any appeal to ecological mechanism. Studies that reduced coupling by landscape selection or orthogonalising transformations stand on firmer but still incomplete ground unless nonlinear separability was explicitly checked, residual nonlinear structure can still sustain suppressor attenuation at lower severity, and the GS25/F26 dataset shows directly that linear independence at standard thresholds does not certify the nonlinear separability the interpretation requires (Wang et al. 2014).

The strongest interpretive weight belongs to studies in which configuration varied at approximately constant habitat amount by design, through direct experimental manipulation (Loke et al. 2019; Fletcher et al. 2023b) or strategic landscape selection via PCA-based orthogonalisation (Mortelliti et al. 2010; Trzcinski et al. 1999), structured subsampling (Melo et al. 2017), or natural patch geometry (Yaacobi et al. 2007; Galán-Acedo et al. 2024), or where an equivalent bidirectional nonlinear separability check was demonstrated. Even there, convergence on near-null fragmentation coefficients across multiple studies does not constitute independent ecological convergence by itself. For studies using subdivision-based configuration metrics in habitat-loss landscapes verified only with linear diagnostics, the class this analysis targets most directly, and the class F26’s specification falls within. Convergence on near-null fragmentation coefficients across multiple studies is the expected product of shared analytical structure under the observed predictor geometry, not independent ecological replication. Studies with low but non-zero linear correlations between amount and configuration occupy an intermediate position: Martin et al. (2025b), for example, report a patch density–forest amount correlation of −0.11 across a continental US dataset and justify the additive model on that basis, but low linear correlation does not establish nonlinear separability when the dataset spans a wide cover gradient where the hump-shaped amount–patch number relationship can generate substantial shared nonlinear variance that standard linear collinearity diagnostics fail to detect (Dormann et al. 2013; Koper et al. 2007; Smith et al. 2009; Fahrig 2013; Andrén 1994; Fletcher et al. 2023a). Across all study classes, the prior condition is the same: demonstrated nonlinear separability in the realised predictor space is required before near-null additive configuration coefficients can be read as ecologically decisive (Koper et al. 2007; Smith et al. 2009; Freckleton 2002; Fletcher et al. 2018; Fig. 1d).

The critique developed here applies most directly to landscape-scale observational fragmentation studies that control for habitat amount additively and treat predictor separability as secured by model specification alone, a condition that characterises the majority of such studies in current practice (Hadley and Betts 2016; Fahrig 2017; Riva et al. 2024b). Suppressor attenuation under process-coupled predictors is not specific to fragmentation settings. The inferential problem illustrated here arises wherever such predictors are entered additively into regression models that assume predictor separability.

The benchmark validating partial regression coefficients as unbiased estimators addressed direct effects only, not effects mediated by other model predictors (Smith et al. 2009), and under causal coupling those direct estimates are biased for total ecological effects that include indirect pathways through which habitat loss generates configuration change (Ruffell et al. 2016). Landscape-scale framing of the estimand, statistical control of habitat amount, and replacement of categorical contrasts by continuous predictors are still not enough unless the realised predictor geometry is shown to satisfy the separability condition that gives the additive coefficient ecological meaning (Didham et al. 2012; Fletcher et al. 2018; Fahrig et al. 2019; Hadley and Betts 2016). The alternative approaches Smith et al. (2009) evaluated alongside partial coefficients, including model-averaged coefficients and AIC-based variable importance, do not resolve this limitation. Both Akaike weights and partial correlations mis-rank predictors under correlated conditions in ecological simulation comparisons and overlap allocation between predictors remains method-dependent (Murray and Conner 2009).

Conservation implications that rest on near-zero landscape-scale fragmentation coefficients inherit the same inferential limitation as the coefficients on which they rest. Null effects generated under suppressor geometry cannot establish that configuration is ecologically inert. The recommendation that biodiversity protection should proceed irrespective of patch sizes, where it rests specifically on the F26 reanalysis of this dataset, inherits that inferential limitation. That inferential indeterminacy should not, however, be read as grounds against protecting all habitat irrespective of patch size, for the conservation case for small patches and area-based design for biodiversity conservation rests on complementary and independent evidence that does not depend on additive landscape-scale fragmentation inference (Riva et al. 2022, 2023b, 2024a).

### Summary and prospects for unlocking the fragmentation debate

The inferential limits documented here follow from a structural feature of habitat-loss landscapes, that amount and configuration are co-generated as interdependent components of the same landscape-change process rather than varying as parallel independent causes, and any advance in fragmentation inference must engage that feature directly (Didham et al. 2012; Fahrig 2003; Fletcher et al. 2023a). The three conditions required for independent configuration inference are grounded in the process-to-inference chain Fig. 1 documents across all four panels — habitat loss generates a coupled predictor space (Fig. 1a) that structurally constrains the attainable landscape contrasts (Fig. 1b), producing the SC–BW–R²sp mismatch that signals non-identification under additive models (Fig. 1c–d).

Geometric separability is the prior condition. Without it, a near-zero additive coefficient cannot support claims about independent ecological effects, and additive evidence can address statistical detectability and ecological interpretation only after that first step has been demonstrated, not assumed (Freckleton 2002; Fletcher et al. 2018; Lundberg et al. 2021). Much of the fragmentation debate has treated the second step as if it already secured the third (Didham et al. 2012; Hadley and Betts 2016; Valente et al. 2023).

Operationalising the shift from theory demonstration to theory investigation that Valente et al. (2023) identify as the field’s most productive next step requires testing a concrete methodological hypothesis for why comparable study designs return conflicting results without requiring any difference in the underlying ecology. This reanalysis should also incorporate context-specific stratification by climatic regime, biome-specific land-use history, matrix quality, and taxon-level sensitivity traits. Such stratification is not merely a design refinement. It enables the shift from pooled additive analyses that average across ecological heterogeneity to testing under which conditions fragmentation effects are expected to be detectable (Ewers and Didham 2006).

The diagnostic approach documented in this study provides a replicable template for assessing predictor geometry in any of these databases before fragmentation coefficients are interpreted (Fig. 1). Such reanalysis should report three things explicitly: bidirectional nonlinear coupling asymmetry before and after residualisation, the SC–BW–R²sp mismatch diagnostic, and the evidential status of any subset analysis, allowing readers to evaluate predictor separability rather than infer it from model form. The same coupling diagnostics can be applied prospectively — candidate landscape sets can be screened for nonlinear amount–configuration coupling before data collection begins, minimizing predictor interdependence by design (Hadley and Betts 2016; Fahrig et al. 2019). Landscapes where amount and configuration vary more independently are globally available and could be targeted across biomes (Riva et al. 2024b), though candidate pairs require bidirectional coupling verification. Linear diagnostics alone do not certify that the shared gradient has been sufficiently reduced, since residual nonlinear structure can sustain suppressor attenuation even when linear correlation is low (Koper et al. 2007; Wang et al. 2014). Even where geometric separability is achieved, near-zero direct fragmentation coefficients may still reflect indirect ecological pathways through which habitat loss operates via configuration change rather than ecological absence of fragmentation effects — recovering total effects under causal coupling requires path-analytic frameworks that model those indirect pathways explicitly (Ruffell et al. 2016).

Configuration metrics that attempt amount-independence by construction, i.e., through normalisation by the theoretical maximum or minimum attainable for a given habitat proportion, are more defensible primary inferential vehicles precisely because they reduce the predictor-geometry problem at source (Neel et al. 2004). For already-sampled landscapes, little beyond switching to such metrics can alleviate the problem without redesign. Metrics such as Clumpiness (CLUMPY), the Aggregation Index (AI), and Normalized Landscape Shape Index (nLSI), have shown low nonlinear coupling with habitat amount in both empirical and simulated landscapes (Wang et al. 2014), representing more suitable candidates to index spatial aggregation in patch mosaics generated by habitat-loss-driven fragmentation processes (Neel et al. 2004; Wang et al. 2014). Traditional metrics with stronger amount coupling, including patch count and patch density remain valuable but should be used as robustness checks rather than defaults. Convergence between low-coupling and high-coupling metrics strengthens inference, whereas divergence signal that suppressor attenuation may be driving the result under the coupled specification. The recommendation that “simplest is best” relative to fragmentation metric selection (Martin et al. 2025a) captures a genuine concern for interpretability, but simplicity is not itself an inferential safeguard. A fragmentation metric that achieves simplicity by binding itself to habitat amount through the subdivision identity (Fletcher et al. 2023a) has not simplified the inferential problem, but it has instead embedded it.

## Conclusions

Reliable inference requires models adequate to the data-generating process so that parameter estimates reflect true ecological signal rather than bias. Inferential failures can arise from how the data-generation and representation process shapes what a model can identify, independently of model form (Chadwick et al. 2024). The need for fit-for-purpose models is well established for the response side of ecological models (Guillera-Arroita et al. 2015; Iknayan et al. 2014; Richter et al. 2021). The predictor-side analogue, although articulated conceptually in the context of fragmentation research (Didham et al. 2012; Ruffell et al. 2016), has not been systematically implemented in the empirical landscape ecology literature. The predictor-geometry problem diagnosed here is a specific instance of that gap. In landscapes shaped by progressive habitat loss, amount and configuration are co-generated through a directed causal pathway (Didham et al. 2012; Ruffell et al. 2016). Partial coefficients from additive models are agnostic to this directional interdependence and the predictor-coupling it produces along realised landscape gradients. The inferential limits encountered in coefficient interpretation are therefore inherited from inadequate representation of the system-level process that generates the predictors — not introduced by the model specification alone.

The methodological and ecological components of that diagnosis were each available in the published literature. Among methods for disentangling collinear predictors, partial coefficients perform best under linear orthogonality but fail under nonlinear coupling that linear diagnostics do not detect (Smith et al. 2009; Graham 2003). Under symmetric shared gradients, residuals cannot distinguish between ecological effects of amount and fragmentation. They reallocate shared variance without resolving causal attribution (Koper et al. 2007). Under causal coupling, partial regression gives unbiased direct effects but biased total effects. Path-analytic frameworks are required to recover them (Ruffell et al. 2016). Structure coefficients separate gradient alignment from unique predictor contribution, providing the diagnostic decomposition that partial slopes alone cannot supply (Ray-Mukherjee et al. 2014). A coefficient can be statistically valid as an estimator while its estimand does not correspond to the independent ecological effect the analysis is intended to measure — identifiability requires that the realised predictor geometry supports the contrast the estimand demands (Freckleton 2002; Lundberg et al. 2021). Reliable estimation of fragmentation effects further requires study designs operating at the spatial and ecological scales relevant to the process under investigation (Fahrig 2003; Fahrig et al. 2019; Jackson and Fahrig 2012, 2015; Fletcher et al. 2023a). Fragmentation effects are context-dependent, a structural and ecological source of heterogeneity that dilutes the fragmentation signature in pooled analyses agnostic to the non-random variability that structures the system (Ewers and Didham 2006; Villard and Metzger 2014).

The fragmentation debate has persisted not because of underdeveloped ecological theory, lack of statistical sophistication, nor availability of data against which to test alternative hypotheses (Valente et al. 2023), but in part because advancing fragmentation inference requires three conditions to be satisfied jointly: a statistical architecture adequate to the causal structure of the data-generating process, an estimand that corresponds to an ecological contrast the realised predictor geometry can identify, and explicit stratification of the conditions under which fragmentation effects are expected to be ecologically detectable. These have not been consistently treated as prior conditions before conclusions are drawn. Satisfying each independently is not sufficient. The inferential problem arises precisely when any one of the three is treated as resolved while the others are assumed rather than demonstrated.

What had not been done was the theoretical integration of these components into a coherent inferential architecture and its empirical application to a large-scale fragmentation dataset — showing in practice what process-driven predictor coupling and unmodelled ecological heterogeneity produce for the coefficients on which the fragmentation debate has turned. The present analysis provides that integration and empirical grounding for the first time.

In revisiting the GS25–F26 dataset, this study shows that the full-dataset null is non-diagnostic: whenever the shared habitat-amount–configuration constraint is reduced, the recoverable fragmentation-associated signal is consistently and non-trivially negative rather than null. That pattern does not identify an independent fragmentation effect size; instead, it defines an inferential limit. In landscapes shaped by habitat loss, additive control for habitat amount does not by itself generate the contrast required for fragmentation-per-se inference. Habitat amount control, as F26 correctly insist, is necessary, but it becomes sufficient only when the realised predictor space is shown to support geometric separability (Martínez-Lanfranco 2026) while ecological heterogeneity is also considered. More broadly, this helps explain why the fragmentation debate has produced opposed conclusions from comparable data for decades. When habitat loss co-generates habitat amount and configuration, a near-zero additive fragmentation coefficient can be an expected consequence of the analytical architecture, arising from both the algebraic dependence of subdivision metrics on habitat amount and the landscape-generating process that produces them, rather than evidence that configuration is ecologically irrelevant. Progress will depend less on stronger positions or additional additive nulls than on demonstrating that the landscapes analysed actually identify the ecological contrast under dispute.

## Supporting information

Supplementary Material S1 Design and Geometry

Supplementary Material S2 Results and Sensitivity

Supplementary Material S3 Derivations and Code

## Acknowledgements

This work was supported in part by the National Agency for Research and Innovation (*Agencia Nacional de Investigación e Innovación*, ANII, Uruguay), the BIOS^2^ Computational Biodiversity Science and Services Program, and the Bayne Lab (Department of Biological Sciences, University of Alberta). I am grateful to Isabelle Lebeuf-Taylor and other colleagues in the Bayne Lab for valuable discussions and feedback during the conceptualization of this piece. I also thank the authors of the original studies for making their data and code available, which made this synthesis possible.

## Data and Code Availability

The dataset was originally compiled and analyzed by Gonçalves-Souza et al. (2025a,b) and is freely available at Zenodo (https://zenodo.org/records/12206838). The same dataset was reanalysed by Fahrig et al. (2026), with the code available from https://osf.io/hu3xw. Code for all diagnostic workflows and analysis of this study is available at https://github.com/jacoloml/Reanalyses_GS25_F26 (archived at https://doi.org/10.5281/zenodo.19546467).

