## Supplementary Material S1 Design and Geometry for "Habitat-loss-driven predictor coupling limits inference about the independent effects of configuration in additive habitat-amount models: implications for the fragmentation debate"

*Juan Andrés Martínez-Lanfranco*

### Table of Contents

**S1.1 Predictor distributions**
- S1.1.1 Amount-dominance in shared predictor space
- S1.1.2 Design-reconstruction diagnostic

**S1.2 Parameterisations, orthogonality diagnostics, and formal variance decomposition**
- S1.2.1 Parameterisations
- S1.2.2 Raw predictor coupling
- S1.2.3 Residual coupling after parameterisation
- S1.2.4 Residualisation asymmetry

**S1.3 References**

**Tables**

Table S1.1 Distributional statistics for the two landscape-level predictors
Table S1.2 Model parameterisations (M1–M4)
Table S1.3 Raw predictor coupling in both diagnostic orientations (n = 74 landscapes)
Table S1.4 Residual coupling diagnostics after parameterization

**Figures**

Figure S1.1 Joint predictor space (habitat amount × fragmentation) and marginal distributions by landscape type

### S1 Empirical conditions motivating diagnostic analyses

Design-level diagnostics in this section document the realised predictor structure of the GS25–F26 (Gonçalves-Souza et al. 2025 and Fahrig et al. 2026, respectively) dataset and provide the empirical context for the suppressor analysis in the main text. The same design evidence is developed as the primary argument of a companion commentary (Martínez-Lanfranco 2026, EcoEvoRxiv preprint); results reported here are complementary and draw on the same underlying data. The logistic partR² analysis in the main text and Section S1.1.1 is an independent statistical application of the suppressor diagnostic framework to the design contrast.

#### S1.1 Predictor distributions

Three features of the realised predictor space motivate the diagnostic analyses applied in the main text: the distributional properties of the two predictors, their structural size asymmetry, and the scarcity of same-amount, different-configuration contrasts. All three are documented here from the F26 OSF dataset. The dataset comprises 74 landscapes from 37 independent multi-taxa studies (Gonçalves-Souza et al. 2025, as compiled by Fahrig et al. 2026). Each study contributes exactly two landscapes classified by GS25 as continuous or fragmented — a categorical factor that encodes, in coarsened form, the same gradient captured by the two continuous predictors used by F26: habitat amount (% forest cover within the 2-km buffer) and number of patches (the F26 operational fragmentation metric; used throughout as shorthand for "fragmentation" or "fragmentation predictor" where context is clear). The terms "fragmentation per se" and "configuration" are reserved for the ecological concept of landscape structure independent of habitat amount. The habitat-amount distribution is right-skewed and concentrated at high cover values (median 70%, IQR 44%): 34 of 74 landscapes from 17 studies (45.9%) have both landscapes exceeding 50% cover (study-level minimum filter), while the low-cover tail — where the nonlinear amount–patches constraint is steepest — is represented by a minority of landscapes. Patch number is also right-skewed with high variance (median 30, IQR 62), reflecting structural heterogeneity concentrated at intermediate cover values. The two landscape types are also strongly asymmetric in size: reconstructing the per-study ratio of largest continuous forest to smallest fragment from GS25's Supplementary Table 7 (Gonçalves-Souza et al. 2025) yields a median of 1,672× (range: 4.6× to 10,169×). This structural size asymmetry means the two landscape types occupy different regions of the habitat-amount distribution by construction, explaining why same-amount, different-configuration contrasts are structurally scarce in the realised predictor space. Note on source attribution: all landscape-level predictor values used in this supplement are computed from the F26 OSF dataset. The 1,672× median size ratio is a reconstruction from GS25 Supplementary Table 7 (Martínez-Lanfranco 2026, EcoEvoRxiv preprint). Passages citing GS25's Extended Data Figures or Supplementary Texts refer to published GS25 source materials (Gonçalves-Souza et al. 2025). Table S1.1 reports full distributional statistics for both predictors; Figure S1.1 shows their marginal distributions and joint predictor space by GS25 landscape type.

***Table S1.1.*** *Distributional statistics for the two landscape-level predictors. 34 landscapes from 17 studies meet the corrected study-level minimum filter (both landscapes ≥50%); the filter operates at study level (both landscape types must exceed 50% cover; fragmented paired landscapes within selected studies may be below 50%); the distribution is right-skewed and concentrated at higher values (mean 66.9%, median 69.9%). Patch number is right-skewed (median 30, mean 48.2) with high variance (SD 56.1), reflecting structural variation at intermediate cover values. Amount values are percentages of the 2-km buffer area.*

| **Quantity** | **Habitat amount (%)** | **Number of patches** |
|---|---|---|
| n landscapes | 74 | 74 |
| n studies | 37 | 37 |
| Landscapes per study (min/median/max) | 2 / 2 / 2 | 2 / 2 / 2 |
| Mean | 66.9 | 48.2 |
| Median | 69.9 | 30 |
| SD | 26.0 | 56.1 |
| IQR | 43.9 | — |
| % below 20% | 5.4% (4/74) | — |
| % below 30% | 12.2% (9/74) | — |
| % above 50% | 71.6% (53/74) | — |
| **Study-mean range (min/median/max)** | **34.5 / 65.8 / 98.4** | **—** |

#### S1.1.1 Amount-dominance in shared predictor space

A mixed-effects logistic regression of the GS25 categorical landscape classification (continuous vs. fragmented) on the two continuous F26 predictors, with study as a random intercept, was applied as a diagnostic test of whether the same suppressor geometry that operates in the diversity regression also structures the representational overlap between the two study designs. The test asks whether the SC–BW–$R^2$sp mismatch diagnostic, developed in Section S1.2 for the diversity response, reappears when the response is landscape type rather than species richness. The marginal $R^2$ (fixed effects only) was 0.452; the conditional $R^2$ was identical, confirming that the study random intercept explained no additional variance beyond the fixed predictors — the between-study variation in landscape type is fully captured by the two continuous predictors at the 2-km buffer scale. The Tjur $R^2$ on marginal fitted values was 0.352, indicating that 35.2% of the categorical classification is captured by the predictor pair at the population level, with 64.8% remaining unexplained by the buffer-scale metrics. Structure coefficients and semi-partial $R^2$ were computed using the same SC/BW/$R^2$sp diagnostic framework applied to the diversity models (Section S1.2). The pattern mirrors the cross-over suppressor signature in Table 1: both predictors had large structure coefficients (SC amount = −0.985; SC patches = +0.708), indicating that both align strongly with the fitted classification gradient. However, the partial coefficients diverged sharply: the beta weight for amount was large (BW = −1.613) while the beta weight for patches was near zero (BW = +0.048), and the semi-partial $R^2$ for patches was effectively zero ($R^2$sp = −0.10%) against $R^2$sp = +2.60% for amount. The near-zero $R^2$sp for patches ($R^2$sp = −0.10%; slightly negative values are bootstrap artifacts at the zero boundary, not literal negative explained variance) is consistent with the suppressor signature: patches track the classification gradient (large SC) but contributes no independent explanatory variance once amount is controlled. This is the same identification constraint that operates in the diversity regression — the two predictors share a directed habitat-loss gradient that prevents their independent contributions from being isolated by additive control, whether the response is species diversity or landscape type itself. The result does not favour one study design over the other; it shows that the representational overlap between the two frameworks is structured by the same amount-dominated gradient that drives the suppressor attenuation in the main analysis. This directional structure is not an artefact of a single analytical choice: GS25's own Extended Data Figs. 3 and 4 (Gonçalves-Souza et al. 2025) show that continuous landscapes have consistently higher habitat amount and fragmented landscapes consistently more patches across the full 200–2,000 m buffer series — the same directional contrast that the logistic decomposition above quantifies at the 2-km grain.

**S1.1.2 Design-reconstruction diagnostic**

To evaluate whether the categorical landscape contrast corresponds to regions of the continuous predictor space used in the additive model, we compared fragmented and continuous landscapes directly within each of the 37 paired studies on standardised number of patches and habitat amount. This diagnostic requires no model fitting and is not subject to the collinearity or generated-regressor problems associated with logistic-residual approaches. For each study, two pairwise inequalities were evaluated: (i) whether the fragmented landscape contained more patches than the continuous landscape, and (ii) whether the fragmented landscape had lower habitat amount. Both conditions are expected if the categorical design encodes the same landscape gradient captured by the continuous predictors.

Fragmented landscapes exceeded continuous landscapes on patch count in 31 of 37 studies (84%) and had lower habitat amount in 35 of 37 studies (95%). Both conditions held simultaneously in 31 of 37 studies (84%). GS25 themselves acknowledged in Supplementary Text 4 (Gonçalves-Souza et al. 2025) that pairwise comparisons with precisely the same habitat amount could not be selected within studies because too few studies contained a sufficient number of fragments in both landscape types at the same habitat-amount level — a constraint the present design-reconstruction diagnostic confirms empirically: only 3 of 37 study pairs had $|\Delta HA| \leq 5\%$, and only 2 of 37 satisfied that constraint within the highest habitat-amount class (77–100%). The two studies where habitat amount violated the expected direction (Lima_1999_a and Lima_1999_b) were a strict subset of the six studies where patch count did not discriminate landscape type. The result was consistent across geographic regions: 12 of 13 non-South American studies (92%) and 19 of 24 South American studies (79%) satisfied both conditions simultaneously, indicating that the pattern is not driven by a single continent.

The six studies where patch count did not encode the expected design contrast fall into three distinct categories. Two studies had identical patch counts in both landscape types (Nemesio_2010: 52 patches each; Rocha_2016: 1 patch each), representing uninformative ties rather than genuine design violations. Three studies had habitat cover exceeding 88% in at least one landscape (Chiarello_1999: 91%/94%; Lima_1999_a: 99%/94%; Lima_1999_b: 98%/89%), where the design contrast reflects configurational differences rather than the amount-driven gradient typically present at lower cover levels. One study (Bossart_2016, Africa) showed a genuine reversal (31 vs. 72 patches; 66% vs. 75% habitat amount) with no structural explanation from the available predictors.

These results demonstrate that the GS25 categorical classification largely corresponds to regions of the continuous predictor space used in the F26 additive model. This is the dataset condition under which suppressor attenuation can arise: when the categorical design and the continuous predictors encode the same underlying landscape gradient, additive regression assigns shared variance to the dominant predictor and attenuates the partial coefficient of the other, producing near-zero coefficients that do not reflect ecological absence. The equal-amount, different-configuration contrast that fragmentation-per-se inference requires is therefore not directly recoverable from this dataset by reparameterisation alone. A conclusion independently supported by GS25's own acknowledgment in Supplementary Text 4 (Gonçalves-Souza et al. 2025) that such contrasts are infeasible given the empirical distribution of habitat amounts across landscape pairs.

### S1.2 Parameterisations, orthogonality diagnostics, and formal variance decomposition

#### S1.2.1 Parameterisations

The orthogonality criterion described in this section is used as a dataset-specific diagnostic for the GS25–F26 predictor geometry; it is not presented as a general requirement for additive ecological models. Four parameterisations span the residualisation space between habitat amount and fragmentation (Table S1.2). M1 is the additive F26 baseline; M2 and M3 are diagnostic decompositions of the shared habitat-loss gradient; M4 is a negative control for bidirectional residualisation.

**M1 (additive baseline).** Diversity is modelled as a function of raw fragmentation and raw habitat amount, reproducing the Fahrig et al. (2026) specification. Coefficients estimate partial associations after linear adjustment for the other predictor, under the assumption that predictors span separable dimensions of predictor space.

**M2 (fragmentation residualised on amount).** Fragmentation is residualised on habitat amount; diversity is modelled as a function of residual fragmentation plus raw habitat amount. The fragmentation coefficient estimates the association of diversity with the component of configuration that is statistically independent of habitat amount. This is not fragmentation per se in the ecological sense — it is fragmentation variation with its association to amount removed.

**M3 (amount residualised on fragmentation).** Habitat amount is residualised on fragmentation; diversity is modelled as a function of raw fragmentation plus residual amount. Symmetrically, the amount coefficient estimates the component of habitat amount independent of configuration. In the present dataset, M3 is diagnostically informative because it is the only parameterisation achieving verified near-zero residual nonlinear coupling.

**M4 (bidirectional residualisation; negative control).** Both predictors are residualised and the pair is evaluated as a negative control. In this dataset, M4 fails because it reintroduces substantial nonlinear coupling (residual GAM $R^2 = 0.63$) rather than eliminating it, showing that the orthogonality achieved by M3 is not a trivial consequence of residualising predictors in any direction.

***Table S1.2.*** *Model parameterisations. M3 (shaded) is the only parameterisation achieving diagnostically verified residual uncoupling. M4 confirms that bidirectional residualisation does not resolve the shared gradient.*

| | Frag. predictor | Amount predictor | Estimand | Orthogonality |
|---|---|---|---|---|
| M1 | Patches (raw) | Amount (raw) | Partial association after linear adjustment | Not verified — raw predictors share nonlinear gradient |
| M2 | resFrag (patches − $f$(amount)) | Amount (raw) | Configuration variation independent of amount (linear sense only) | Partial — linear $r \approx 0$ but nonlinear coupling retained |
| M3 | Patches (raw) | resAmount (amount − $f$(patches)) | Amount variation independent of configuration | Verified ✓ — residual GAM $R^2 \approx 0$ in both orientations |
| M4 | resFrag | resAmount | Negative control | Fails ✗ — reintroduces nonlinear coupling (GAM $R^2$ = 0.63) |

#### S1.2.2 Raw predictor coupling

Linear and generalised additive model (GAM) fits were computed in both diagnostic orientations (Table S1.3). Raw coupling was strong and markedly asymmetric: the nonlinear component was 7.75-fold larger in the reverse direction (Amount ∼ s(Patches)) than in the forward (Patches ∼ s(Amount)), while linear $R^2$ was identical in both directions (0.454) by definition. Pearson $r$ = −0.67; VIF = 1.83. These standard linear diagnostics detect only the linear component of predictor coupling; the nonlinear component documented by the asymmetric GAM fits above remains after linear partialling. Crucially, VIF characterises variance inflation in regression coefficient standard errors and was developed for prediction accuracy assessment, not for detecting distortion of coefficient direction through shared gradient allocation. Neither metric is sensitive to the defining feature of a suppressor: a predictor that is strongly aligned with the model-predicted response gradient (large SC) yet contributes a near-zero partial regression coefficient (BW ≈ 0). In this dataset, a predictor pair satisfying both conventional thresholds is associated with a ~51-fold SC-to-BW discrepancy (α median SC = −0.725; α median BW = −0.014) — the defining signature of cross-over suppression — demonstrating that datasets within conventional collinearity thresholds may still harbour suppressor structures whenever predictors share a dominant landscape gradient. GS25's own Extended Data Fig. 5 (Gonçalves-Souza et al. 2025) provides descriptive published evidence consistent with this pattern: number of patches, patch density, edge density, and mean perimeter–area ratio all vary nonlinearly across habitat-amount classes, with the continuous–fragmented gap narrowing markedly in the highest (90–100%) habitat-amount class (Gonçalves-Souza et al. 2025).

***Table S1.3.*** *Raw predictor coupling in both diagnostic orientations (n = 74 landscapes). Nonlinear component = GAM R² − linear R². Linear R² is identical in both orientations by definition; the nonlinear component is asymmetric, being 7.75-fold larger in the reverse direction (ratio computed from unrounded pipeline values; tabulated values are rounded to two decimal places).*

| Diagnostic orientation | Linear $R^2$ | GAM $R^2$ | Nonlinear component | Pearson r |
|---|---|---|---|---|
| Forward: Patches ~ s(Amount) | 0.45 | 0.49 | 0.03 | -0.67 |
| Reverse: Amount ~ s(Patches) | 0.45 | 0.70 | 0.25 | -0.67 |

#### S1.2.3 Residual coupling after parameterisation

Residual coupling was evaluated post-residualisation using the same bidirectional GAM diagnostics (Table S1.4). M2 zeroed the linear correlation but retained substantial nonlinear coupling in both orientations and therefore failed the orthogonality criterion. M3 achieved near-zero residual GAM $R^2$ in both orientations — the only parameterisation satisfying the criterion in this dataset. M4 reintroduced strong coupling (GAM $R^2$ = 0.63), confirming that orthogonality is not achieved by bidirectional residualisation per se.

The orthogonality criterion used here is stricter than linear predictor separability, as applied to the GS25–F26 dataset. Linear separability requires only low shared variance (Pearson r ≈ 0); the criterion here additionally requires that the nonlinear gradient structure is removed, verified by near-zero residual GAM $R^2$ in both diagnostic orientations. M2 satisfies the linear condition but fails the nonlinear criterion.

***Table S1.4.*** *Residual coupling diagnostics. M2 zeroes the linear correlation but retains nonlinear coupling in both orientations. M3 achieves near-zero residual GAM R² in both orientations (orthogonality criterion satisfied). M4 reintroduces strong nonlinear coupling.*

| Parameterisation | Residual GAM $R^2$ (fwd) | Residual GAM $R^2$ (rev) | Pearson r | Orthogonality |
|---|---|---|---|---|
| M1 (raw predictors) | 0.70 | — | -0.67 | None |
| M2 (resFrag + raw Amount) | 0.44 | 0.44 | ≈ 0 | Partial only |
| M3 (Patches + resAmount) | ≈0 | ≈0 | ≈ 0 | Verified ✓ |
| M4 (resFrag + resAmount) | 0.63 | — | ≈ 0 | Fails ✗ |

#### S1.2.4 Residualisation asymmetry

Residualising one predictor on the other changes which predictor retains the shared gradient, not whether the shared gradient exists. Residualising fragmentation on amount (M2) allocates the shared component to the amount axis; residualising amount on fragmentation (M3) reverses the allocation. Because the coupling is nonlinear and asymmetric across diagnostic orientations, no single residualisation removes the shared gradient equally from both predictors simultaneously

— M4 confirms this by reintroducing rather than eliminating coupling. M2 and M3 are therefore diagnostic decompositions of the same shared-gradient geometry, not alternative ecological models. The apparent shift in predictor importance across parameterisations reflects gradient reallocation rather than discovery of independent ecological signal (Koper et al. 2007). The practical consequence is an estimand mismatch: the coefficient each parameterisation returns estimates a different conditional quantity, and only the parameterisation achieving verified residual uncoupling reduces the geometric constraint sufficiently to be diagnostic — it does not recover a definitive estimate of fragmentation per se, but it is the least confounded quantity the dataset supports.

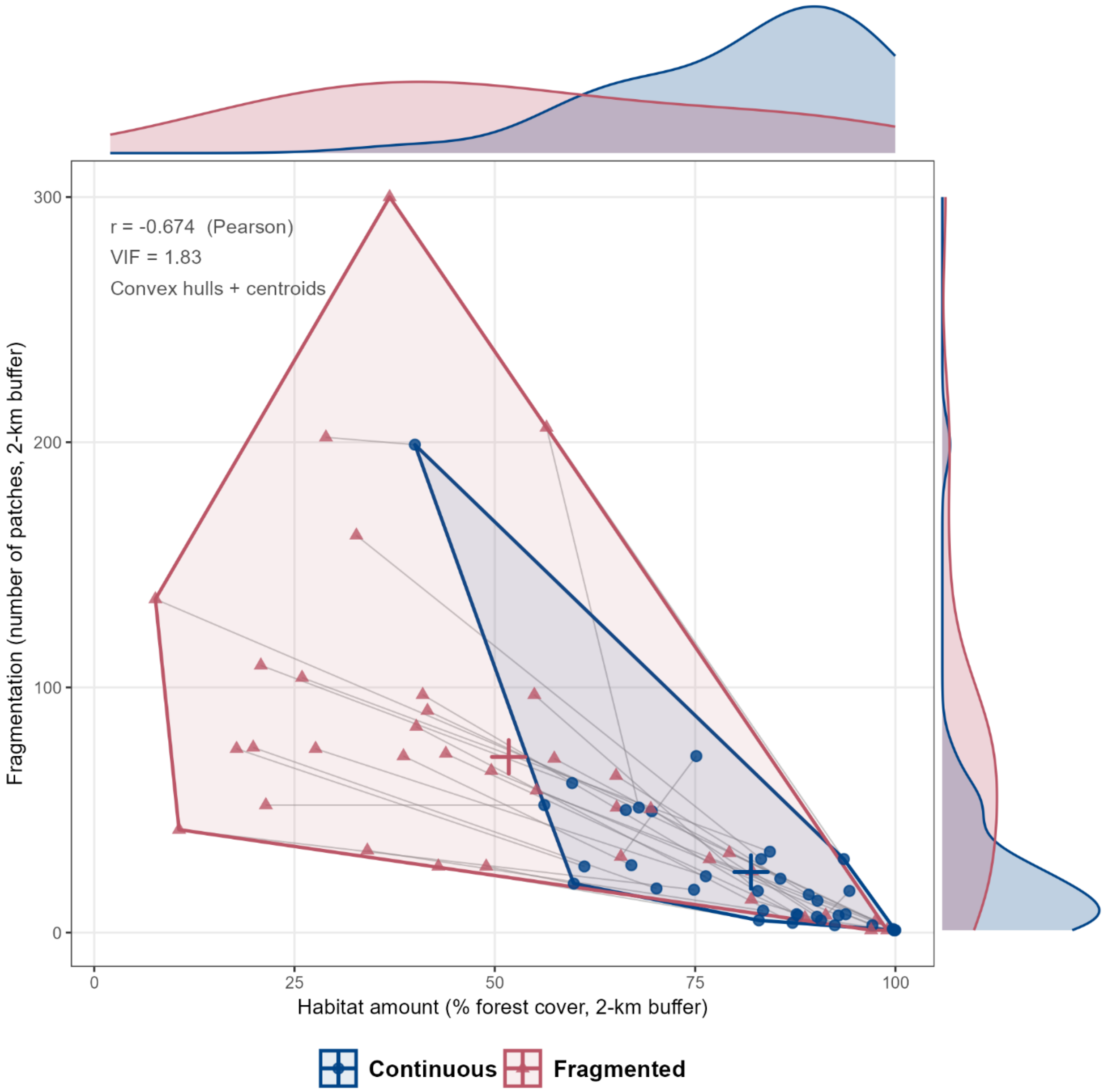


**Figure S1.1.** Joint distribution of the two landscape-level predictors — habitat amount (% forest cover, 2-km buffer; x-axis) and fragmentation (number of patches, 2-km buffer; y-axis) — for all 74 landscapes from 37 studies, separated by GS25 landscape type (continuous: blue circles; fragmented: red triangles). Marginal density plots show the univariate distribution of each predictor by landscape type. Convex hulls delineate the realised predictor space of each group; crosshairs mark group centroids. Pearson $r = -0.674$; $VIF = 1.83$. The negative correlation and overlapping convex hulls illustrate the shared habitat-loss geometry that motivates the suppressor analysis.
