## Supplementary Material S2 Results and Sensitivity for "Habitat-loss-driven predictor coupling limits inference about the independent effects of configuration in additive habitat-amount models: implications for the fragmentation debate"

*Juan Andrés Martínez-Lanfranco*

### Table of Contents

**S2.1 Full-dataset suppressor diagnostics**
- S2.1.1 Full dataset — all 18 variants

**S2.2 Sensitivity analyses and subset analyses**
- S2.2.1 High-cover subset (≥50% habitat amount)
  - S2.2.1.1 Fragmentation predictor
- S2.2.2 Tropical-only subset
- S2.2.3 Continental subsets (SA vs non-SA)
  - S2.2.3.1 Suppressor diagnostics by continental group
  - S2.2.3.2 Habitat-amount overlap, biome contingency, and power
- S2.2.4 Crossed tropical sensitivity checks
- S2.2.5 Ecological conditionality

**S2.3 Path model — causal hierarchy check**
- S2.3.1 Full 18-variant results
- S2.3.2 Continental sensitivity
- S2.3.3 AIC comparison (directed vs correlated specification)
- S2.3.4 $R^2$ and residual diagnostics
- S2.3.5 Log-patches robustness

**S2.4 Model diagnostics**
- S2.4.1 Residual diagnostics
- S2.4.2 Random-effect stability
- S2.4.3 AICc comparisons

**S2.5 References**

**Tables**


Table S2.1 Full diagnostic table corresponding to main-text Table 1
Table S2.2 α diversity — fragmentation predictor (SC, BW, $R^2$sp; M1–M4)
Table S2.3 α diversity — habitat amount predictor
Table S2.4 β diversity — fragmentation predictor
Table S2.5 β diversity — habitat amount predictor
Table S2.6 γ diversity — fragmentation predictor
Table S2.7 γ diversity — habitat amount predictor
Table S2.8 Subset analysis design: sample sizes, rationale, and evidential tiers
Table S2.9 Predictor coupling and design-reconstruction diagnostics — tropical-only subset
Table S2.10 M1 and M3 fragmentation coefficient summary — tropical-only subset
Table S2.11 M1–M4 fragmentation BW and SC by continental group (all 18 variants)
Table S2.12 Path model coefficients across all 18 analytical variants
Table S2.13 Path model coefficients — Tropical SA groups, all analytical variants
Table S2.14 Random-effect stability across parameterisations


**Figures**


Figure S2.1 Bidirectional GAM coupling diagnostics, full dataset (M1–M4)
Figure S2.2 Variance partitioning — fragmentation predictor, all 18 variants (M1–M4)
Figure S2.3 Variance partitioning — habitat amount predictor, all 18 variants (M1–M4)
Figure S2.4 M4 negative control: SC×BW biplot across parameterisations M1–M4
Figure S2.5 Coupling diagnostics, global ≥50% cover subset (n = 17 studies)
Figure S2.6 SC–BW forest plot, global ≥50% cover subset
Figure S2.7 Coupling diagnostics, Tropical SA — full coverage range (n = 23 studies)
Figure S2.8 SC–BW forest plot, Tropical SA — full coverage range
Figure S2.9 Coupling diagnostics, Tropical SA ≥50% cover (n = 9 studies)

**Table S2.1.** Full diagnostic table corresponding to the compact main-text Table 1. Suppressor diagnostics across model parameterisations (M1–M3) for α and γ diversity. SC = structure coefficient (cor[predictor, $\hat{y}$]); BW = standardised beta weight (partial regression coefficient per 2 SD); $R^2$sp = semi-partial $R^2$ (unique variance in observed response attributable to each predictor after controlling for the other). Large |SC| with BW ≈ 0 and $R^2$sp ≈ 0 is the pattern predicted by cross-over suppressor attenuation (Smith et al. 2009). All values are medians across 6 analytical variants (3 diversity-order metrics × 2 pairing designs). M1 replicates the Fahrig et al. (2026) additive specification. M3 is the only parameterisation meeting the dataset-specific residual-uncoupling criterion (residual GAM $R^2 < 0.02$ in both orientations; criterion met) in both Section A and B; M2 fails; M4 (negative control, not shown) reintroduces strong coupling — full M4 results in Figure S2.4 (see also Section S2.1). Section B presents two Tier 1 conditionality subsets: B1 (Global ≥50% cover, n = 17, both landscapes ≥50% cover, study-level minimum filter) evaluates geometric robustness; B2 (Tropical SA, n = 23, 0–100% cover) evaluates biome-level heterogeneity. β diversity excluded from the main table (no consistent directional pattern; Supplementary Material 2 (Section S2.1)); the main inferential burden is carried by α and γ diversity, where the null is deployed to support claims of ecological independence. Full variant-level outputs: Sections S2.1, S2.2.3.1, and S2.2.4 of this Supplementary Material. M4 results (negative control): Figure S2.4. F26 = Fahrig et al. 2026.

| Diagnostic | M1 — Additive (F26) | M2 — resFrag + Amt | M3 — Frag + resAmt ✓ verified | Notes |
|---|---|---|---|---|
| **Section A · Full dataset (n = 74 landscape pairs · 37 studies)** | | | | |
| **Predictor coupling (residual GAM $R^2$, fwd / rev)** | | | | |
| Residual nonlinear coupling | Nonlin rev = 0.243; fwd = 0.032; asymmetry = 7.75× | 0.444 / 0.444 | 0.000 / 0.008 | M1 fwd = raw GAM $R^2$ (no residualisation). M2 fwd = rev by symmetry. M3 satisfies dataset-specific criterion (residual GAM $R^2 <$ 0.02 in both orientations; criterion met). M4 reintroduces coupling ($R^2$ = 0.63). |
| **Fragmentation predictor (patch number) — α / γ diversity [medians across 6 variants]** | | | | |

| | | | | |
|---|---|---|---|---|
| Structure coefficient (SC) | −0.725 / −0.747 | −0.198 / −0.218 | −0.896 / −0.908 | |
| Beta weight (BW) | −0.014 / −0.020 | −0.030 / −0.034 | −0.114 / −0.101 | M3: 6/6 variants negative for both α and γ. Rarefaction attenuates M3 BW ~46% (α) and ~29% (γ) vs observed richness. |
| Semi-partial $R^2$ ($R^2sp$) | 0.0001 / 0.0002 | 0.0007 / 0.0012 | 0.0086 / 0.0074 | 144-fold increase M1→M3 for α; 37-fold for γ. Negligible unique variance in M1 despite large SC. |
| **Habitat amount predictor (forest cover) — α / γ diversity [medians across 6 variants]** | | | | |
| Structure coefficient (SC) | +0.997 / +0.992 | +0.978 / +0.972 | +0.443 / +0.417 | |
| Beta weight (BW) | +0.108 / +0.108 | +0.127 / +0.122 | +0.041 / +0.047 | |
| Semi-partial $R^2$ ($R^2sp$) | 0.0068 / 0.0075 | 0.0182 / 0.0168 | 0.0041 / 0.0045 | |
| **Suppressor diagnosis** | | | | |
| Suppressor type (Smith et al. 2009) | Cross-over: large \|SC\|, BW ≈ 0; 2/12 sign reversals; ~51-fold SC/BW ratio; ~8,800-fold shared/unique (α) | Partial: mismatch reduced; nonlinear coupling persists (GAM $R^2$ = 0.44) | Attenuation no longer expressed under M3: BW consistently negative for both α and γ; residual coupling ≈0 in both orientations | M3 is the only parameterisation satisfying the orthogonality criterion; its coefficients are diagnostic, not definitive estimates of fragmentation per se. |

| Section B1 · Global ≥50% cover (n = 17 studies \| both landscapes ≥50% \| criterion met \| Tier 1) | | | | |
|---|---|---|---|---|
| **Predictor coupling** | | | | |
| Residual nonlinear coupling | Nonlin rev = 0.259; fwd = 0.016; asymmetry = 16.2× | — | ≈0.000 / 0.000 | M3 orthogonality verified (criterion met). Reverse nonlinear component (0.259) exceeds forward (0.016); 16.2× asymmetry vs 7.75× full dataset. |
| **Fragmentation predictor (patch number) — α / γ diversity [medians across 6 variants]** | | | | |
| Structure coefficient (SC) | −0.951 / −0.953 | −0.273 / −0.350 | −0.932 / −0.940 | Suppressor SC–BW mismatch preserved. Large \|SC\| at M1 and M3. |
| Beta weight (BW) | −0.077 / −0.060 | −0.033 / −0.039 | −0.122 / −0.099 | M3 BW amplified vs full dataset (−0.122 vs −0.114 α; −0.099 vs −0.101 γ). 6/6 variants negative at M1 and M3. |
| Semi-partial $R^2$ ($R^2$sp) | 0.004 / 0.003 | 0.003 / 0.003 | 0.015 / 0.010 | Full variant-level bootstrap CIs in Section S2.1. |
| **Habitat amount predictor (forest cover) — α / γ diversity [medians across 6 variants]** | | | | |
| Structure coefficient (SC) | +0.945 / +0.941 | +0.961 / +0.937 | +0.362 / +0.342 | Amount SC remains positive under M3 (contrast with Tropical SA subset). |

| | | | | |
|---|---|---|---|---|
| Beta weight (BW) | +0.057 / +0.046 | +0.118 / +0.103 | +0.033 / +0.029 | β-weights directional only; $R^2$sp needed to interpret magnitude. |
| Semi-partial $R^2$ ($R^2$sp) | 0.005 / 0.004 | 0.015 / 0.012 | 0.004 / 0.004 | |
| **Suppressor diagnosis** | | | | |
| Suppressor type | Cross-over: large \|SC\|, BW 0; 6/6 sign-consistent | Partial: SC–BW mismatch reduced | Attenuation resolved: BW consistently negative 6/6 | Geometric robustness: suppressor signature preserved with clean paired design. M3 BW marginally more negative than full dataset. Amount SC stays positive under M3—amount retains independent positive alignment globally at high cover. Criterion met. |
| **Section B2 · Tropical SA — 0–100% cover (n = 23 studies \| criterion met \| Tier 1 \| biome conditionality)** | | | | |
| **Predictor coupling** | | | | |
| Residual nonlinear coupling | Nonlin rev = 0.323; fwd = 0.000; asymmetry = ∞ (reverse-only) | — | ≈0.000 / 0.000 | M3 orthogonality verified (criterion met). Nonlinear coupling exclusively reverse: habitat loss generates configuration change; forward nonlinear = 0. Strongest asymmetry across all subsets. |
| **Fragmentation predictor (patch number) — α / γ diversity [medians across 6 variants]** | | | | |
| Structure coefficient (SC) | −0.869 / −0.904 | −0.308 / −0.536 | −0.996 / −0.966 | M3 SC approaches −1.000 (α): near- |

| | | | | |
|---|---|---|---|---|
| | | | | total alignment of fragmentation with fitted diversity gradient once amount gradient removed. |
| Beta weight (BW) | −0.049 / −0.106 | −0.036 / −0.077 | −0.113 / −0.143 | M3 BW more negative for γ (−0.143 vs −0.101 full dataset). 6/6 negative at M1 and M3. |
| Semi-partial $R^2$ ($R^2$sp) | 0.0002 / 0.001 | 0.0002 / 0.001 | 0.004 / 0.006 | R $R^2$sp available from full partR2 workflow at n = 23. |
| **Habitat amount predictor (forest cover) — α / γ diversity [medians across 6 variants]** | | | | |
| Structure coefficient (SC) | +0.950 / +0.822 | +0.950 / +0.822 | −0.091 / −0.257 | M3 amount SC flips negative: once fragmentation residualised out, amount loses positive gradient alignment in tropical SA. Consistent with configuration dominating residual diversity variation in this biome. |
| Beta weight (BW) | +0.081 / +0.038 | +0.106 / +0.101 | −0.010 / −0.036 | M3 amount BW near zero or negative: direct amount effect absent once shared gradient removed in tropical SA. |
| Semi-partial $R^2$ ($R^2$sp) | 0.003 / 0.003 | 0.003 / 0.003 | 0.002 / 0.004 | |
| **Suppressor diagnosis** | | | | |

| | | | | |
|---|---|---|---|---|
| Suppressor type | Cross-over: large \|SC\|, BW 0; 6/6 sign-consistent | Partial: SC–BW mismatch reduced | Attenuation resolved: BW consistently negative 6/6 | Biome heterogeneity: suppressor signature intensified in tropical SA. M3 SC approaches −1.000 and amount SC flips negative under M3—configuration dominates once habitat-loss gradient removed. Contrast with the Global ≥50% cover subset (amount SC stays positive under M3) confirms that the amount–configuration hierarchy is biome-specific. |

**Note:** Full variant-level outputs: Supplementary Material S1 (Section S1.2), Supplementary Material S2 (Section S2.1), and Supplementary Material S3 and S2.2.3.1 and S2.2.4. M4 results (negative control): Figure S2.4 (see also Section S2.1). Semi-partial $R^2$ 95% bootstrap CIs reported in Section S2.1. The path model (main text Figures 5 and 6) shows that restricting to ≥50% cover landscapes (study-level minimum filter; both landscapes must exceed 50%, $n = 17$) strengthens the direct Fragmentation → Diversity path (Global ≥50% cover ($n = 17$, both landscapes ≥50%)): $\alpha = -0.22$, $\gamma = -0.17$; 6/6 variants negative) relative to the full dataset ($\alpha = -0.03$, $\gamma = -0.04$; 5/6 negative), consistent with cover stratification concentrating the configuration signal independently of continental composition.

#### S2.1 Full-dataset suppressor diagnostics

Tables in this section report the full variant-level SC, BW, and semi-partial $R^2$ for both predictors across all parameterisations (M1–M4) and all 18 analytical variants. Continental subset suppressor diagnostics are in Section S2.2.

##### S2.1.1 Full dataset — all 18 variants

Tables S2.2–S2.7 report the full variant-level SC, BW, and semi-partial $R^2$ for both predictors across all 18 analytical variants and all four parameterisations (M1–M4), for α, β, and γ diversity. The tables confirm the SC–BW mismatch documented in the main text is consistent across all variants and diversity components. β diversity is included for completeness but is not interpreted under the directed hierarchy.

Across α and γ diversity, the full variant-level tables confirm the same pattern summarised in the main text. A near-zero additive coefficient is not self-interpreting: the full SC–BW–$R^2$sp mismatch below shows that habitat amount control is necessary but not sufficient for fragmentation-per-se inference when predictors share a dominant nonlinear gradient. Large negative fragmentation structure coefficients in M1 (median SC = −0.72), near-zero fragmentation beta weights under the additive specification (median BW = −0.01 for α, −0.020 for γ), partial correction in M2, and directional alignment in M3 (median BW = −0.11 for α, −0.101 for γ; 6/6 variants negative) constitute the suppressor signature under the shared-gradient geometry documented in S2. The discrepancy between the inclusive variance ($SC^2 \approx 0.53$, i.e. fragmentation's total alignment with the fitted gradient) and the unique variance contribution (semi-partial $R^2 \approx 0.01\%$) for fragmentation in M1 is approximately 5,000-fold (using rounded Table 1 values; the main text reports ~8,800-fold based on unrounded pipeline values): fragmentation aligns strongly with the fitted diversity gradient but contributes negligible independent variance once habitat amount is included. The tight confidence intervals around near-zero BW values across all 18 analytical variants are structural suppressor output — the predicted output of a stable geometric constraint operating consistently across parameterisations — not evidence of ecological precision. β diversity showed no consistent directional pattern and is reported in full. All 12 tables are presented as full/subset pairs (a/b) to allow direct comparison across the two sample sizes.

***Table S2.2***. *α diversity — fragmentation predictor. Rows: six analytical variants (Obs = observed richness; q = 0 and q = 2 = Hill numbers). Columns: structure coefficient (SC), standardised beta weight (BW), and semi-partial R² (R²sp) for M1 (additive baseline), M2 (fragmentation residualised on amount), and M3 (amount residualised on fragmentation). Point estimates only.*

| | **M1 (additive baseline)** | | | **M2 (resFrag + Amount)** | | | **M3 (Patches + resAmount)** | | |
|---|---|---|---|---|---|---|---|---|---|
| **Variant** | **SC** | **BW** | **R²sp (%)** | **SC** | **BW** | **R²sp (%)** | **SC** | **BW** | **R²sp (%)** |
| Obs, all pairs | -0.91 | -0.07 | 0.19 | -0.46 | -0.06 | 0.34 | -1.00 | -0.13 | 1.60 |
| Obs, close pairs | -0.73 | -0.02 | 0.00 | -0.20 | -0.03 | 0.08 | -0.80 | -0.12 | 0.82 |
| q = 0, all pairs | -0.61 | 0.01 | 0.01 | -0.03 | 0.00 | 0.00 | -0.85 | -0.06 | 0.25 |
| q = 0, close pairs | -0.67 | 0.00 | 0.00 | -0.11 | -0.01 | 0.01 | -0.88 | -0.07 | 0.31 |
| q = 2, all pairs | -0.72 | -0.01 | 0.00 | -0.20 | -0.03 | 0.06 | -0.91 | -0.11 | 0.89 |
| q = 2, close pairs | -0.77 | -0.03 | 0.01 | -0.26 | -0.04 | 0.12 | -0.94 | -0.12 | 1.08 |

***Table S2.3***. *α diversity — habitat amount predictor. Rows: six analytical variants (Obs = observed richness; q = 0 and q = 2 = Hill numbers). Columns: structure coefficient (SC), standardised beta weight (BW), and semi-partial R² (R²sp) for M1 (additive baseline), M2 (fragmentation residualised on amount), and M3 (amount residualised on fragmentation). Point estimates only.*

| | **M1 (additive baseline)** | | | **M2 (resFrag + Amount)** | | | **M3 (Patches + resAmount)** | | |
|---|---|---|---|---|---|---|---|---|---|
| **Variant** | **SC** | **BW** | **R²sp (%)** | **SC** | **BW** | **R²sp (%)** | **SC** | **BW** | **R²sp (%)** |
| Obs, all pairs | 0.92 | 0.07 | 0.00 | 0.89 | 0.12 | 1.33 | -0.02 | -0.00 | 0.02 |
| Obs, close pairs | 1.00 | 0.15 | 0.52 | 0.98 | 0.16 | 2.42 | 0.59 | 0.09 | 0.23 |
| q = 0, all pairs | 1.00 | 0.10 | 0.29 | 1.00 | 0.09 | 0.84 | 0.53 | 0.04 | 0.02 |
| q = 0, close pairs | 1.00 | 0.09 | 0.22 | 0.99 | 0.09 | 0.83 | 0.48 | 0.04 | 0.00 |
| q = 2, all pairs | 1.00 | 0.13 | 0.41 | 0.98 | 0.14 | 1.85 | 0.41 | 0.05 | 0.00 |
| q = 2, close pairs | 0.99 | 0.12 | 0.29 | 0.96 | 0.14 | 1.83 | 0.34 | 0.04 | 0.00 |

***Table S2.4***. *β diversity — fragmentation predictor. Rows: six analytical variants (Obs = observed richness; q = 0 and q = 2 = Hill numbers). Columns: structure coefficient (SC), standardised beta weight (BW), and semi-partial R² (R²sp) for M1 (additive baseline), M2 (fragmentation residualised on amount), and M3 (amount residualised on fragmentation). Point estimates only.*

| | **M1 (additive baseline)** | | | **M2 (resFrag + Amount)** | | | **M3 (Patches + resAmount)** | | |
|---|---|---|---|---|---|---|---|---|---|
| **Variant** | **SC** | **BW** | **R²sp (%)** | **SC** | **BW** | **R²sp (%)** | **SC** | **BW** | **R²sp (%)** |
| Obs, all pairs | 0.96 | 0.24 | 3.09 | 0.86 | 0.16 | 2.44 | 0.86 | 0.20 | 4.21 |
| Obs, close pairs | 0.64 | 0.18 | 1.85 | 0.99 | 0.12 | 1.58 | 0.82 | 0.08 | 0.78 |
| q = 0, all pairs | -0.99 | -0.05 | 0.12 | -0.13 | 0.00 | 0.00 | -0.84 | -0.05 | 0.30 |
| q = 0, close pairs | -0.84 | -0.01 | 0.01 | 0.20 | 0.01 | 0.01 | -0.76 | -0.03 | 0.05 |
| q = 2, all pairs | -0.86 | -0.04 | 0.06 | -0.56 | -0.07 | 0.39 | -0.79 | -0.08 | 0.42 |
| q = 2, close pairs | -0.95 | -0.06 | 0.16 | -0.66 | -0.07 | 0.43 | -0.66 | -0.07 | 0.18 |

***Table S2.5.*** *β diversity — habitat amount predictor. Rows: six analytical variants (Obs = observed richness; q = 0 and q = 2 = Hill numbers). Columns: structure coefficient (SC), standardised beta weight (BW), and semi-partial $R^2$ ($R^2$sp) for M1 (additive baseline), M2 (fragmentation residualised on amount), and M3 (amount residualised on fragmentation). Point estimates only.*

| | **M1 (additive baseline)** | | | **M2 (resFrag + Amount)** | | | **M3 (Patches + resAmount)** | | |
|---|---|---|---|---|---|---|---|---|---|
| **Variant** | **SC** | **BW** | **$R^2$sp (%)** | **SC** | **BW** | **$R^2$sp (%)** | **SC** | **BW** | **$R^2$sp (%)** |
| Obs, all pairs | -0.45 | 0.07 | 0.43 | -0.51 | -0.09 | 0.75 | 0.51 | 0.12 | 1.67 |
| Obs, close pairs | 0.14 | 0.14 | 1.32 | 0.13 | 0.02 | 0.04 | 0.58 | 0.06 | 0.45 |
| q = 0, all pairs | 0.58 | -0.01 | 0.01 | 0.99 | 0.03 | 0.07 | -0.54 | -0.03 | 0.16 |
| q = 0, close pairs | 0.97 | 0.03 | 0.01 | 0.98 | 0.04 | 0.16 | 0.64 | 0.03 | 0.03 |
| q = 2, all pairs | 0.96 | 0.07 | 0.07 | 0.83 | 0.10 | 0.92 | 0.61 | 0.07 | 0.17 |
| q = 2, close pairs | 0.87 | 0.04 | 0.00 | 0.75 | 0.08 | 0.58 | 0.75 | 0.08 | 0.38 |

***Table S2.6****. γ diversity — fragmentation predictor. Rows: six analytical variants (Obs = observed richness; q = 0 and q = 2 = Hill numbers). Columns: structure coefficient (SC), standardised beta weight (BW), and semi-partial R² (R²sp) for M1 (additive baseline), M2 (fragmentation residualised on amount), and M3 (amount residualised on fragmentation). Point estimates only.*

| | **M1 (additive baseline)** | | | **M2 (resFrag + Amount)** | | | **M3 (Patches + resAmount)** | | |
|---|---|---|---|---|---|---|---|---|---|
| **Variant** | **SC** | **BW** | **R²sp (%)** | **SC** | **BW** | **R²sp (%)** | **SC** | **BW** | **R²sp (%)** |
| Obs, all pairs | -0.92 | -0.06 | 0.14 | -0.51 | -0.05 | 0.27 | -1.00 | -0.10 | 0.89 |
| Obs, close pairs | -0.66 | 0.00 | 0.00 | -0.11 | -0.02 | 0.02 | -0.77 | -0.11 | 0.59 |
| q = 0, all pairs | -0.69 | 0.00 | 0.00 | -0.06 | -0.01 | 0.00 | -0.92 | -0.07 | 0.35 |
| q = 0, close pairs | -0.71 | -0.01 | 0.00 | -0.10 | -0.01 | 0.00 | -0.88 | -0.07 | 0.36 |
| q = 2, all pairs | -0.79 | -0.04 | 0.03 | -0.32 | -0.05 | 0.23 | -0.92 | -0.13 | 1.27 |
| q = 2, close pairs | -0.81 | -0.04 | 0.06 | -0.34 | -0.06 | 0.28 | -0.90 | -0.13 | 1.28 |

***Table S2.7****. γ diversity — habitat amount predictor. Rows: six analytical variants (Obs = observed richness; q = 0 and q = 2 = Hill numbers). Columns: structure coefficient (SC), standardised beta weight (BW), and semi-partial $R^2$ ($R^2$sp) for M1 (additive baseline), M2 (fragmentation residualised on amount), and M3 (amount residualised on fragmentation). Point estimates only.*

| | **M1 (additive baseline)** | | | **M2 (resFrag + Amount)** | | | **M3 (Patches + resAmount)** | | |
|---|---|---|---|---|---|---|---|---|---|
| **Variant** | **SC** | **BW** | **$R^2$sp (%)** | **SC** | **BW** | **$R^2$sp (%)** | **SC** | **BW** | **$R^2$sp (%)** |
| Obs, all pairs | 0.91 | 0.05 | 0.00 | 0.86 | 0.09 | 0.77 | 0.06 | 0.01 | 0.00 |
| Obs, close pairs | 1.00 | 0.15 | 0.68 | 0.99 | 0.15 | 2.25 | 0.64 | 0.09 | 0.30 |
| q = 0, all pairs | 1.00 | 0.09 | 0.20 | 1.00 | 0.09 | 0.81 | 0.40 | 0.03 | 0.00 |
| q = 0, close pairs | 1.00 | 0.09 | 0.21 | 1.00 | 0.10 | 0.92 | 0.47 | 0.04 | 0.00 |
| q = 2, all pairs | 0.99 | 0.13 | 0.33 | 0.95 | 0.15 | 2.22 | 0.39 | 0.05 | 0.00 |
| q = 2, close pairs | 0.98 | 0.13 | 0.29 | 0.94 | 0.15 | 2.34 | 0.44 | 0.06 | 0.01 |

#### S2.2 Sensitivity and subset analyses

Sensitivity analyses were conducted across multiple dataset subsets to evaluate the robustness of the primary suppressor diagnosis.

All subset analyses were predeclared prior to examining results. Each subset was assigned to one of three evidential tiers based on available sample size, following the rule: subsets with 20 or more studies are eligible for the full SC–BW–semi-partial $R^2$ suppressor tripod; subsets with 10 to 19 studies are treated as secondary sensitivity checks with SC and BW reported only; subsets with fewer than 10 studies are fitted under the same model form as directional/descriptive checks only, with no independent inferential claims and no path models. Table S2.8 summarises all nine subsets with their sample sizes, ecological rationale, the specific objection each addresses, their evidential tier, and what analyses were conducted within each.

**Table S2.8.** Subset analysis design: sample sizes, rationale, objection addressed, evidential tier, and analytical scope.

| Subset | n | Rationale | Objection addressed | Tier / Analyses conducted |
|---|---|---|---|---|
| Full dataset | n=37 | Primary analysis — all studies pooled | — | Tier 1 — full tripod (SC+BW+$R^2$sp); path model |
| High-cover (≥50% cover, both landscapes) | n=17 | Tests whether suppressor geometry requires the low-cover nonlinear gradient tail | Result driven by steep nonlinear tail at low cover | Tier 1 — full tripod; no path model |
| Tropical-only | n=32 | Aggregate climate robustness check — removes the 5 temperate studies as a whole | Result driven by temperate minority | Tier 1 — full tripod; no path model |
| SA continental | n=24 | GS25-parallel continental pooling check; tests suppressor persistence in SA | Suppressor is a South-American pooling artefact | Tier 1 — SC+BW only; $R^2$sp not computed (SA qualifies for Tier 1 by sample size ($n = 24 \geq 20$); $R^2$sp is not reported because the partR2 decomposition did not converge for the continental subset model structure in glmmTMB — SC and BW from glmmTMB constitute the available variance-allocation diagnostics); path model |
| non-SA continental | n=13 | GS25-parallel continental check; tests | Same as SA continental | Tier 2 — SC+BW only (secondary sensitivity); |

| | | | | |
|---|---|---|---|---|
| | | suppressor persistence outside SA | | path model |
| Tropical SA | n=23 | Ecologically refined SA partition — removes Aizen_1994 (temperate SA), correcting non-orthogonality of tropical-only and SA/non-SA splits | Tropical-only and SA/non-SA splits are not fully orthogonal | Tier 1 — full tripod; no path model |
| Tropical non-SA | n=9 | Refined non-SA partition restricted to tropical studies | Same as tropical SA | Tier 3 — directional only; no path model |
| Tropical SA high-cover | n=9 | Tests whether the configuration-associated signal concentrates in high-cover tropical SA — consistent with the extinction-filtering hypothesis. | High-cover result is SA in disguise | Tier 3 — directional only; path model run for consistency with Figures 5--6 but coefficients are directional indicators only given n = 9. |
| Tropical non-SA high-cover | n=7 | Thin directional check — tests non-SA high-cover independently | Same as tropical SA high-cover | Tier 3 — directional only; no path model |

Tropical low-cover cells (SA: n=2; non-SA: n=1) were not analysed — sample sizes do not support mixed models with two fixed effects and a random intercept. Path models were run for the full dataset and the GS25-parallel SA/non-SA split only. A supplementary exploratory tropical SA vs tropical non-SA path model is reported in Section S2.2.3 without inferential claims.

#### S2.2.1 High-cover subset (≥50% habitat amount)

All analyses were repeated within the corrected high-cover subset, defined by a study-level minimum filter requiring both the continuous and fragmented landscape to exceed 50% habitat cover (n = 17 studies, 34 landscapes). The corrected filter preserves complete landscape pairs. The previous landscape-level implementation retained 30 studies (53 landscapes) but broke 19 pairs by retaining only the continuous landscape when the fragmented partner fell below 50%, yielding singleton studies whose random intercepts could not estimate within-study contrasts. The corrected n = 17 produces a stronger fragmentation signal (M3 α BW = −0.122 vs −0.109 in the broken filter), confirming the result is not artefactual. To evaluate whether suppressor attenuation persists under reduced nonlinearity, nonlinear smooth terms were retained in both diagnostic orientations (forward edf = 3.20, $p < 0.001$; reverse edf = 1.99, $p < 0.001$), and linear coupling strengthened within this range ($R^2 = 0.634$ vs. 0.454 in the full dataset). The reverse nonlinear component (0.259) remained 16.2-fold larger than the forward (0.016). The full suppressor signature was preserved across α and γ diversity.

This asymmetry is preserved and amplified relative to the full dataset (16.2× vs. 7.75×; 0.259 vs. 0.243 in the reverse direction). This amplification reflects the ecological constraint underlying the asymmetry: patch number is bounded above by total habitat area, but habitat area is not equivalently constrained by patch count. This directional relationship becomes invisible to additive regression because additive coefficient allocation has no mechanism for representing causal direction; restricting the dataset to landscapes with ≥50% cover reduces but does not resolve the asymmetry.

The subset result also confirms that nonlinearity amplifies suppressor attenuation but is not its source. Within the subset, linear coupling strengthened ($R^2 = 0.634$ vs. 0.454 in the full dataset) while the nonlinear component declined modestly (0.259 vs. 0.243). Suppressor attenuation arises from the shared variance structure of the predictor pair, which is present in the linear correlation; nonlinear curvature intensifies the shared component but does not create it. The implication is general: the problem is not confined to datasets with strongly nonlinear predictor relationships, and metric-selection strategies that reduce nonlinear coupling without achieving predictor separability do not resolve it.

Coupling diagnostics for this subset are shown in Figure S2.5; coefficient patterns (SC and BW for fragmentation and habitat amount, M1–M3) in Figure S2.6.

##### S2.2.1.1 Fragmentation predictor

#### S2.2.2 Tropical-only subset

The GS25–F26 dataset includes 32 tropical and 5 temperate studies, as classified in Gonçalves-Souza et al. 2025 (hereafter GS25; Supplementary Table 7 (Climate column)). To evaluate whether the main inferential conclusions are sensitive to the small non-tropical component, the full diagnostic pipeline was repeated after excluding the five temperate studies (Aizen_1994, Bolger_1997, Fujita_2008, Gaublomme_2008, Gibb_2002). Predictor standardisation, GAM residualisation, and orthogonality verification were rebuilt within the resulting 32-study tropical subset; tropical-subset M3 therefore uses a res_amount_z that is orthogonal to patches within that subset, not the full-dataset residual. All 18 analytical variants (3 diversity indices × 2 pairing designs × 3 species weightings) were refit for M1 and M3.

Table S2.9 summarises the coupling diagnostics and design-reconstruction check for the tropical subset. The linear predictor correlation was stable (r = −0.705 vs −0.674 full dataset). The design-reconstruction check yielded 26/32 study pairs (81.2%) satisfying both directional conditions (fragmented lower amount, fragmented more patches), compared to 31/37 (84%) in the full dataset. These results confirm that the directional predictor structure motivating the main analysis is present in tropical studies independently of the temperate minority.

Table S2.10 summarises the M1 and M3 fragmentation coefficients for the tropical subset across all 6 variants. M1 returned median beta weights of −0.002 (α) and −0.040 (γ) in the tropical subset, comparable to the full-dataset values (−0.014 and −0.020 respectively), confirming that the additive null persists in the dominant tropical portion of the dataset. M3 recovered negative fragmentation coefficients across all 6/6 analytical variants for both α (median = −0.056) and γ (median = −0.069), with full-dataset M3 values of −0.114 and −0.101 respectively; directional comparisons should be interpreted with caution because residualisation was rebuilt within the tropical subset, meaning the estimands are not identical. The qualitative suppressor signature (SC ≠ BW, structure coefficient negative, BW near zero in M1) remained detectable for α diversity (median M1 SC = −0.142 vs −0.725 full dataset) but was weaker for γ (−0.104 vs −0.747), consistent with reduced nonlinear coupling in the smaller tropical-only predictor space. Beta-diversity coefficients are not interpreted as the suppressor pattern was absent in this component in both the full dataset and the tropical subset. Together, these results indicate that the main inferential conclusion — that the additive null does not establish ecological independence, and that the recoverable α and γ direction is negative — is robust to exclusion of the five temperate studies.

***Table S2.9.*** *Predictor coupling and design-reconstruction diagnostics for the tropical-only subset (n = 32 studies). Temperate studies excluded: Aizen_1994, Bolger_1997, Fujita_2008, Gaublomme_2008, Gibb_2002 (classified per GS25 Supplementary Table 7). Reference values for the full 37-study dataset: linear r = −0.674; design reconstruction 31/37 (84%).*

| **Diagnostic** | **Tropical subset (n=32)** | **Full dataset (n=37)** | **Status** |
|---|---|---|---|
| Linear r (amount–patches) | −0.705 | −0.674 | Stable |
| Design reconstruction: both conditions | 26/32 (81.2%) | 31/37 (84%) | Stable |
| Fragmented lower amount | 30/32 (93.8%) | 35/37 (94.6%) | Stable |
| Fragmented more patches | 26/32 (81.2%) | 31/37 (83.8%) | Stable |

***Table S2.10.*** M1 and M3 fragmentation coefficient summary for the tropical-only subset (n = 32 studies, 6 variants per cell). BW = standardised beta weight (median across 6 variants); n− = number of variants with negative BW; SC = structure coefficient (M1 only). Note: M3 tropical-subset residualisation was rebuilt within the 32-study predictor distribution; M3 values are therefore not directly comparable in magnitude to full-dataset M3 values. Directional comparison (sign and n−) is the primary basis for inference.

| ***Component*** | ***M1 BW (trop)*** | ***M1 BW (full)*** | ***M1 SC (trop)*** | ***M3 BW (trop)*** | ***M3 BW (full)*** | ***M3 n−/6*** |
|---|---|---|---|---|---|---|
| α diversity | −0.002 | −0.014 | −0.142 | −0.056 | −0.114 | 6/6 |
| γ diversity | −0.040 | −0.020 | −0.104 | −0.069 | −0.101 | 6/6 |
| β diversity | −0.020 | +0.064 | +0.127 | −0.027 | +0.022 | Inconsistent. |

Coupling diagnostics for the Tropical SA subset are shown in Figure S2.7; coefficient patterns (SC and BW, M1–M3) in Figure S2.8.

#### S2.2.3 Continental subsets (SA vs non-SA)

##### S2.2.3.1 Suppressor diagnostics by continental group

The M1–M4 suppressor diagnostic was repeated separately for the 24 South American (SA) and 13 non-South American (non-SA) studies. GAM coupling diagnostics, residualisation, and mixed models were rebuilt within each group. The SC–BW mismatch (large |SC|, near-zero BW under M1) was present in both SA and non-SA, consistent with the full-dataset suppressor pattern not being an artefact of continental pooling: the SC–BW mismatch under M1 was independently present within both continental subsets (full dataset: SC/BW ratio ≈51×; SA: ≈20×; non-SA: ≈4×), confirming that pooling amplifies the mismatch but does not create it (noting that non-SA is a Tier 2 sensitivity check without semi-partial $R^2$ confirmation). Three features of the within-group geometry require explicit statement.

First, the predictor coupling asymmetry that drives M3 orthogonality in the full dataset (7.75× forward nonlinear component) does not generalise within either subset. In SA the directional nonlinear dependence reverses: GAM forward $R^2$ = 0.430 (patches ∼ s(amount)), reverse = 0.760 (amount ∼ s(patches)); in non-SA the asymmetry is weaker (forward = 0.642, reverse = 0.577). Second, M3 orthogonality was satisfied within both continental subsets (SA: residual GAM $R^2$ = 0.000/0.000, criterion met; non-SA: 0.000/0.000, criterion met). The orthogonality criterion (both residual GAM $R^2 < 0.02$) was not satisfied in all subsets: the tropical-only subset (n = 32) produced M3 reverse residual coupling of 0.032 (marginally above the 0.02 threshold) and the Tropical non-SA high-cover subset (n = 7) produced 0.193, a clear failure at Tier 3. Subset-level M3 coefficients are therefore interpretable within the two continental strata, and directional indicators only in the two boundary-condition subsets. Third, SA shows a diagnostically anomalous M4 result: fragmentation coefficients remain consistently negative under double residualisation (median α = −0.051, 6/6 variants), while non-SA returns near-zero M4 values as expected for a negative control (median α = +0.032). The expected cancellation under double residualisation did not occur in SA; this diagnostic anomaly is reported in Section S2.2 and discussed in the main text. Semi-partial $R^2$ is not reported for continental subsets; BW and SC from glmmTMB are the available variance-allocation outputs for this analysis.

Table S2.11. M1–M4 fragmentation BW and SC by continental group across all 18 analytical variants. Columns: group (SA/non-SA), parameterisation (M1–M4), diversity component, variant, median BW [range], n negative/6, median SC (M1 only). Semi-partial $R^2$ is not reported for the SA continental subset; BW and SC from glmmTMB constitute the available variance-allocation diagnostics for both continental groups (see S2.2.1 for rationale).

#### S2.2.3.2 Habitat-amount overlap, biome contingency, and power

The continental contrast in path coefficients cannot be attributed to differences in within-landscape habitat cover. The two groups were statistically indistinguishable in within-landscape habitat amount (SA: mean 67.0%, median 64.0%, SD 17.1%, range 38.8–98.4%; non-SA: mean 62.9%, median 67.3%, SD 21.9%, range 34.5–94.3%; Welch $t = 0.55$, $p = 0.587$). Ten of twelve non-SA studies fall within the SA amount range. Neither group includes studies with mean cover below 35%, ruling out classical extinction filtering at the 20–30% threshold. The continental contrast in Frag→Div path coefficients is therefore not an artefact of differential habitat loss among the sampled landscapes.

Four clarifications about the evidential status of this pattern are important. First, it is not classical extinction filtering at within-landscape cover thresholds: no study in either group falls below the 20–30% threshold at which sensitive species are typically lost, and cover distributions overlap substantially. Second, it is not a habitat amount effect: the groups are statistically indistinguishable within-landscape amount (see habitat-amount overlap results above). Third, it is not proven: the SA Frag→Div point estimates are consistent and directionally coherent across 6/6 $\alpha$ and 6/6 $\gamma$ variants, but confidence intervals are wide at current sample sizes, and the pattern requires prospective confirmation. Fourth, it is not noise: the consistency across 6/6 variants, convergence with a dedicated Atlantic Forest path analysis (Püttker et al. 2020), and internal coherence across metrics and pairing designs distinguish the SA pattern from sampling artefact.

Power to detect the SA Frag→Div observed richness effect ($\beta \approx -0.21$) at $n = 48$ SA landscape pairs is approximately 18–19%, meaning the study would fail to return $p < 0.05$ roughly four times in five even if this were the true effect. The 95% confidence intervals ($\approx -0.60$ to $+0.18$) indicate limited precision rather than evidence of no effect: they also exclude the possibility of ruling out ecologically substantial effects in the −0.20 to −0.40 range. Approximately 325–336 SA landscape pairs would be required for 80% power and 435–450 for 90% power at this effect magnitude. These requirements should guide future within-biome study design.

The biome-contingency hypothesis proposed here has four testable conditions: (1) biome-level deforestation exceeding 80% of original extent filters the regional species pool to forest specialists for whom configuration is a genuine ecological constraint; (2) hard, high-contrast matrix amplifies functional isolation even at moderate fragment distances; (3) long fragmentation histories (>50–100 years) allow extinction debts to manifest in observed richness, making the configuration signal more recoverable; (4) at high within-landscape cover (>85%), amount variation is near-saturated and configuration becomes the dominant residual axis of diversity variation. The Atlantic Forest satisfies all four conditions. Many non-SA landscapes in this dataset satisfy one or two. Prospective studies stratified by fragmentation history and matrix contrast would allow explicit testing of which conditions are necessary and which are sufficient for configuration effects to be statistically recoverable.

#### S2.2.4 Crossed tropical sensitivity checks

Crossed tropical sensitivity checks confirmed that the orthogonality criterion was satisfied within both full continental subsets (S2.2.3); one crossed tropical cell failed outright (Tropical non-SA high-cover, n = 7; reverse residual coupling = 0.193). After restricting both groups to tropical studies only, one of the four crossed tropical cells satisfied the M3 orthogonality criterion at adequate sample size: Tropical SA (n = 23). Two further cells — Tropical non-SA (n = 9) and Tropical SA high-cover (n = 9; Tier 3) — returned 0.000/0.000 residual GAM values, but at that sample size the GAM lacks leverage to detect nonlinear structure; those values reflect estimation failure rather than confirmed orthogonality, and both cells are retained as Tier 3 directional indicators on sample-size grounds. In all four tropical cells, the forward nonlinear coupling component is near zero, so the shared predictor structure is predominantly linear in the forward direction (r = −0.68 to −0.89). The crossed tropical M3 coefficients are directional indicators only, not orthogonalised estimates, consistent with their Tier 1/3 classification.

In tropical South American studies (n = 23), the M1 SC–BW mismatch persisted (SC = −0.869, BW = −0.049, 6/6 $\alpha$ variants negative) and fragmentation coefficients became more negative under M3 (median BW = −0.112, 6/6 negative); the Tropical SA high-cover cell (n = 9; Tier 3, directional indicators only) showed directional consistency (M1 BW = −0.088; M3 BW = −0.060, 6/6 negative) but no intensification relative to the full Tropical SA result.

In tropical non-South American studies (n = 9; directional indicators only), the additive model yielded a positive fragmentation coefficient (M1 BW = +0.289, SC = +0.257, 0/6 negative) that reversed sign under M3 (BW = −0.205, 6/6 negative), revealing that the additive model allocates the shared gradient differently in this subgroup than in tropical SA, and that pooled additive estimates obscure sharply divergent context-specific allocation patterns. Because the tropical non-South-American cell is thin (n = 9) and semi-partial $R^2$ is not reported for these cells, these patterns are reported as directional observations consistent with the heterogeneity argument and are not used as independent inferential anchors.

Coupling diagnostics for the Tropical SA ≥50% cover subset are shown in Figure S2.9. The 0.000/0.000 residual GAM values in this cell reflect GAM leverage failure at n = 9, not verified orthogonality; the cell is retained as a Tier 3 directional indicator on sample-size grounds.

#### S2.2.5 Ecological conditionality

The suppressor geometry documented in the main text is the primary reason the F26 additive null coefficient cannot be uniquely interpreted as evidence of ecological independence. A second, independent reason also applies, and it operates even if predictor separability were verified: fragmentation effects are context-dependent in ways that cause pooled multi-study estimates to average over opposing ecological signals, leaving aggregate coefficients non-adjudicative regardless of predictor geometry. In the Neotropical vertebrate assemblage that dominates the GS25/F26 dataset, feeding guild and forest dependency are the strongest trait-level predictors of fragmentation sensitivity — insectivores and obligate forest species show consistently higher proportions of negative responses — and those authors note explicitly that fragmentation in the Neotropics always involves habitat loss, independently confirming the structural coupling the main text diagnoses statistically (Vetter et al. 2011). At the community level, interactions between rarity and ecological specialisation produce synergistic rather than additive extinction risk in fragments, so pooled analyses averaging across species with varying trait profiles systematically underestimate sensitivity in the subset of taxa where the isolating conditions for fragmentation effects are actually met (Henle et al. 2004). The type of land cover surrounding patches is among the strongest moderators of sensitivity to patch area and isolation — stronger than taxon, diet, dispersal, or spatial scale — and where organisms use the intervening matrix, patches can be functionally larger and more connected than their geometric delineation implies, so weak or inconsistent area and isolation coefficients need not indicate that habitat structure is ecologically irrelevant (Prugh et al. 2008). Multi-taxa synthesis confirms this: matrix quality can, for several reviewed taxa, outweigh spatial configuration as a direct predictor of biodiversity (Arroyo-Rodríguez et al. 2020).

These two sources of ambiguity — predictor-geometry indeterminacy and ecological context-dependence — are logically independent but compound in multi-study syntheses spanning heterogeneous biomes and land-use histories. Predictor geometry operates at the level of statistical architecture: even if the underlying ecology were perfectly heterogeneous, the suppressor signature in M1 arises from the shared habitat-loss gradient, not from the biological signal itself. Ecological context-dependence operates at the level of the biological signal: even a perfectly orthogonal predictor design faces a pooled estimate that averages context-specific effects with opposing directions across heterogeneous systems. Both sources would independently produce a non-adjudicative result in a multi-study synthesis; in this dataset they compound. Explicitly testing the ecological-conditionality hypothesis requires within-biome designs with sufficient replication to separate predictor-geometry effects from biome-specific ecological mechanisms — a distinct research programme from the inferential-standard question the main text addresses, and one that requires purpose-built study designs rather than reanalysis of existing data.

#### S2.3 Path model — causal hierarchy check

The path model was fitted as a complementary causal-hierarchy check, not as an independent test of fragmentation effect sizes. The structural path (Amount → Fragmentation) formalises the directed coupling documented in Section S2.2 as a prior in the model specification (Ruffell et al. 2016; Grace and Irvine 2020; Püttker et al. 2020). Path coefficients use 1-SD standardisation throughout (not the 2-SD standardisation used for M1–M4 beta weights; see main text Methods). The model is saturated (Fisher's $C = 0$, $df = 0$, $p = 1.000$ by construction; confirmed empirically). Full numerical outputs for all 18 variants are in Sections S2.3.1–S2.3.5.

##### S2.3.1 Full 18-variant results

A simple path model was fitted to test whether a causal hierarchy in which habitat amount drives configuration change is consistent with the GS25–F26 dataset, and whether the resulting path coefficients are consistent with the main GLMM diagnostics. The causal direction (Amount → Fragmentation) was specified as a theoretical prior, justified by the algebraic identity $\bar{A}p = Al/Np$ and the 7.75× nonlinear coupling asymmetry; it was not inferred from the same data (Grace and Irvine 2020; Ruffell et al. 2016). Two mixed models were fitted per variant on a clean four-column data frame (refshort, response, frag_z, amount_z) to prevent piecewiseSEM from accessing NA-named columns: a structural path frag_z ~ amount_z + (1|refshort), encoding the directed coupling prior; and a response path diversity ~ frag_z + amount_z + (1|refshort), matching the M1 specification. Standardised path coefficients were computed as raw coefficient × (SD predictor / SD response). The indirect effect of habitat amount on diversity via fragmentation was computed as the product of the two standardised coefficients; its approximate standard error was derived via the delta method: $SE_{indirect} = \sqrt{(\beta_{fd} \times SE_{af})^2 + (\beta_{af} \times SE_{fd})^2}$. The model is saturated (3 observed variables, all paths present), so Fisher's C = 0, df = 0, p = 1 by construction (Shipley 2000, 2009). Path coefficients were verified using piecewiseSEM (Lefcheck 2016), which confirmed coefficient equivalence and returned marginal $R^2 = 0.460$ for the structural path (Amount → Frag) and marginal $R^2 = 0.002$–$0.039$ (conditional $R^2 = 0.32$–$0.99$) for the response path across all 18 variants (Table S2.12). Path coefficients use conventional 1-SD standardisation (raw coefficient × SD predictor / SD response), not the 2-SD standardisation used for M1–M4 beta weights (Gelman 2008); these scales are not directly comparable. All 18 analytical variants were fitted (3 diversity indices × 2 pairing designs × 3 species weightings). To evaluate whether path coefficients were consistent across the dataset's main continental grouping, path models were additionally fitted separately for the 24 South American and 13 non-South American studies (continental classification from GS25 Supplementary Table 7), following the continental sensitivity approach used by GS25 in their own Extended Data Fig. 10 analysis.

The structural path Amount → Fragmentation was strongly negative and invariant across all 18 variants (standardised coefficient = −0.688 in every case), confirming that habitat amount robustly predicts configuration regardless of how biodiversity is measured. Table S2.12 summarises all path coefficients. The direct Fragmentation → Diversity path was weak and imprecise in all variants (no significant p-values); medians across the 6 α- and γ-diversity variants were −0.029 and −0.041 respectively, with the path negative in 4/6 α-variants and 5/6 γ-variants. This near-zero direct path mirrors M1 and is expected: the SEM uses the same additive response specification as M1 and does not residualise the shared gradient; suppressor attenuation therefore persists. The indirect effect of Amount on Diversity via Fragmentation was positive for α and γ in all-pairs designs (α: +0.092, SE ≈ 0.110; γ: +0.078, SE ≈ 0.108, observed richness; median across all 6 α-variants: +0.020; median γ: +0.028), indicating that part of habitat amount's biodiversity benefit is routed through reduced fragmentation under the causal hierarchy. These indirect effects are directionally consistent with mediation but imprecisely estimated; full-gradient path models are expected to attenuate configuration paths relative to analyses conducted within habitat-amount strata, because the shared amount gradient continues to suppress the fragmentation coefficient under the full observational range. Observed β-diversity path coefficients are reported in Table S2.12 but are not interpreted under the directed Amount → Fragmentation hypothesis; the positive observed β Fragmentation → Diversity path is consistent with the known opposing β-diversity dynamic documented in the main analysis.

Residual diagnostics (Shapiro–Wilk normality test on Pearson residuals from each component model) confirmed that response path residuals were approximately normal across all 18 variants (W = 0.944–0.994, p = 0.003–0.971; the three variants with $p < 0.05$ all showed |skewness| < 0.17, indicating minor departures). Structural path residuals were severely non-normal in all variants (W = 0.742, p ≈ 0, skewness = 2.40), driven by the right-skewed distribution of raw patch counts (landscape-level skewness = 2.1). This does not affect the directional conclusion — the Amount → Fragmentation coefficient of −0.688 is invariant across all 18 variants — but the standard error of that coefficient may be anti-conservative. A robustness check replacing raw patch count with log-transformed patch count (log(number_patches), 2-SD standardised) confirmed: (1) structural path residuals normalised substantially ($W = 0.965$, $p = 0.036$, skewness = −0.42); (2) the directional structure was preserved — Amount → log(Frag) strongly negative ($\beta = -0.784$), direct Frag → Diversity path near-zero and negative (α: $\beta = -0.231$, $p = 0.171$; γ: $\beta = -0.182$, $p = 0.277$), indirect effect positive (α: +0.181, SE = 0.132; γ: +0.143, SE = 0.131); (3) the coupling magnitude is metric-conditional — Amount → log(Frag) is stronger than Amount → raw(Frag) (−0.784 vs −0.688), because the log-linear identity $\log(N_p) = \log(A_l) - \log(\bar{A}_p)$ linearises the amount–configuration relationship. This confirms that metric choice shapes predictor geometry and therefore suppressor severity: studies should treat metric selection as an identifiability decision and conduct sensitivity analyses across metric transformations before interpreting additive fragmentation coefficients from observational data. The main analysis uses raw patch count throughout, consistent with F26.

#### S2.3.2 Continental sensitivity

The path model and M3 address different questions that habitat amount structures configuration. The path model retains the observed amount–configuration coupling and asks whether a hierarchical path model is compatible with the data; in this dataset it confirms a strong Amount → Fragmentation relationship but recovers only a weak direct Fragmentation → Diversity path, because the response path specification is identical to M1 and the shared gradient therefore continues to attenuate the fragmentation coefficient. M3 addresses a different estimand by removing the amount-structured component of fragmentation via nonlinear GAM residualisation (verified near-zero residual coupling, GAM $R^2 < 0.02$) and estimating the biodiversity association of the amount-orthogonalised configuration signal. Read together, the two parameterisations show that the observed additive null is compatible with strong hierarchical coupling, and that a negative configuration-associated direction becomes clearer once the shared gradient is removed. The contrast between the Frag → Diversity path in the path model and the fragmentation beta weight in M3 shows how strongly the observed configuration signal depends on shared-gradient structure: the path model returns a near-zero direct path (median $\alpha = -0.029$) consistent with M1 (−0.014), while M3 recovers a consistently negative direction (median $\alpha = -0.114$). These are not alternative estimates of the same quantity; they are estimates of different, complementary quantities under different predictor representations of the same ecological system.

***Table S2.12.*** *Path model coefficients for the global dataset, all analytical variants, α and γ diversity. Two groups (n and β Amt→Frag shown in group headers): full dataset (n = 37; β Amt→Frag = −0.688) and high-cover subset (n = 17, both landscapes ≥50%; β Amt→Frag = −0.796). Response path: Frag → Diversity and Amount → Diversity. All coefficients standardised (1-SD). Indirect = β_amt_frag × β_frag_div; SE(Indirect) via delta method. β diversity available in block_sem_results.csv. * p < 0.05; ** p < 0.01.*

| Div | Variant | β Frag→Div | p (Frag→Div) | β Amt→Div | p (Amt→Div) | Indirect | SE (Indirect) |
|---|---|---|---|---|---|---|---|
| *Global (all) (n = 37; β Amt→Frag = -0.688)* | | | | | | | |
| α | Obs, all pairs | -0.133 | 0.404 | +0.144 | 0.326 | +0.092 | 0.110 |
| α | Obs, close pairs | -0.032 | 0.807 | +0.292 | 0.016* | +0.022 | 0.089 |
| α | q=0, all pairs | +0.019 | 0.777 | +0.195 | 0.002** | -0.013 | 0.046 |
| α | q=0, close pairs | -0.000 | 0.999 | +0.182 | 0.004** | +0.000 | 0.047 |
| α | q=2, all pairs | -0.027 | 0.846 | +0.255 | 0.051 | +0.019 | 0.096 |
| α | q=2, close pairs | -0.054 | 0.701 | +0.236 | 0.071 | +0.037 | 0.097 |
| γ | Obs, all pairs | -0.113 | 0.474 | +0.102 | 0.481 | +0.078 | 0.108 |
| γ | Obs, close pairs | +0.007 | 0.962 | +0.305 | 0.019* | -0.005 | 0.096 |
| γ | q=0, all pairs | -0.005 | 0.946 | +0.175 | 0.007** | +0.003 | 0.048 |
| γ | q=0, close pairs | -0.012 | 0.876 | +0.182 | 0.009** | +0.008 | 0.051 |
| γ | q=2, all pairs | -0.071 | 0.649 | +0.253 | 0.081 | +0.049 | 0.107 |
| γ | q=2, close pairs | -0.085 | 0.585 | +0.251 | 0.083 | +0.059 | 0.107 |
| *Global (≥50%) (n = 17; β Amt→Frag = -0.796)* | | | | | | | |
| α | Obs, all pairs | -0.294 | 0.308 | +0.358 | 0.323 | +0.234 | 0.228 |
| α | Obs, close pairs | -0.206 | 0.448 | +0.225 | 0.508 | +0.164 | 0.214 |
| α | q=0, all pairs | -0.060 | 0.626 | +0.183 | 0.241 | +0.048 | 0.097 |
| α | q=0, close pairs | -0.067 | 0.581 | +0.185 | 0.229 | +0.053 | 0.096 |
| α | q=2, all pairs | -0.249 | 0.458 | +0.141 | 0.736 | +0.198 | 0.265 |
| α | q=2, close pairs | -0.232 | 0.491 | +0.206 | 0.625 | +0.185 | 0.266 |
| γ | Obs, all pairs | -0.190 | 0.528 | +0.168 | 0.656 | +0.151 | 0.238 |
| γ | Obs, close pairs | -0.147 | 0.573 | +0.192 | 0.557 | +0.117 | 0.206 |
| γ | q=0, all pairs | -0.087 | 0.464 | +0.148 | 0.323 | +0.069 | 0.094 |
| γ | q=0, close pairs | -0.106 | 0.412 | +0.176 | 0.280 | +0.085 | 0.102 |
| γ | q=2, all pairs | -0.335 | 0.350 | +0.039 | 0.930 | +0.266 | 0.283 |
| γ | q=2, close pairs | -0.324 | 0.367 | +0.051 | 0.909 | +0.258 | 0.284 |

***Table S2.13.*** *Path model coefficients for Tropical SA groups, all analytical variants, α and γ diversity. Two groups (n and β Amt→Frag shown in group headers): Tropical SA full coverage range (n = 23; β Amt→Frag = −0.702) and Tropical SA high-cover subset (n = 9, both landscapes ≥50%; β Amt→Frag = −0.787; Tier 3 — directional indicators only). Response path: Frag → Diversity and Amount → Diversity. All coefficients standardised (1-SD). No p-values reach α = 0.05 in this table.*

| Div | Variant | β Frag→Div | p (Frag→Div) | β Amt→Div | p (Amt→Div) | Indirect | SE (Indirect) |
|---|---|---|---|---|---|---|---|
| *Tropical SA (all) (n = 23; β Amt→Frag = -0.702)* | | | | | | | |
| α | Obs, all pairs | -0.221 | 0.172 | -0.109 | 0.506 | +0.155 | 0.114 |
| α | Obs, close pairs | -0.114 | 0.346 | +0.117 | 0.345 | +0.080 | 0.085 |
| α | q=0, all pairs | -0.014 | 0.828 | +0.089 | 0.192 | +0.010 | 0.046 |
| α | q=0, close pairs | -0.035 | 0.604 | +0.076 | 0.267 | +0.024 | 0.047 |
| α | q=2, all pairs | -0.039 | 0.755 | +0.096 | 0.461 | +0.028 | 0.088 |
| α | q=2, close pairs | -0.063 | 0.615 | +0.078 | 0.546 | +0.044 | 0.088 |
| γ | Obs, all pairs | -0.222 | 0.172 | -0.163 | 0.323 | +0.156 | 0.115 |
| γ | Obs, close pairs | -0.091 | 0.495 | +0.113 | 0.411 | +0.064 | 0.094 |
| γ | q=0, all pairs | -0.047 | 0.537 | +0.073 | 0.347 | +0.033 | 0.053 |
| γ | q=0, close pairs | -0.058 | 0.453 | +0.068 | 0.388 | +0.041 | 0.054 |
| γ | q=2, all pairs | -0.117 | 0.335 | +0.008 | 0.948 | +0.083 | 0.085 |
| γ | q=2, close pairs | -0.143 | 0.266 | +0.000 | 0.999 | +0.100 | 0.090 |
| *Tropical SA (≥50%) (n = 9; β Amt→Frag = -0.787)* | | | | | | | |
| α | Obs, all pairs | -0.428 | 0.140 | -0.343 | 0.421 | +0.337 | 0.226 |
| α | Obs, close pairs | -0.329 | 0.218 | -0.098 | 0.804 | +0.259 | 0.208 |
| α | q=0, all pairs | -0.064 | 0.533 | -0.028 | 0.857 | +0.050 | 0.079 |
| α | q=0, close pairs | -0.083 | 0.422 | -0.029 | 0.851 | +0.065 | 0.080 |
| α | q=2, all pairs | -0.098 | 0.742 | -0.027 | 0.953 | +0.077 | 0.231 |
| α | q=2, close pairs | -0.116 | 0.691 | -0.028 | 0.949 | +0.092 | 0.226 |
| γ | Obs, all pairs | -0.331 | 0.249 | -0.550 | 0.207 | +0.261 | 0.223 |
| γ | Obs, close pairs | -0.264 | 0.299 | -0.040 | 0.916 | +0.208 | 0.197 |
| γ | q=0, all pairs | -0.111 | 0.338 | -0.072 | 0.676 | +0.087 | 0.090 |
| γ | q=0, close pairs | -0.131 | 0.187 | -0.054 | 0.711 | +0.103 | 0.077 |
| γ | q=2, all pairs | -0.132 | 0.635 | -0.112 | 0.789 | +0.104 | 0.216 |
| γ | q=2, close pairs | -0.168 | 0.562 | -0.140 | 0.748 | +0.132 | 0.225 |

The Tropical SA high-cover subset (n = 9, Tier 3; directional indicators only) shows a pattern of negative direct Fragmentation → Diversity and negative direct Amount → Diversity alongside a positive indirect effect (Amount → Fragmentation → Diversity), consistent with Figure 5. The positive indirect effect is a mathematical consequence of the sign combination in the path model: $\beta$_AmtFrag is negative (more cover = fewer patches, the standard coupling direction) and $\beta$_FragDiv is negative (more patches = lower diversity), so their product — the indirect effect — is positive. This is not a paradox: it encodes the conservation-relevant chain in which more habitat amount suppresses fragmentation, and reduced fragmentation increases diversity. The negative direct Amount → Diversity path, which emerges once the fragmentation-mediated pathway is held constant, is tentatively consistent with structural homogenisation at the upper end of the habitat-cover gradient: at very high cover, continuous landscapes may become compositionally homogeneous relative to fragmented counterparts that retain higher within-landscape heterogeneity, reducing $\gamma$ diversity in continuous landscapes once the positive effect of amount operating through reduced fragmentation is accounted for. This interpretation is speculative at n = 9 and is offered as a Tier 3 directional observation; no inferential claims are drawn from it.

The multigroup() function in piecewiseSEM does not support mixed models with random intercepts, precluding a formal test of path coefficient equality between continental groups. The SA vs. non-SA comparison is therefore descriptive. Singular random intercept estimates occurred in small subsets (non-SA, Tropical SA high-cover, Tropical non-SA, Tropical non-SA high-cover), consistent with negligible between-study variance in those subsets; fixed-effect estimates are unaffected. These subsets are reported as Tier 2 or Tier 3 (directional indicators only).

#### S2.3.3 AIC comparison (directed vs correlated specification)

AIC comparison favoured the directed specification over the correlated specification ($\Delta$AIC = −70.8). In the piecewise SEM framework used here, this comparison evaluates whether explicitly modelling the Amount → Fragmentation relationship as a directed structural path improves fit relative to treating the two predictors as freely correlated. The comparison reflects structural path fit only, not variation in the diversity response. The large negative $\Delta$AIC is consistent with habitat amount being a strong predictor of patch number ($\beta$ = −0.69 for the full dataset, n=37), making the directed model a substantially better description of the data-generating structure than the correlated alternative.

#### S2.3.4 $R^2$ and residual diagnostics

The path model showed adequate fit across all 18 analytical variants. Marginal $R^2$ values for the structural path (Amount → Fragmentation) were consistently high (marginal $R^2$ = 0.460, reported in Section S2.3.1), confirming that habitat amount accounts for the large majority of variance in patch number. Conditional $R^2$ for the response paths (Amount + Fragmentation → Diversity) were comparable to those of the corresponding M1 glmmTMB models (Table S2.1). Shapiro-Wilk normality tests on Pearson residuals from the path model component models showed approximate normality for response paths across all 18 variants (W = 0.944–0.994; the three variants with $p < 0.05$ all showed |skewness| < 0.17). Structural path residuals were non-normal in all variants (W = 0.742, $p \approx 0$, skewness = 2.40), driven by right-skewed raw patch counts; this does not affect the directional conclusion. Note: these Shapiro-Wilk diagnostics apply to path model residuals only; DHARMa simulation-based diagnostics for the primary GLMM models (M1–M4) are in Section S2.4.1.

Singular fits occurred in several small subsets (non-SA, n = 13; Tropical SA high-cover, n = 9; Tropical non-SA and Tropical non-SA high-cover, n = 7–9), indicating that the random intercept variance was estimated at or near zero in those subsets — the study-level grouping explained negligible additional variance beyond the fixed effects. Singular fits do not bias fixed-effect estimates but render the random-effects structure uninformative. These subsets are Tier 2 or Tier 3 by design and are reported as directional indicators only; no inferential claims depend on them.

##### S2.3.5 Log-patches robustness

Path model results were qualitatively unchanged when log-transformed patch count replaced raw patch count as the fragmentation metric (α and γ diversity): the sign, direction, and relative magnitude of all direct and indirect paths were preserved. The Amount → Fragmentation structural path remained strongly negative (consistent with the algebraic identity and the coupling asymmetry documented in Supplementary Material S1 (Section S1.2.4)), and the direct Fragmentation → Diversity path remained near zero, confirming that the path model result is not sensitive to the choice of patch count scale.

#### S2.4 Model diagnostics

##### S2.4.1 Residual diagnostics

Residual adequacy was assessed using DHARMa (Hartig 2022; n = 250 simulations per model) for 16 representative models spanning M1–M4, both diversity components (α and γ), and both diversity orders (q = 0 and q = 2), with random intercepts by study as in the main analysis. Global tests (testUniformity, testDispersion, testZeroInflation, testOutliers) and predictor-specific quantile residual tests (plotResiduals against frag_z and amount_z) were applied. Residuals were additionally aggregated per study (recalculateResiduals(group = refshort)) and subjected to testUniformity to detect systematic study-level over- or underprediction. No model showed significant departure in any diagnostic test (0 of 16 models flagged; median KS test p = 0.272; median dispersion = 1.030). Any mild quantile deviations were present across M1, M2, M3, and M4 alike, confirming they reflect the data structure rather than a reparametrisation artefact. The inferential exclusion conclusions drawn from these diagnostics — that coefficient shifts do not arise from differential model adequacy or heterogeneous residual structure — are stated in the main-text Discussion (Limits of inference section).

Levene tests for equality of residual variance between fragmented and continuous landscape types were conducted across all 18 analytical variants and all four parameterisations (M1--M4). No variant returned a significant result (all $p > 0.05$), confirming that residual variance was homogeneous across landscape types and that coefficient shifts across parameterisations do not reflect differential residual heterogeneity.

##### S2.4.2 Random-effect stability

Per-study random intercepts were compared across M1, M2, and M3 to assess whether reparametrisation transfers variance into the study-level random effect. Random intercept standard deviations were stable: 0.867 (M1), 0.865 (M2), 0.861 (M3) — a range of 0.006 across parameterisations (0.7% relative change). Intercept shifts from M1 to M2 were small in absolute magnitude (range −0.026 to +0.031) and shifts from M1 to M3 were similarly limited (−0.084 to +0.113), well below the intercept SD of ≈ 0.867. Shift magnitudes co-vary moderately with study-level predictor means in some parameterisations; intercept SDs are nonetheless stable and absolute shifts are small relative to the random-effect SD throughout.

***Table S2.14.*** *Random-effect stability across parameterisations. Intercept SDs are stable (range < 1%).* denotes |r| > 0.3. Absolute shifts are small relative to the intercept SD in all cases.*

| Parameterisation | Intercept SD | Shift range | r vs mean amount | r vs mean patches |
|---|---|---|---|---|
| M1 (baseline) | 0.87 | — | — | — |
| M2 (resFrag + Amount) | 0.86 | [−0.026, +0.031] | -0.07 | 0.613 * |
| M3 (Patches + resAmount) | 0.86 | [−0.084, +0.113] | 0.526 * | 0.377 * |

#### S2.4.3 AICc comparisons

AICc comparisons were conducted for GS25-style versus F26/M1 matched specifications across all 18 variants, and for M1 versus M3 within variants. In the GS25 vs. F26 comparison, the F26 additive specification was favoured in 5 of 18 variants, the GS25 categorical design was favoured in 2 variants, and 11 variants fell within ΔAICc < 2 (equivalent predictive fit). In the within-variant M1 vs. M3 comparison, M1 was preferred in 9 variants and M3 in 1 (median |ΔAICc| = 1.91), consistent with both models representing the data similarly. These comparisons speak to relative predictive fit, not to whether additive coefficients are geometrically interpretable as independent ecological effects.

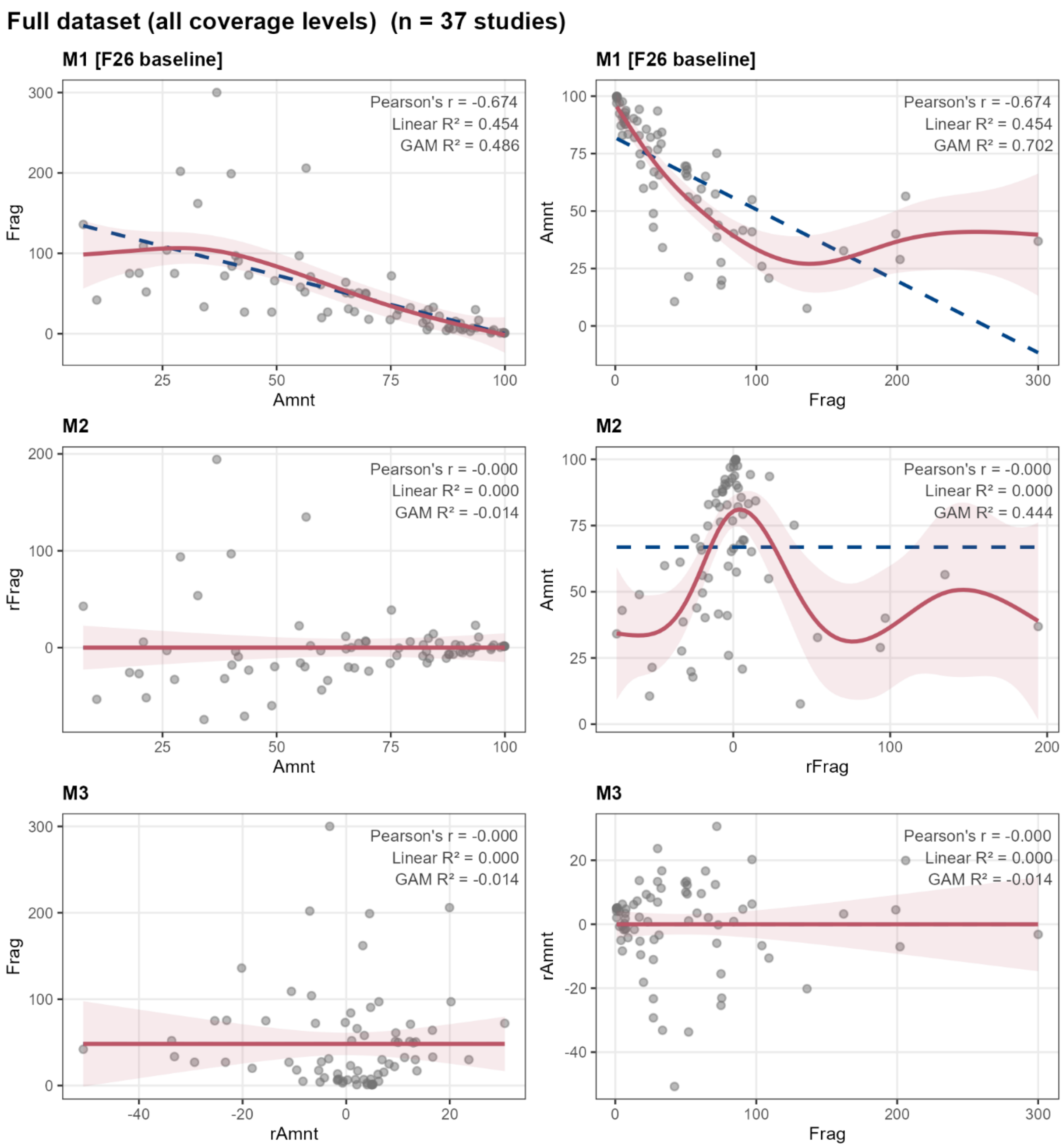


**Figure S2.1.** Bidirectional GAM coupling diagnostics between number of patches (fragmentation) and habitat amount (forest cover) across parameterisations M1–M3, full dataset (n = 74). Left column: fragmentation as response (Frag ~ f(Amount)); right column: amount as response (Amount ~ f(Frag)). Linear fit shown as dashed blue line; GAM fit as solid red line with 95% CI. The nonlinear component is 7.75× larger in the right column (Amount ~ f(Frag), nonlinear $R^2 = 0.248$) than in the left (Frag ~ f(Amount), nonlinear $R^2 = 0.032$), producing a directional asymmetry in which amount is more nonlinearly constrained by fragmentation than the reverse. M2 retains substantial nonlinear coupling in the reverse orientation (right column, residual GAM $R^2 = 0.44$). M3 residualises amount on fragmentation (right column direction) and achieves near-zero residual coupling in both axis orientations (residual GAM $R^2 \approx 0.00$).

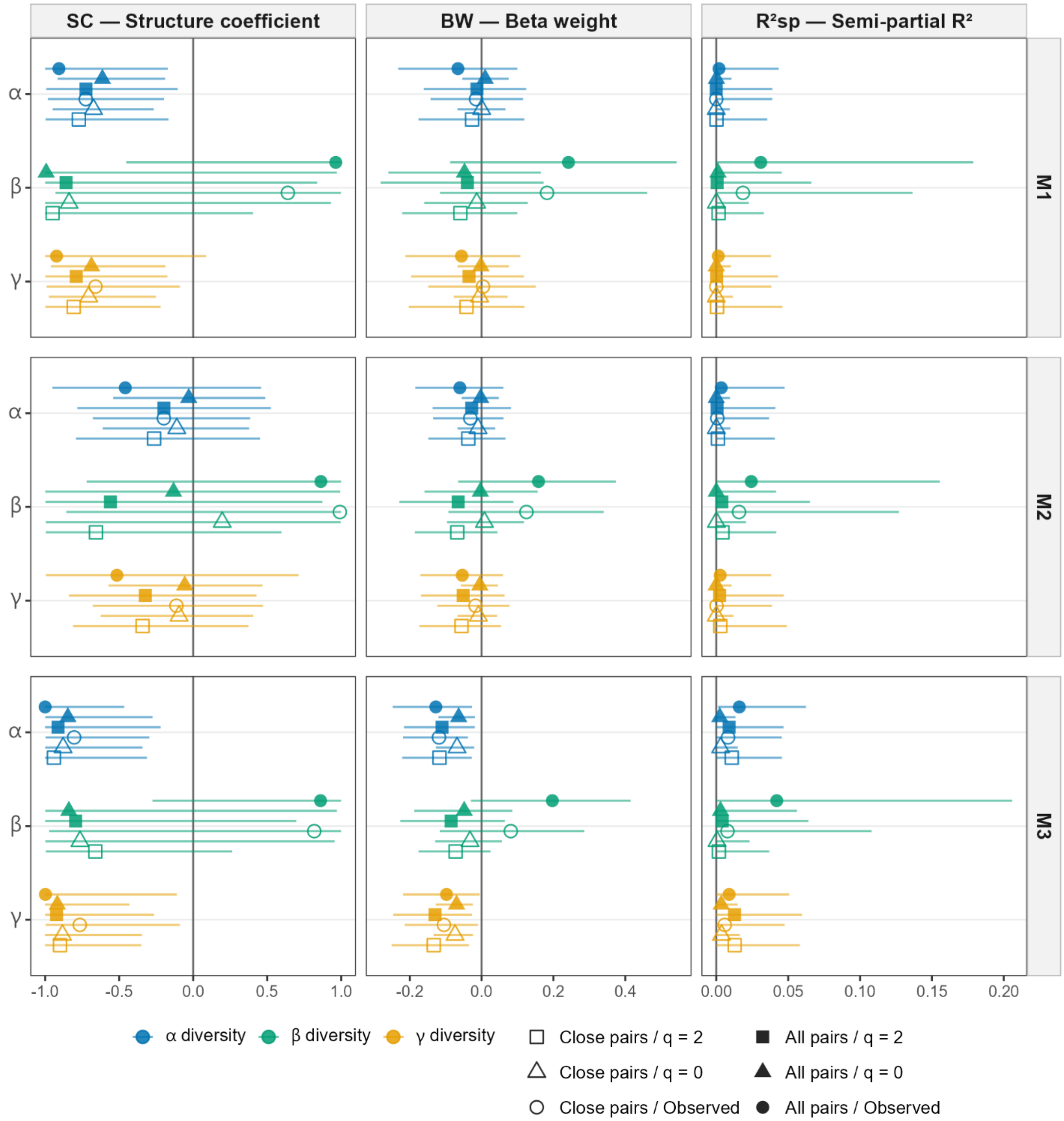


**Figure S2.2.** Variance partitioning for the fragmentation predictor (number of patches) across all 18 analytical variants and parameterisations M1–M3, α, β, and γ diversity. Columns show structure coefficient (SC), standardised beta weight (BW), and semi-partial $R^2$ ($R^2$sp). Point shape encodes pairing design and diversity metric; horizontal lines show 95% bootstrap confidence intervals. Under M1, the fragmentation predictor shows large negative SC across all 18 variants while BW and $R^2$sp remain near zero — the suppressor signature is geometrically invariant, not variant-specific.

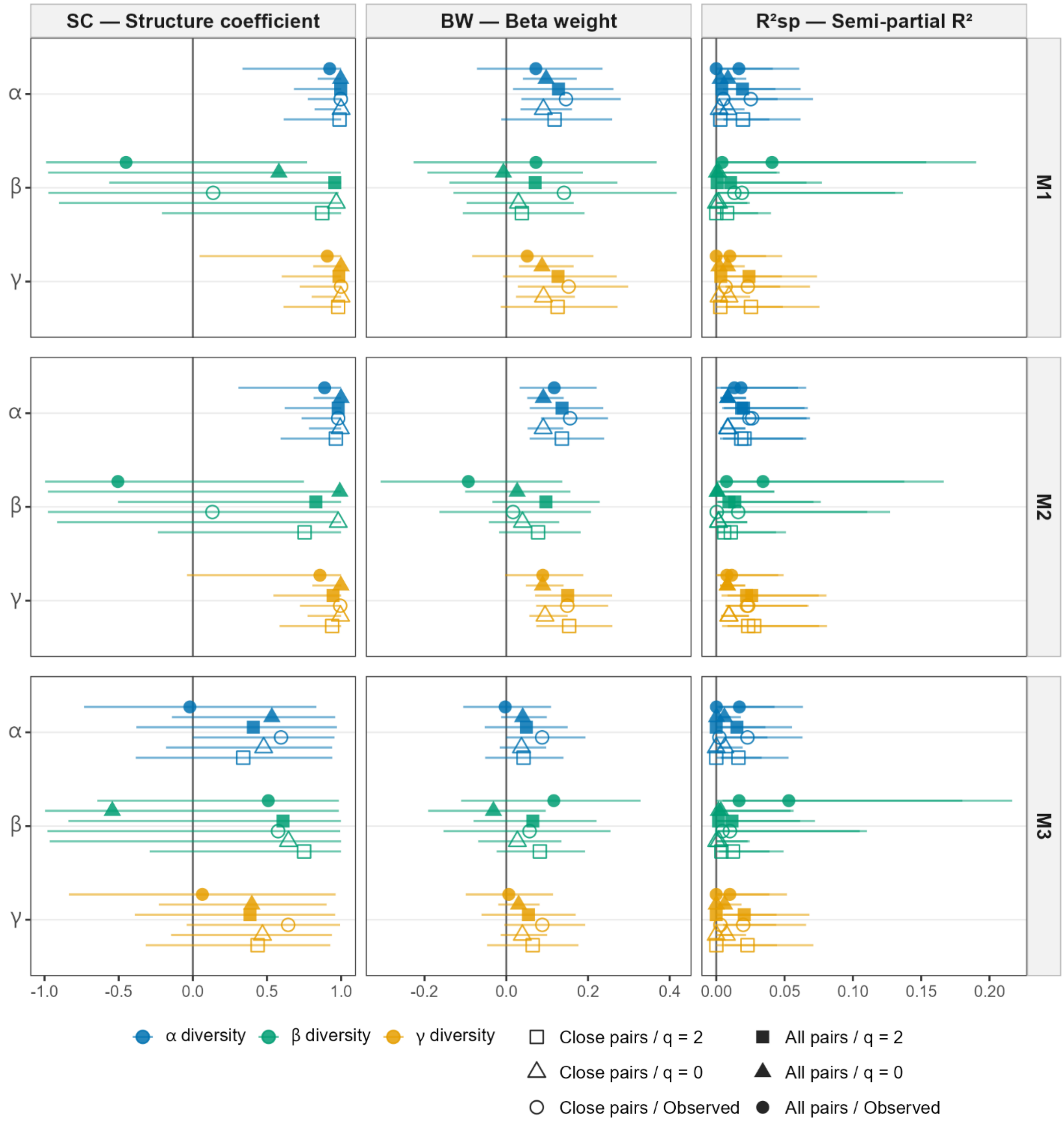


**Figure S2.3.** Variance partitioning for the habitat amount predictor (forest cover) across all 18 analytical variants and parameterisations M1–M3, α, β, and γ diversity. Same format as Figure S2.2. Habitat amount BW is positive across all variants and parameterisations, and decreases under M3 as shared gradient variance is redistributed to the orthogonalised fragmentation term.

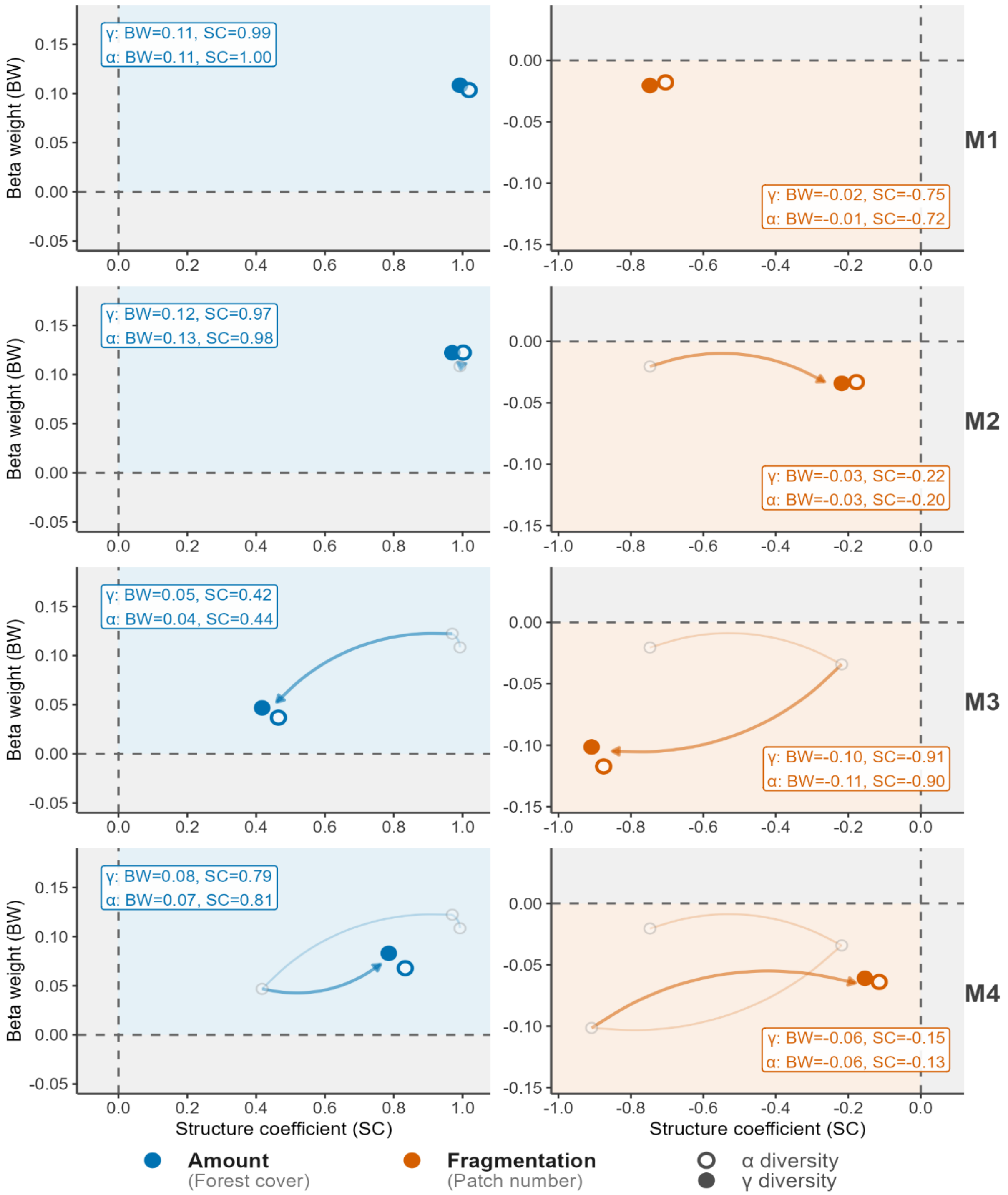


**Figure S2.4.** M4 negative control — SC×BW biplot across parameterisations M1–M4 for α and γ diversity. Fragmentation predictor (right panels) and habitat amount predictor (left panels). Arrows trace the progression M1→M2→M3→M4. Under M1, fragmentation sits in the upper region of the negative-SC half-plane with BW near zero — the cross-over suppressor signature. M3 moves fragmentation into the lower-left quadrant (both SC and BW negative). M4 (negative control) returns near the M1 position, confirming that the M3 shift reflects gradient allocation to the orthogonalised term.

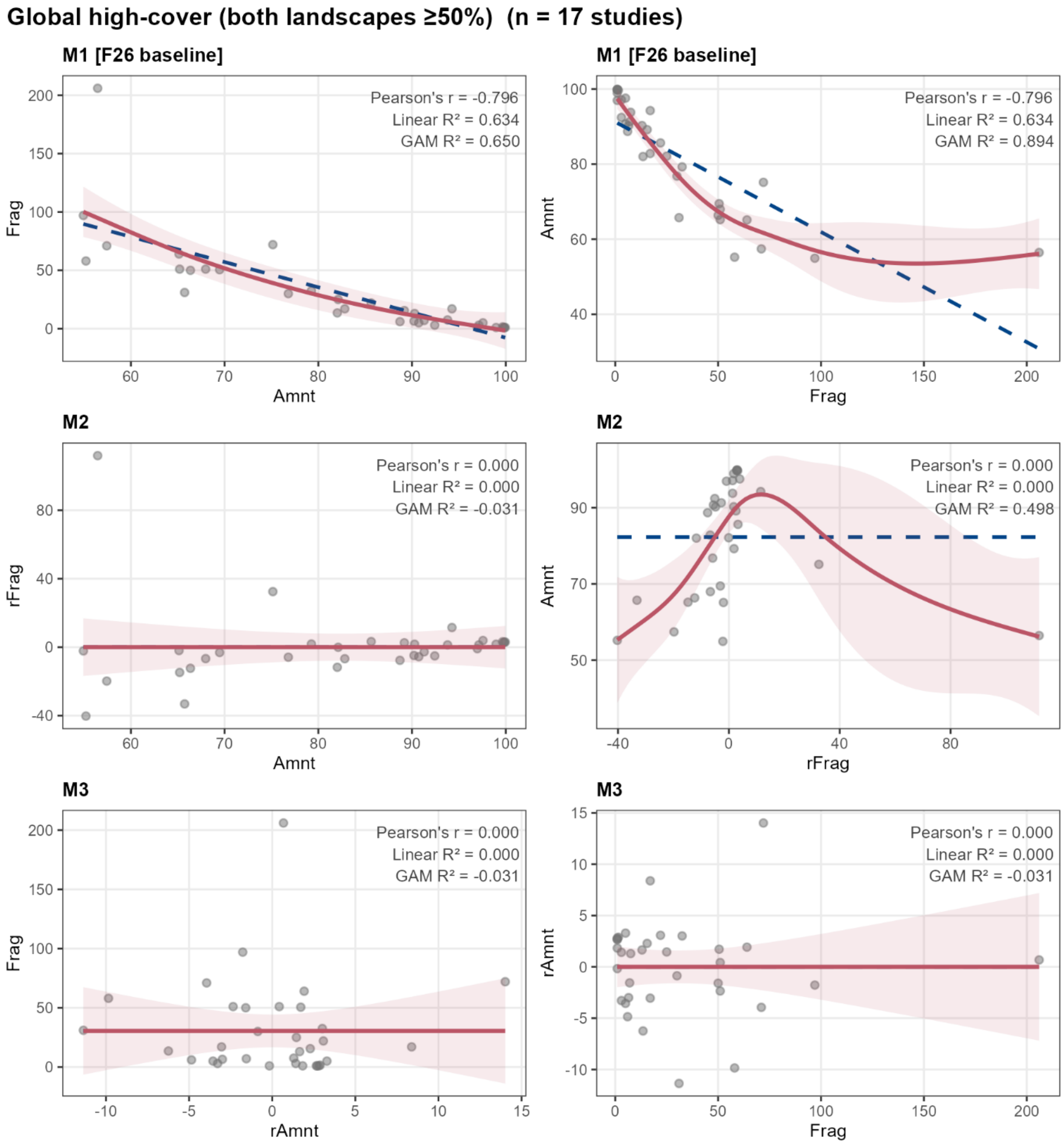


**Figure S2.5.** Bidirectional GAM coupling diagnostics, global ≥50% cover subset (n = 17 studies, both landscapes exceeding 50% habitat cover). Left column: fragmentation as response (Frag ~ f(Amount)); right column: amount as response (Amount ~ f(Frag)). Linear fit shown as dashed blue line; GAM fit as solid red line with 95% CI. The nonlinear component is 16.2× larger in the right column (Amount ~ f(Frag), nonlinear $R^2 = 0.260$) than in the left (Frag ~ f(Amount), nonlinear $R^2 = 0.016$) — a directional asymmetry more than twice as pronounced as in the full dataset (7.75×). M2 retains substantial nonlinear coupling in the reverse orientation (right column, residual GAM $R^2 = 0.498$). M3 residualises amount on fragmentation (right column direction) and achieves near-zero residual coupling in both axis orientations (residual GAM $R^2 \approx 0.00$).

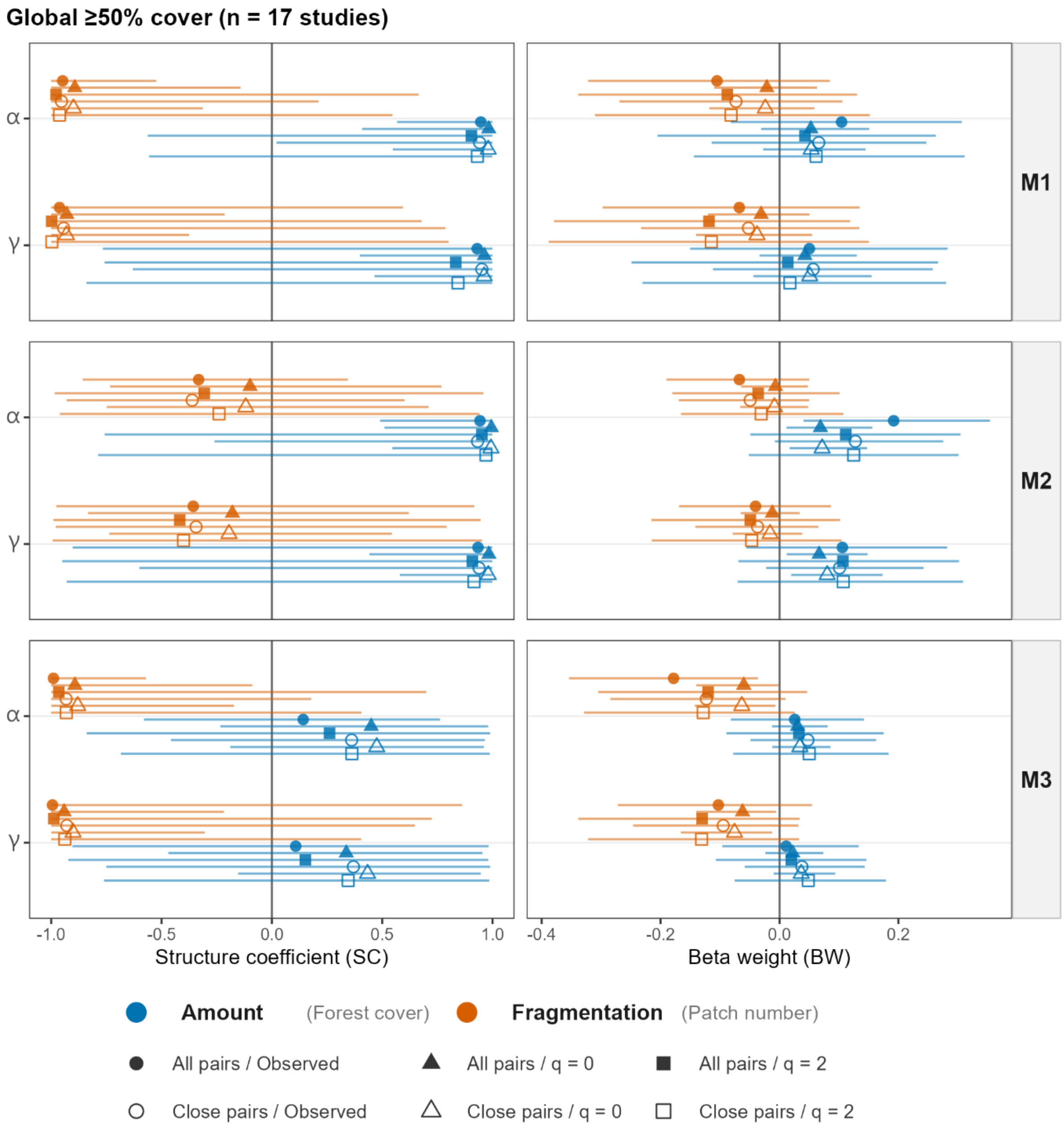


**Figure S2.6.** SC and BW for fragmentation and habitat amount predictors, global ≥50% cover subset (n = 17 studies), parameterisations M1–M3, α and γ diversity. Format as main-text Figure 4. The full suppressor signature (large negative SC, near-zero BW for fragmentation under M1) is preserved within this subset.

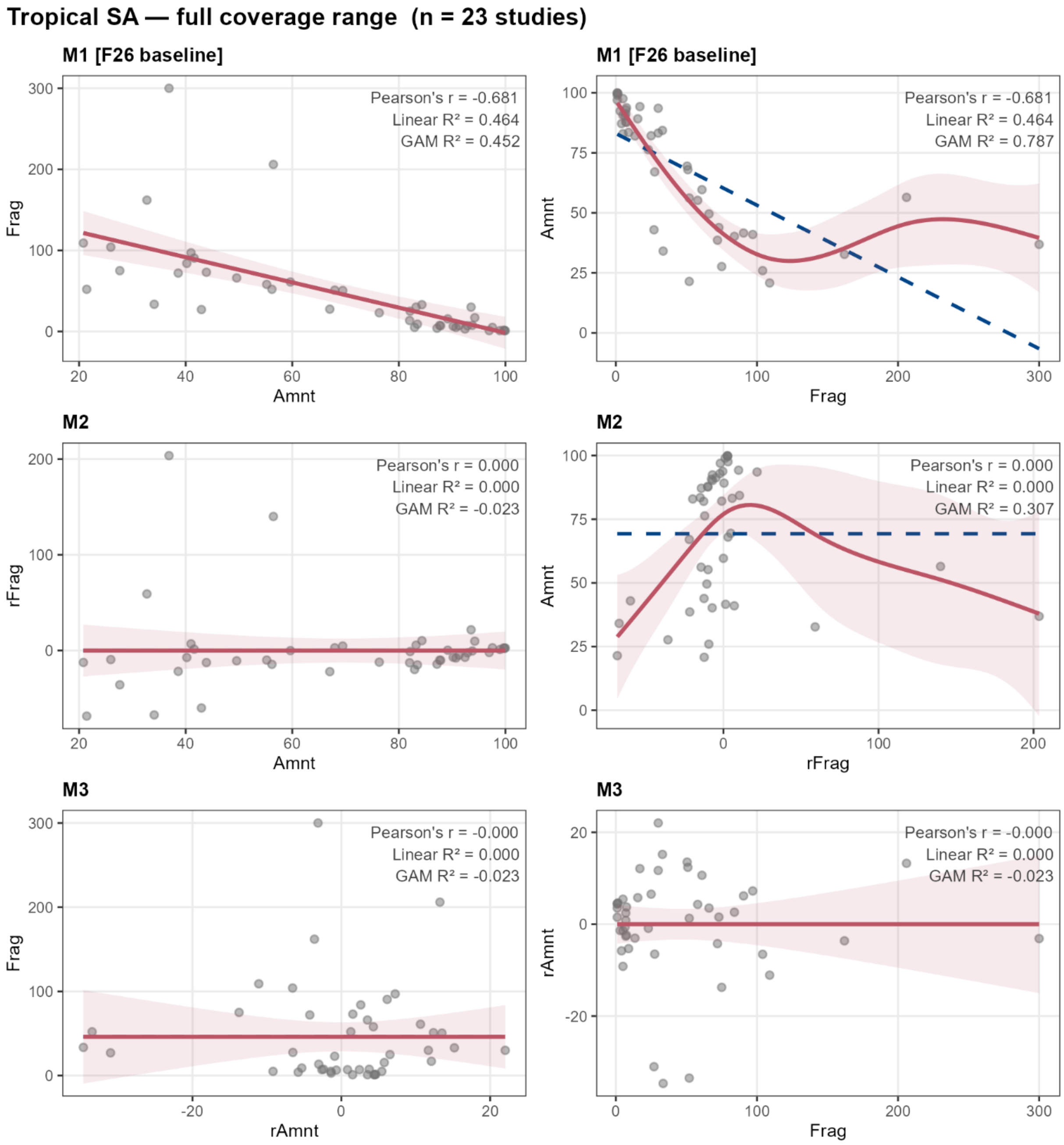


**Figure S2.7.** Bidirectional GAM coupling diagnostics, Tropical SA subset — full coverage range (n = 23 studies). Left column: fragmentation as response (Frag ~ f(Amount)); right column: amount as response (Amount ~ f(Frag)). Linear fit shown as dashed blue line; GAM fit as solid red line with 95% CI. In the left column, the forward relationship (Frag ~ f(Amount)) is approximately linear (GAM $R^2 \approx$ linear $R^2$), while the right column reveals pronounced nonlinearity in the reverse direction (Amount ~ f(Frag), nonlinear $R^2 = 0.323$) — a directional asymmetry consistent with the full dataset pattern. M2 retains substantial nonlinear coupling in the reverse orientation (residual GAM $R^2 = 0.307$). M3 residualises amount on fragmentation and achieves near-zero residual coupling in both axis orientations (residual GAM $R^2 \approx 0.00$).

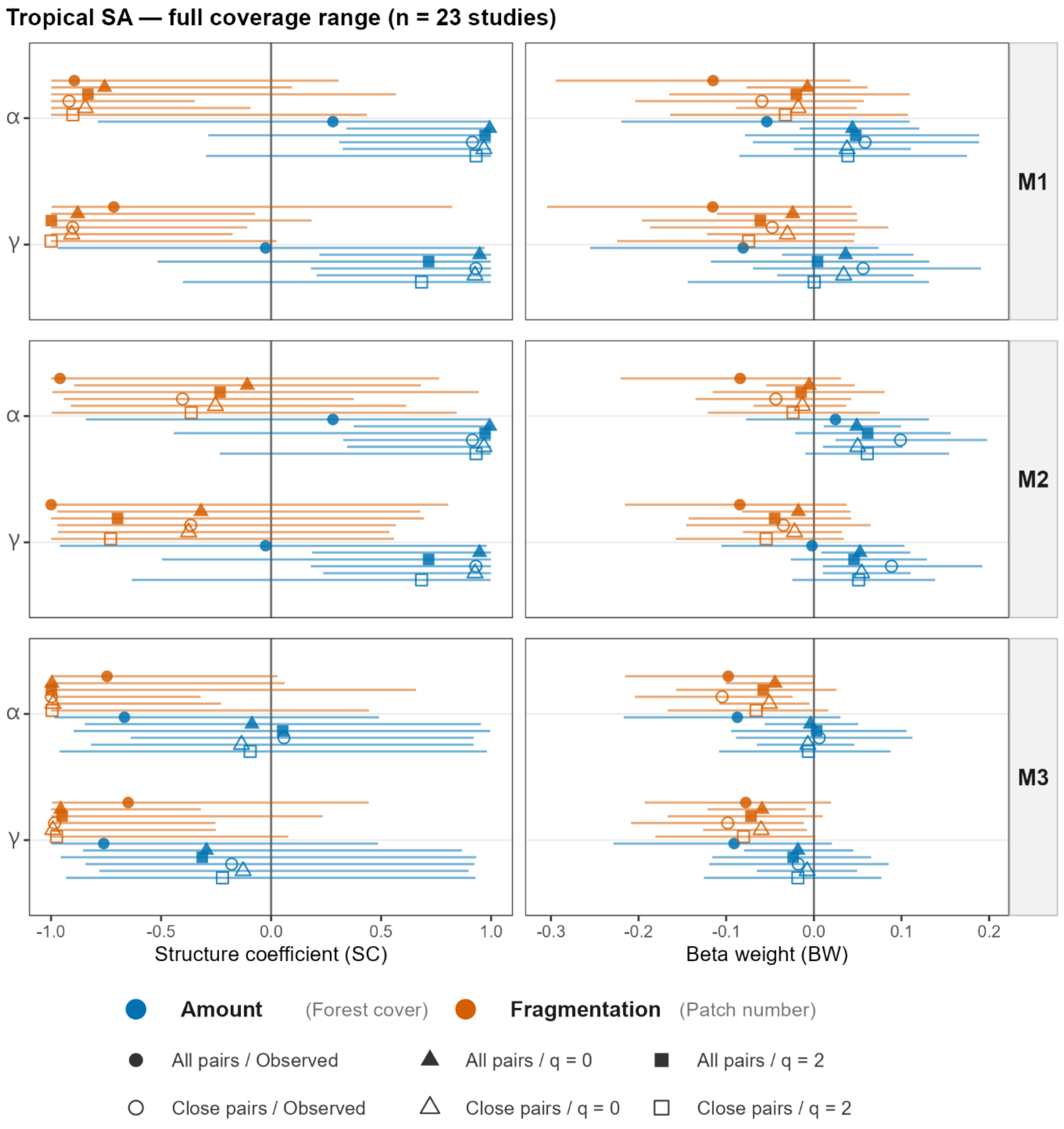


**Figure S2.8.** SC and BW for fragmentation and habitat amount predictors, Tropical SA subset — full coverage range (n = 23 studies), parameterisations M1–M3, α and γ diversity. Format as main-text Figure 4.

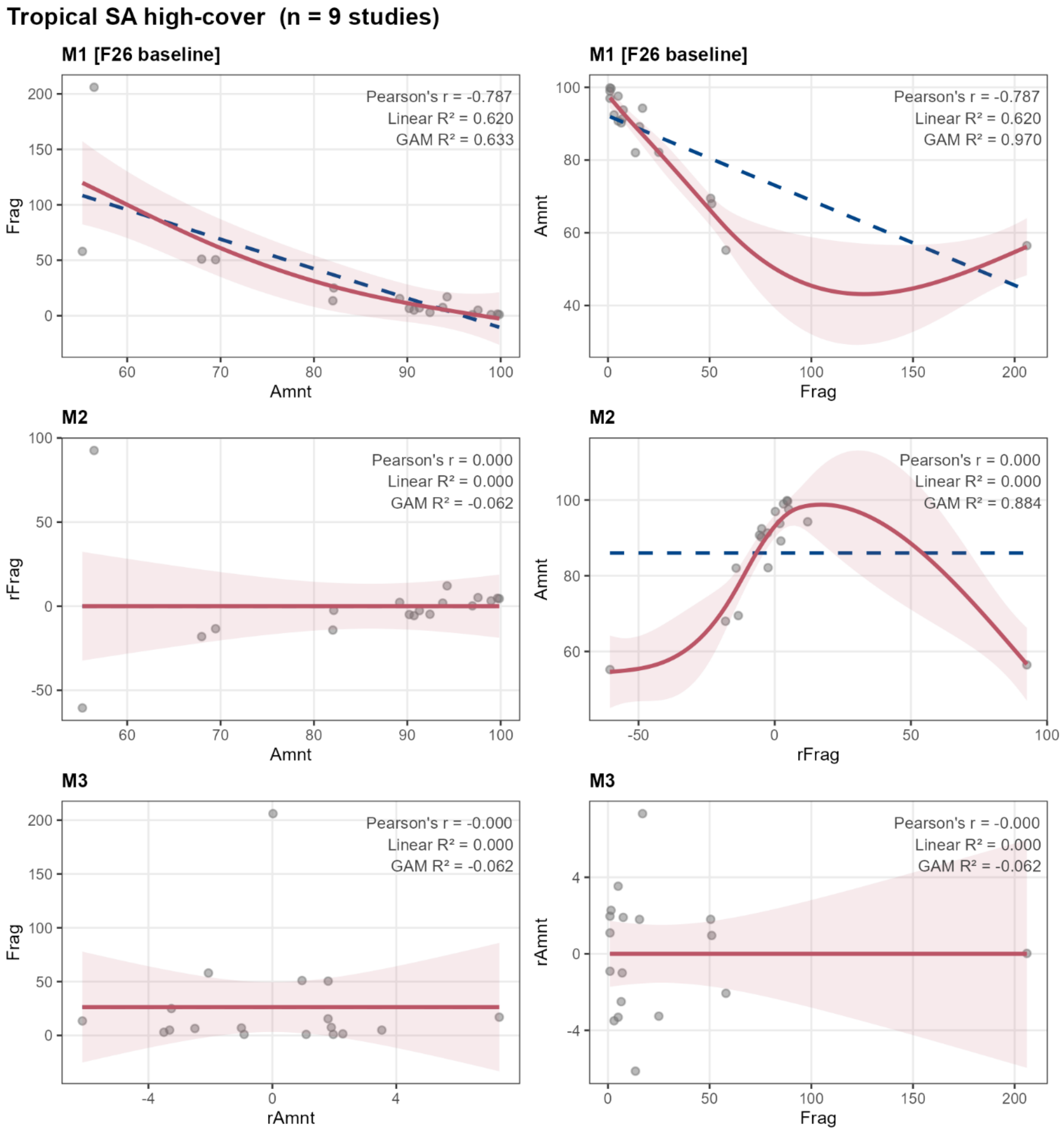


**Figure S2.9.** Bidirectional GAM coupling diagnostics, Tropical SA ≥50% cover subset (n = 9 studies). Same format as Figure S2.1. The M3 panels show residual coupling at the noise floor (Linear $R^2 = 0.000$, GAM $R^2 = -0.062$ in both orientations); the 0.02 orthogonality threshold is met, but the cell is retained as Tier 3 directional indicators only given the small sample (n = 9).
