## Supplementary Material S3 Derivations and Code for "Habitat-loss-driven predictor coupling limits inference about the independent effects of configuration in additive habitat-amount models: implications for the fragmentation debate"

*Juan Andrés Martínez-Lanfranco*

### Table of Contents

**S3.1 Formal derivations underlying the formal inferential argument**
- S3.1.1 The fixed-extent algebraic identity and structural closure of the design space
- S3.1.2 OLS near-cancellation: geometric basis of the near-zero fragmentation coefficient
- S3.1.3 Structure coefficients, variance decomposition, and the suppressor condition

**S3.2 Reproducibility and software details**
- S3.2.1 Software and package versions
- S3.2.2 Key implementation details
- S3.2.3 Pipeline files and code availability

**S3.3 References**

**Tables**

- Table S3.1 Components of the fixed-extent algebraic identity and their inferential implications
- Table S3.2 OLS near-cancellation computation for the M1 α-diversity case
- Table S3.3 Software and R packages used in this analysis

### S3.1 Formal derivations underlying the formal inferential argument

The derivations below clarify the algebraic basis of the suppressor diagnosis reported for this dataset.

#### S3.1.1 The fixed-extent algebraic identity and structural closure of the design-escape route

For a landscape of fixed spatial extent, total habitat area Al, number of habitat patches Np, and mean patch area Āp satisfy the identity:

$$\bar{A}p = Al / Np$$

This relationship is an algebraic consequence of how habitat is partitioned within a bounded landscape, not an empirical regularity that varies across regions or taxa (Fletcher et al. 2023). It implies that any change in habitat amount necessarily alters either patch number or mean patch size or both, making configuration mathematically dependent on amount within any fixed-extent sampling unit.

For patch-number metrics specifically, this dependence has a direction that makes suppressor attenuation structurally predicted rather than contingent: as habitat loss progresses and Al decreases, progressive subdivision means both Al decreases and Āp decreases simultaneously, so Np = Al/Āp can increase even as Al falls — and does so faster than a linear function of Al. Patch-number metrics are therefore not merely correlated with habitat amount; they are functionally coupled through the subdivision process itself, meaning their partial coefficient in additive habitat-amount models is structurally vulnerable to attenuation toward zero, because the subdivision process couples configuration to amount regardless of any independent ecological relationship between configuration and biodiversity.

The standard response to process-coupling concerns is that careful study design could, in principle, select landscapes where amount and configuration vary with sufficiently low correlation to permit additive estimation (Fahrig 2003, 2017). The algebraic identity shows why this escape route is only partial: achieving the orthogonal predictor space that additive habitat-amount models require would demand landscapes where mean patch size varies systematically and independently of both habitat amount and patch number — a condition that characterises landscapes assembled by random spatial processes rather than by progressive land-use conversion. Real habitat-loss landscapes cannot satisfy this condition by design because the coupling is enacted by the land-use process before sampling begins.

***Table S3.1.*** *Components of the fixed-extent algebraic identity and their inferential implications for patch-number metrics in additive models.*

| Symbol | Definition | Interpretation |
|---|---|---|
| Al | Total habitat area | Habitat amount within fixed-extent sampling unit |
| Np | Number of habitat patches | Count of discrete habitat patches within sampling unit |
| Āp = Al / Np | Mean patch area | Algebraic identity: any change in Al or Np must alter Āp; configuration cannot vary independently of amount within fixed extent |

#### S3.1.2 OLS near-cancellation: geometric basis of the near-zero fragmentation coefficient

For a two-predictor regression with standardised predictors under ordinary least squares (OLS), the partial regression coefficient for predictor j is:

$$BW_j = (SC_j - r_{jk} \times SC_k) / (1 - r^2_{jk})$$

where $SC_j$ is the structure coefficient for predictor j (correlation between $X_j$ and the fitted response $\hat{Y}$), $r_{jk}$ is the predictor correlation, and $SC_k$ is the structure coefficient for the co-predictor. When both predictors align strongly with the fitted response but in opposite directions relative to each other (as arises under directed ecological coupling), the numerator $SC_j - r_{jk} \times SC_k$ approaches zero by near-cancellation, even when $|SC_j|$ is large. This is the geometric basis of cross-over suppressor attenuation.

Table S3.2 illustrates the near-cancellation computation for the M1 α-diversity case in the GS25/F26 dataset (Gonçalves-Souza et al. 2025a). The observed values (SC_frag = −0.725, SC_amount = +0.997, r = −0.67) yield a fragmentation numerator of approximately −0.053, producing a near-zero partial coefficient despite the large structure coefficient.

***Table S3.2.*** *OLS near-cancellation computation for the M1 α-diversity case, illustrating the geometric basis of the near-zero fragmentation beta weight.*

| Term | Value / Interpretation |
|---|---|
| SC_frag (M1 α-diversity) | -0.72 |
| SC_amount (M1 α-diversity) | +0.997 |
| r (predictor correlation) | -0.67 |
| Numerator: SC_frag − r × SC_amount | −0.725 − (−0.674 × 0.997) = −0.725 + 0.672 ≈ −0.053 |
| Denominator: 1 − r² | 1 − 0.454 = 0.546 |
| BW_frag ≈ −0.053 / 0.546 | BW_frag (OLS approximation): −0.053 / 0.546 ≈ −0.097 BW_frag (GLMM observed): −0.014 (near-zero; diverges from OLS because random effects are not fully orthogonal to fixed predictors in this dataset) |

The identity holds exactly under OLS. In mixed models (as used in the main analysis), the OLS formula is a pedagogical approximation: the observed GLMM beta weight (−0.014) differs from the OLS approximation (−0.097) because random effects are not fully orthogonal to the fixed predictors in this dataset. The near-cancellation mechanism remains qualitatively identical in both frameworks: numerator suppression through negative predictor correlation drives partial coefficients toward zero regardless of estimation method. The suppressor diagnosis therefore does not depend on the OLS approximation being quantitatively exact.

#### S3.1.3 Structure coefficients, variance decomposition, and the suppressor condition

Structure coefficients (SC) quantify the bivariate correlation between each predictor and the model-predicted response:

$$SCj = cor(Xj, \hat{Y})$$

The square of the structure coefficient equals the inclusive $R^2$ for that predictor:

$$\text{inclusive } R^2_j = SC^2_j$$

Inclusive $R^2$ is expressed relative to $\text{Var}(\hat{Y})$ and measures a predictor's total alignment with the fitted gradient, whether unique or shared with co-predictors. Inclusive $R^2$ decomposes as:

$$SC^2_j = R^2_{j,\text{semi-partial}} + R^2_{j,\text{shared}}$$

The semi-partial $R^2$ (part $R^2$) is the unique variance in the observed response attributable to predictor j after controlling for all other predictors, expressed relative to Var(Y). The shared component is obtained by subtraction. The scale distinction is critical: inclusive $R^2$ is relative to $\text{Var}(\hat{Y})$, while semi-partial $R^2$ is relative to Var(Y). A predictor may show large inclusive $R^2$ while its semi-partial $R^2$ is negligible — the defining pattern of suppressor attenuation.

Cross-over suppressor attenuation is formally identified when:

$$|SC_j| \gg |BW_j| \approx 0 \text{ AND } R^2_{j,\text{semi-partial}} \approx 0 \text{ AND } \text{sign}(SC_j) \neq \text{sign}(BW_j) \text{ for at least some variants}$$

The joint requirement on SC, BW, and semi-partial $R^2$ distinguishes suppressor attenuation from ordinary collinearity, under which both coefficients are proportionally attenuated. Under cross-over suppression, only the suppressed predictor shows near-zero BW and negligible semi-partial $R^2$, while the dominant predictor retains large BW. This mismatch is invisible to standard coefficient inspection, significance testing, and linear collinearity metrics.

### S3.2 Reproducibility and software details

#### S3.2.1 Software and package versions

***Table S3.3.*** *Software and R packages used in this analysis, with versions and primary roles.*

| Package / Function | Version | Use in this analysis |
|---|---|---|
| R | ≥ 4.3.0 | All analyses |
| lme4::lmer | 1.1-35 | Primary model fitting (M1–M4), REML = TRUE for coefficient estimation (Bates et al. 2015) |
| glmmTMB | 1.1.8 | AICc comparisons (ML = TRUE required for valid AIC across fixed-effect structures) (Brooks et al. 2017) |
| mgcv::gam | 1.9-0 | Bidirectional GAM coupling diagnostics; thin-plate splines, k selected by GCV (Wood 2011) |
| partR2 | 0.9.1 | Semi-partial $R^2$ and inclusive $R^2$ via parametric bootstrapping (nboot = 1000) (Stoffel et al. 2022) |
| DHARMa | 0.4.6 | Simulation-based residual diagnostics (Hartig 2022) |
| MuMIn | 1.47.5 | Marginal and conditional $R^2$ (Barton 2024; Nakagawa & Schielzeth 2013) |
| piecewiseSEM | 2.3.0 | Path model fitting (SEM); Fisher's C; path coefficient extraction (Lefcheck 2016) |

| | | |
|---|---|---|
| AICcmodavg | 2.3-3 | AICc computation for model comparisons (Mazerolle 2023) |
| tidyverse | 2.0.0 | Data wrangling throughout pipeline (Wickham et al. 2019) |
| ggplot2 | 3.5.1 | All figures (Wickham 2016) |
| patchwork | 1.2.0 | Multi-panel figure assembly (Pedersen 2024) |
| ggExtra | 0.10.1 | Marginal density panels in Figure S2.1 (Attali & Baker 2023) |

#### S3.2.2 Key implementation details

**Model fitting**

All primary models were fitted using lme4::lmer (Bates et al. 2015) with Gaussian errors and random intercepts by study (1|refshort), estimated by restricted maximum likelihood (REML = TRUE). Continuous fixed-effect predictors were centred and scaled by dividing by two standard deviations prior to fitting (Gelman 2008; Schielzeth 2010), making beta weights directly comparable across predictors. AICc comparisons used glmmTMB (Brooks et al. 2017) with identical formula and random structure, estimated by maximum likelihood (ML = TRUE) as required for valid fixed-effect model comparison.

**Residualisation**

Residualisation used GAM fits (thin-plate splines; basis dimension k selected by GCV; results unchanged for $k = 5$ to $k = 10$). Residual uncoupling was verified post-residualisation using the same bidirectional GAM diagnostic in Step 1, with residual GAM $R^2 < 0.02$ in both orientations as the criterion. M3 satisfied this criterion; M2 did not (substantial nonlinear coupling retained despite zeroing linear r).

**Semi-partial $R^2$ and inclusive $R^2$**

Semi-partial $R^2$ was extracted via partR2 (Stoffel et al. 2022; nboot = 1000; CI = 0.95). Note that partR2 operates on the response scale (relative to Var(Y)), while the inclusive $R^2$ (= $SC^2$) is expressed relative to $\mathrm{Var}(\hat{Y})$. The shared variance component is obtained as $SC^2$ − semi-partial $R^2$ after scaling to a common reference, using the decomposition from Ray-Mukherjee et al. (2014). Marginal and conditional $R^2$ were computed using MuMIn (Barton 2024) following Nakagawa and Schielzeth (2013).

**Bidirectional GAM diagnostics**

GAMs were fitted using mgcv::gam (Wood 2011) with thin-plate regression splines (bs = 'tp'). Basis dimension k was selected by generalised cross-validation; sensitivity to k was confirmed by repeating diagnostics for k = 5, 7, and 10, with unchanged results. The GAM $R^2$ values reported are the adjusted $R^2$ returned by mgcv summary; the linear $R^2$ in each orientation is the squared Pearson correlation, identical in both directions.

#### S3.2.3 Pipeline files and code availability

R code implementing all diagnostic workflows reported here — including bidirectional GAM coupling assessment, structure-coefficient extraction, residualisation sequence across M1–M4, orthogonality verification, and semi-partial $R^2$ via partR2 — is available at [Code repository —

URL to be provided upon acceptance to maintain anonymity during review]. The key pipeline script files are:

**00_master.R** — single entry point; sources 01–05 in sequence with configurable REFIT and NBOOT settings02_pipeline_results.R — SC, BW, semi-partial $R^2$ compilation across 18 variants

**01_fit.R** — model fitting (M1–M4) across all analytical variants (partR2, glmmTMB, lme4); saves fit_objects.rds

**02_diagnostics.R** — post-fit diagnostics: AICc, DHARMa residuals, logistic decomposition, SEM, marginal $R^2$

**03_extract.R** — extracts all reported numbers into key_scalars.csv and block_estimates.csv

**04_figures.R** — generates all manuscript and supplement figures from CSVs (but main text Figures 1–2 and 5).

**05_tables.R** — generates all supplement tables from CSVs

All analyses were conducted in R v. ≥ 4.3.0 (R Core Team 2024). The dataset was originally compiled and analyzed by Gonçalves-Souza et al. (2025a,b) and is freely available at Zenodo (https://zenodo.org/records/12206838). The same dataset was reanalysed by Fahrig et al. (2026), with the code available from https://osf.io/hu3xw. Code for all diagnostic workflows and analysis of this study is available at https://github.com/jacoloml/Reanalyses_GS25_F26 (archived at https://doi.org/10.5281/zenodo.19546467). Reporting of software versions, diagnostic workflows, and executable analysis pipelines follows current recommendations for valid and reproducible statistical ecology (Popovic et al. 2024).
